# An Open-Source, Excitation-Resolved UV-NIR Imaging Platform for Bioluminescence, Luminescence-Decay, and Chemical Discrimination Measurements

**DOI:** 10.64898/2026.07.30.741847

**Authors:** John M. Branning, Cameron L. Lyman, Isabella A. Hensley, Elise Jakel, Tyler Weiskopf, Gabrielle Link, Natalie J. Serkova, Adam L. Green, Kevin J. Cash

## Abstract

Multispectral imaging is a cornerstone of chemical imaging, with bioluminescence, fluorescence, and phosphorescence imaging underpin much of preclinical and analytical chemical measurements. However, conventional implementations are typically proprietary, costly, and resolve spectral content from the reflectance of a broadband source, without control over the excitation spectrum. This architecture cannot isolate excitation-dependent photophysical processes. Spectral discrimination through sequentially resolved narrowband excitation, detected on a single broadband-sensitive camera, instead enables excitation-dependent fluorescence, phosphorescence, and reflectance measurements not available to such broadband approaches. We describe AURORA-MSI, an open-source multispectral platform that inverts this arrangement with fourteen narrowband LEDs spanning 367-940 nm, a 120-element annular illumination ring with eight independently addressable azimuthal sectors, and a thermoelectrically cooled monochrome CMOS camera. Radiometric calibration, spatial uniformity mapping, and camera noise characterization establish the quantitative measurement foundation. In bioluminescence imaging, AURORA-MSI localized sources in a calibrated tissue-mimicking mouse phantom comparably to a commercial Revvity IVIS Spectrum, detected luciferase-expressing HSJD-GBM1-001 glioblastoma cells across a dilution series, with reduced replicate consistency at the lowest densities, and mapped substrate-free fungal-pathway emission in an intact bioluminescent *Petunia hybrida*, none requiring photon-counting instrumentation. Time-resolved phosphorescence decay imaging of six inorganic phosphors over 33 minutes found tri-exponential models adequate at early times, while distributed-lifetime models were preferred at extended timescales. Multispectral image-texture features extracted across all fourteen excitation bands discriminated ten pharmaceutical and household powders spanning distinct chemical compositions and, in two cases, distinct formulations of the same compound. Generality beyond these regimes was established through dual-excitation fluorescence fingerprinting of fifteen mineral specimens and wavelength-selective plant-tissue imaging exploiting UV and NIR penetration-depth differences. Excitation-side spectral encoding enables photophysical measurements that detection-side systems with uncontrolled broadband illumination cannot isolate, and the cooled-camera architecture supports weak-signal modalities without photon-counting instrumentation.

## Introduction

Multispectral imaging combines spatial resolution with chemical specificity by acquiring a spectral signature at every pixel, and has become a core measurement tool in chemical and biomedical imaging.^1,2^ It supports intraoperative tissue classification and tumor-margin assessment, and underpins longitudinal, noninvasive monitoring of reporter-gene expression and tumor burden using bioluminescent and fluorescent probes in small-animal models.^3–5^ The same measurement principle operates across geological and remote-sensing surveys, precision agriculture, food-quality inspection, and forensic analysis, where reflectance signatures identify materials without contact or sample preparation.^6–8^

Most commercial multispectral imagers resolve spectral content on the detection side: broadband or white-light illumination floods the sample, and a dispersive element, liquid-crystal tunable filter, acousto-optic device, or snapshot mosaic sensor separates reflected or emitted light into narrow spectral channels.^9–13^ The resulting data cube encodes the spectral reflectance or emission profile of each spatial pixel, providing detailed spectral information about sample composition but no control over the excitation conditions. This approach is optimal for passive remote sensing and material identification based on reflectance spectra but cannot, by design, isolate excitation-dependent photophysical phenomena such as wavelength-specific fluorescence excitation, phosphorescence with controlled excitation dose, or directional scattering that depends on the angle of incidence.

An alternative architecture reverses this paradigm through spectrally resolved excitation: the sample is illuminated sequentially with narrowband sources while emission is integrated on a single broadband-sensitive detector.^14^ This excitation-side configuration decouples spectral content from detection bandwidth and allows one sensitive monochrome camera to serve as the integrating element across all excitation wavelengths. The approach is well established in fluorescence excitation spectroscopy and structured-illumination microscopy^15,16^, and has been extended to low-cost multi-LED systems for microscopy, food inspection, environmental monitoring, and whole-animal fluorescence imaging.^17–23^ Open-source instrumentation has likewise emerged in adjacent domains^24–27^: the OpenHSI project released a push broom spectrometer for remote sensing^28^, and open implementations of the IVIS architecture have been described for small-animal fluorescence imaging.^29^ Open acquisition and analysis platforms have likewise been reported for multiplexed bioluminescence microscopy, pairing conventional detection hardware with freely available control and analysis software.^30^ However, no open platform reported to date, to our knowledge, combines broadband UV-NIR excitation-side spectroscopy, azimuthally resolved angular illumination, and cooled-camera detection within a single modular instrument, or applies such an architecture systematically to wide-field, macroscale imaging in an open, reproducible form.

Here, we introduce the Active UV-NIR Radiance and Optical Reflectance Analyzer Multispectral Imager (AURORAMSI), a multispectral imaging platform designed to address these limitations. We describe its design, performance characterization, and application across five complementary imaging demonstrations spanning mineralogy, luminescence decay, bioluminescence, pharmaceutical powder classification, and plant-tissue stress imaging.

## Results and Discussion

### Platform Design and Engineering Rationale

The AURORA-MSI design (Figure S1) was guided by six engineering principles: (i) spectral coverage from the UV to the NIR; (ii) excitation spectral encoding using wavelength-selective illumination rather than broadband illumination with spectrally resolved detection; (iii) independent sector-level angular illumination control for directional reflectance and photometric-stereo acquisition; (iv) a light-tight enclosure excluding ambient illumination, necessary for excitation-encoded weak-signal imaging rather than ambient-tolerant broadband reflectance; (v) a fully 3D-printed, modular optomechanical assembly requiring no custom machining; and (vi) low-noise imaging capability for weak signal applications including bioluminescence and long-duration phosphorescence decay. These principles follow an emerging design practice in open scientific instrumentation, in which 3D-printed modular optomechanics and documented hardware-software stacks allow multiple imaging modalities to be reconfigured within a single platform by non-specialist users.^31^

### Instrumentation

The mechanical housing is entirely 3D-printed from either PLA or PETG, with modular mounting interfaces and is designed as a light-tight enclosure that excludes ambient illumination during acquisition. Detection is performed by a commercial cooled monochrome CMOS camera (ZWO ASI533MM Pro), thermoelectrically cooled to ∼35 °C below ambient. The monochrome architecture avoids Bayer-pattern interpolation artifacts and allows improved quantum efficiency across 300-1000 nm at every pixel. An interchangeable emission filter holder is installed between lens and sample, enabling rapid band switching without disturbing sample geometry (Figure S1E). The annular illumination ring (65 mm diameter) carries 120 LEDs spanning 14 wavelengths (367-940 nm) distributed symmetrically across eight 45° azimuthal sectors; each sector and each wavelength channel is independently addressable with continuous PWM intensity control (Figures S1-S2). A microcontroller board and Python-based GUI provide integrated control of illumination and camera subsystems, including time-lapse and parameter-sweep acquisition.

### System Calibration

Radiometric calibration established the relationship between drive settings and delivered sample-plane irradiance across all excitation bands (367-940 nm; Figures S3-S8, Supporting Information). Delivered optical output power increased monotonically with PWM drive setting (R^2^ = 0.974 ± 0.081) and sector contributions were nearly equal across visible wavelengths, supporting azimuthally symmetric illumination when all sectors are enabled. Measured signal levels spanned approximately two orders of magnitude across the excitation range, peaking near 568 nm and declining toward the UV (367 nm) and deep-NIR (850, 940 nm) extremes, consistent with the camera sensor’s quantum efficiency roll-off outside the visible range (Figures S3-S4). Signal behavior with exposure was most linear and predictable in the mid-visible bands, becoming less consistent at the spectral extremes. All quantitative imaging in this work was performed within the confirmed linear, unsaturated operating window for each wavelength. Detector noise, spatial nonuniformity, and calibration drift are recognized as dominant error sources limiting quantitative reliability in camera-based luminescence and chemical imaging, and the and comparable characterization of LED beam profile, resolution, and penetration depth has been established as a prerequisite for quantitative measurement in LED-illuminated biomedical imaging systems.^32–34^

### Example Applications

We evaluated AURORA-MSI through three core demonstrations testing the quantitative claims of the architecture, followed by two further demonstrations establishing generality across specimen classes. Bioluminescence imaging of a calibrated tissue-mimicking phantom, a luciferase-expressing cell dilution series, and an intact self-luminous organism tested detection of ultra-weak self-luminous signals against a commercial reference instrument. Time-resolved phosphorescence decay imaging tested extended-duration weak-signal acquisition and decay-kinetics model selection. Multispectral texture classification of pharmaceutical and household powders tested excitation-resolved discrimination of chemically distinct but visually indistinguishable materials. Two additional demonstrations, dual-excitation fluorescence fingerprinting of mineral specimens and wavelength-selective plant-tissue imaging, extend the platform to inorganic solids and living tissue, establishing that excitation-side encoding generalizes across specimen classes with markedly different optical properties.

### Mineral Fluorescence Fingerprinting

Fifteen mineralogical specimens were imaged under 367 and 405 nm excitation with a series of long-pass emission filter cutoffs. Minerals were treated as stable optical samples rather than as geological specimens^35^. The fingerprint matrix (Figure 1A), with rows ordered by hierarchical clustering on cosine distance, shows coherent blocks of minerals sharing similar spectral shape cluster together (Chalcedony, Opalite, Selenite, Fluorite, Calcite-A/B), while minerals with red-shifted or broadband emission (Hackmanite, Scapolite, Tremolite) form distinct branches.

**Figure 1.**
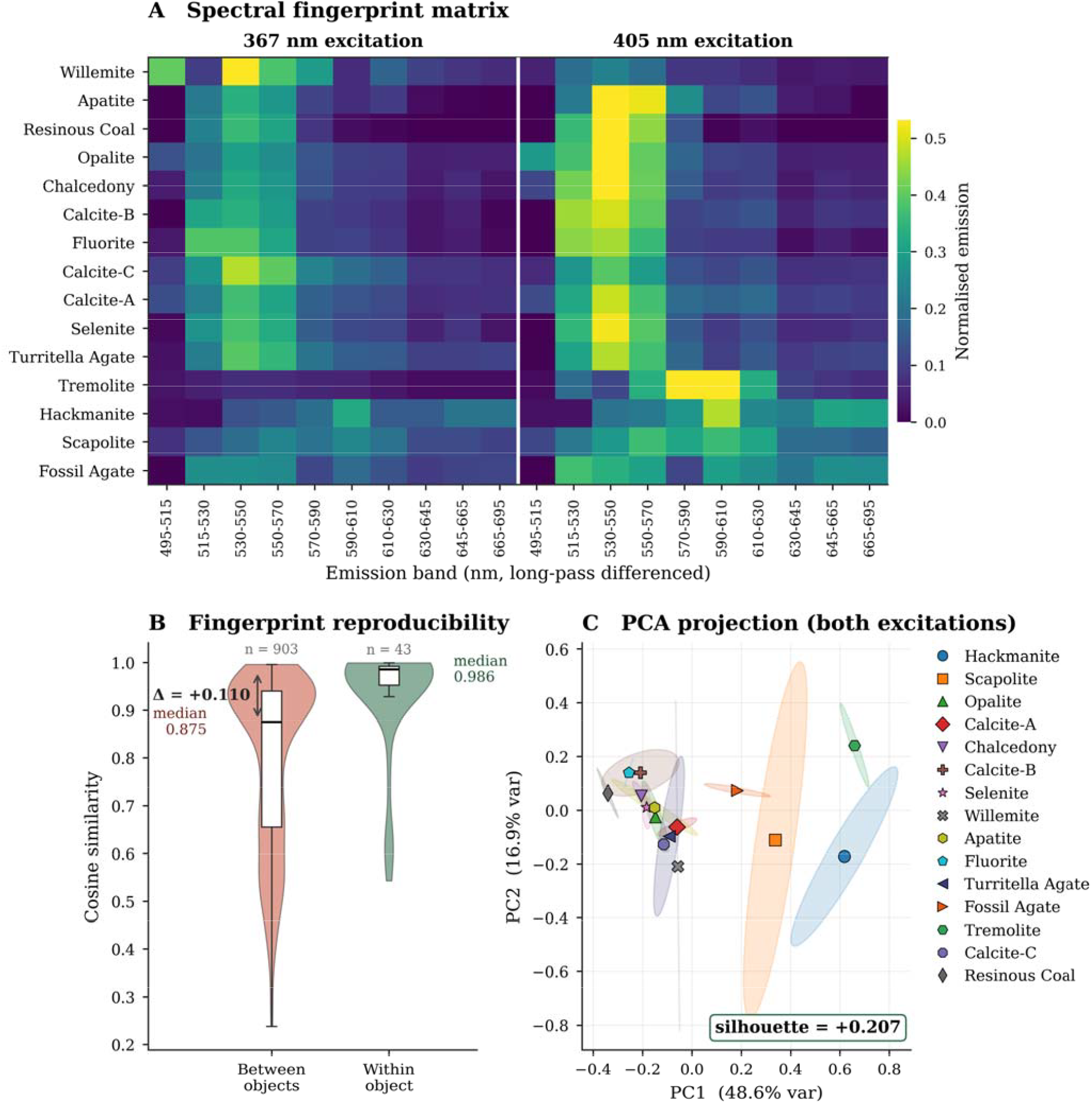
Discrete-wavelength excitation with filtered monochrome detection produces reproducible, separable optical fingerprints **for** 15 mineral samples. (A) Spectral fingerprint matrix for 15 mineral specimens. Rows represent mineral samples ordered by hierarchical clustering, and columns represent band-differenced emission features acquired under 367 nm (left) and 405 nm (right) excitation. **Co**lor intensity indicates normalized emission. (B) Pairwise cosine similarity distributions comparing within-sample and between-sample **RO**Is, demonstrating higher fingerprint reproducibility within individual minerals than between different minerals. (C) Principal component analysis of the combined excitation features. Each point represents an ROI colored by mineral type; ellipses indicate within-class variability. The projection indicates partial separation of mineral classes based on excitation-dependent fluorescence fingerprints.

Per-sample feature vectors exhibited high within-object cosine similarity and substantially lower between-object similarity (Figure 1B), confirming that discrete-wavelength excitation with filtered detection produces reproducible and discriminative optical signatures. The absolute silhouette score^36^ represents a three-to-fivefold improvement over a single-excitation condition (Figure 1C; Figure S9) demonstrating that excitation diversity encodes spectral information not accessible from a single wavelength. We also characterized individual emission profiles, excitation-dependent spectral crossovers, and the photophysical origins of the observed emission for selected minerals (Figures S10-S12). These results indicate that dual-excitation imaging improves discrimination of several mineral specimens compared with the single-excitation conditions examined here.

With 12 of 13 minerals represented by a single sample and three ROIs, the within-object cosine similarity we report is a within-sample reproducibility metric, not a within-mineral one. The three geographically distinct calcite specimens provide a preliminary indication that the platform is sensitive to trace-activator variation,^35^ but generalization to mineral-level identification requires a future replication study with multiple specimens per mineral class.

These results illustrate a core design principle of the platform in which a small number of discrete excitation wavelengths combined with filtered detection is sufficient to produce compact, discriminative fluorescence fingerprints. The silhouette improvement under dual excitation (Figure 1C, Figure S9) suggests that combining excitation diversity with filtered detection allows improved discrimination under the acquisition conditions investigated.

### Time-Resolved Phosphorescence Decay Imaging

Persistent luminescence materials are increasingly employed as reporters in biomedical imaging, security marking, and sensing applications where the decay lifetime, rather than steady-state emission intensity, carries the analytical signal.^37–39^ Time-resolved phosphorescence decay imaging was therefore used to assess whether the platform can distinguish phosphor materials on the basis of their decay kinetics and to evaluate the fidelity of lifetime parameter recovery across spatially resolved regions of interest.

Six commercial photoluminescent pigment powders were evaluated: five color-coded strontium aluminate (SrAl□O□:Eu^2+^,Dy^3+^) formulations (aqua, blue, green, orange, and purple) and one yttrium oxide (Y□O□:Eu^3+^) formulation (Red). These color names denote product designations rather than excitation or emission wavelengths and are used throughout this section to identify individual samples. All six phosphors were well-described by tri-exponential fits within the first 300-second acquisition window (Figure 2, Figure S13), spanned a 6.5-fold range of amplitude-weighted mean lifetimes, and were clearly resolved in lifetime-amplitude space (Figure S14). The tri-exponential fits are consistent with first-order detrapping kinetics when retrapping is negligible at early times.^40^ However, at longer timescales retrapping becomes increasingly significant, causing persistent luminescence decay to deviate from exponential behavior.^41–43^

**Figure 2.**
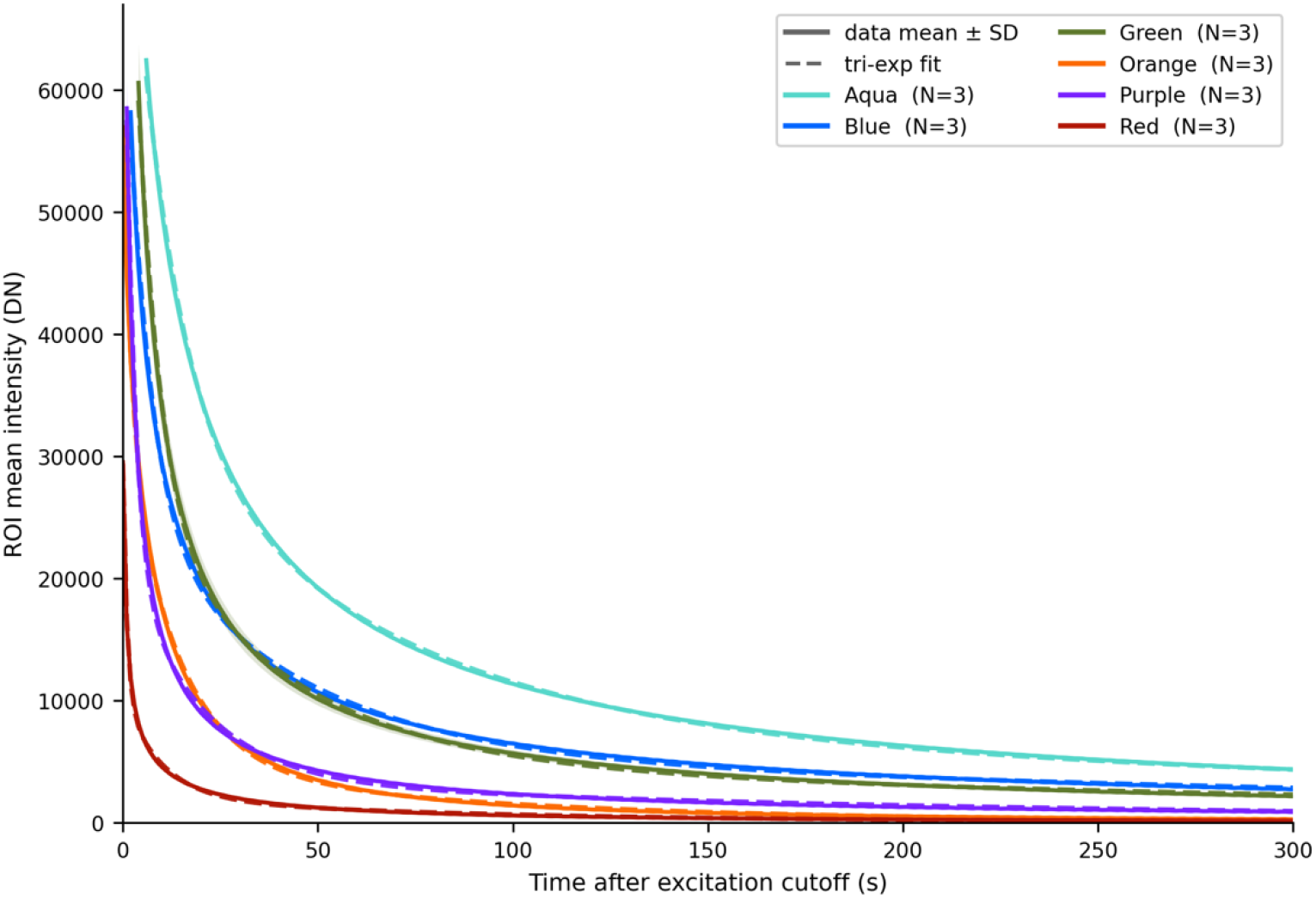
Phosphorescence decay curves for six phosphor samples. Solid lines show mean phosphor intensity ± SD (shaded band, N = 3 per phosphor); dashed lines show tri-exponential fits . The measured decay profiles exhibit nonlinear behavior on a linear intensity scale and are well described by the fitted tri-exponential model.

We therefore evaluated a set of distributed-lifetime and power-law models, stretched-exponential (Kohlrausch– Williams–Watts), power-law, log-normal, gamma-distribution, and Tsallis-q, alongside the discrete multi-exponential models. These distributed models, which do not assume discrete trap populations or negligible retrapping, are better suited to describing the continuous trap-depth distributions that govern persistent luminescence at extended timescales. The deviation from an exponential decay model is apparent on the semi-log plot of the extended lifetime measurement window (Figure S15). Coefficient of determination (R^2^) and Akaike Information Criterion (AIC) model comparison (Table S2-S3) indicate that a power-law model or continuous distribution model outperformed the tri-exponential model (Figure S16-S18).^44,45^

The dominance of distributed-lifetime models for the SrAl□O□:Eu^2+^,Dy^3+^samples is consistent with the broad, continuous trap-depth distributions that characterize this host^46^, arising from the complex crystal field environment of Eu^2+^and the multiple Dy^3+^trap levels introduced by codoping.^47,48^ The faster and more complete decay of Y□O□:Eu^3+^within the measurement window reflects the fundamentally different emission mechanism, □D□→□F□electric dipole transitions of Eu^3+^rather than the 4f□5d^1^→4f□ broadband transitions of Eu^2+^, and the absence of the deep Dy^3+^trap reservoir that sustains long-duration afterglow in the strontium aluminate system.^49,50^

Orange and red phosphors fell below 15 dB SNR at ∼300 s (Figure S19), setting the practical measurement limit for these two phosphors in this measurement configuration. This earlier SNR floor reflects the combined effect of excitation/detection efficiency and photon budget at these wavelengths and should not, on its own, be read as evidence that orange and red have intrinsically shorter afterglow than the other four phosphors. For the aqua, blue, and green phosphors, which maintained SNR > 15 dB throughout 1,980 s, the full decay dynamics are captured within the measurement window, providing adequate photon statistics for quantitative lifetime extraction without photon-counting instrumentation.^51,52^ For inorganic afterglow phosphors with τ_1_ > 1 s, as observed here, camera-based acquisition offers practical advantages since it requires no ultrafast pulsed excitation source, no synchronization electronics, and no photon-counting detector, reducing instrument complexity by orders of magnitude.^53^ Camera-based approaches have likewise been developed to recover lifetime-proportional signals from conventional sensors without photon-counting hardware, extending the accessible regime for luminescence-lifetime measurement.^54^

### Bioluminescence Imaging

Bioluminescence imaging is a cornerstone modality in preclinical biomedical research, enabling longitudinal, noninvasive monitoring of tumor burden, reporter gene expression, and infection in small animals.^5,55,56^ Sustained development of engineered luciferases and synthetic luciferin analogues has since extended the modality toward multiplexed and spectrally resolved reporter detection, placing corresponding demands on detector sensitivity and acquisition flexibility.^30,57^ To assess the platform’s sensitivity for bioluminescence imaging across complementary biological and physical sources, we evaluated three targets: a calibrated tissue-mimicking phantom, a self-luminescent intact organism (genetically engineered bioluminescent Petunia hybrida), and a dilution series of luciferase-expressing mammalian cells in a 96-well plate format.

To benchmark sensitivity against a calibrated, instrument-independent reference, we imaged a commercial Revvity XPM-2 calibrated mouse phantom and compared results to images acquired on a Revvity IVIS Spectrum system^3,5^. The XPM-2 phantom contains two calibrated luminescent sources embedded in a tissue-mimicking material shaped as a supine mouse in dorsal orientation, providing a standardized target for cross-instrument comparison. Figure 3 shows results with the phantom’s “Mode A” source active (single-source illumination, thoracic midline) and Figure S20 shows the results with “Mode B”. Panel A presents the IVIS Spectrum reference image, while panels B-D are AURORA-MSI images acquired with 1 s, 10 s, and 30 s exposure times, respectively.

**Figure 3.**
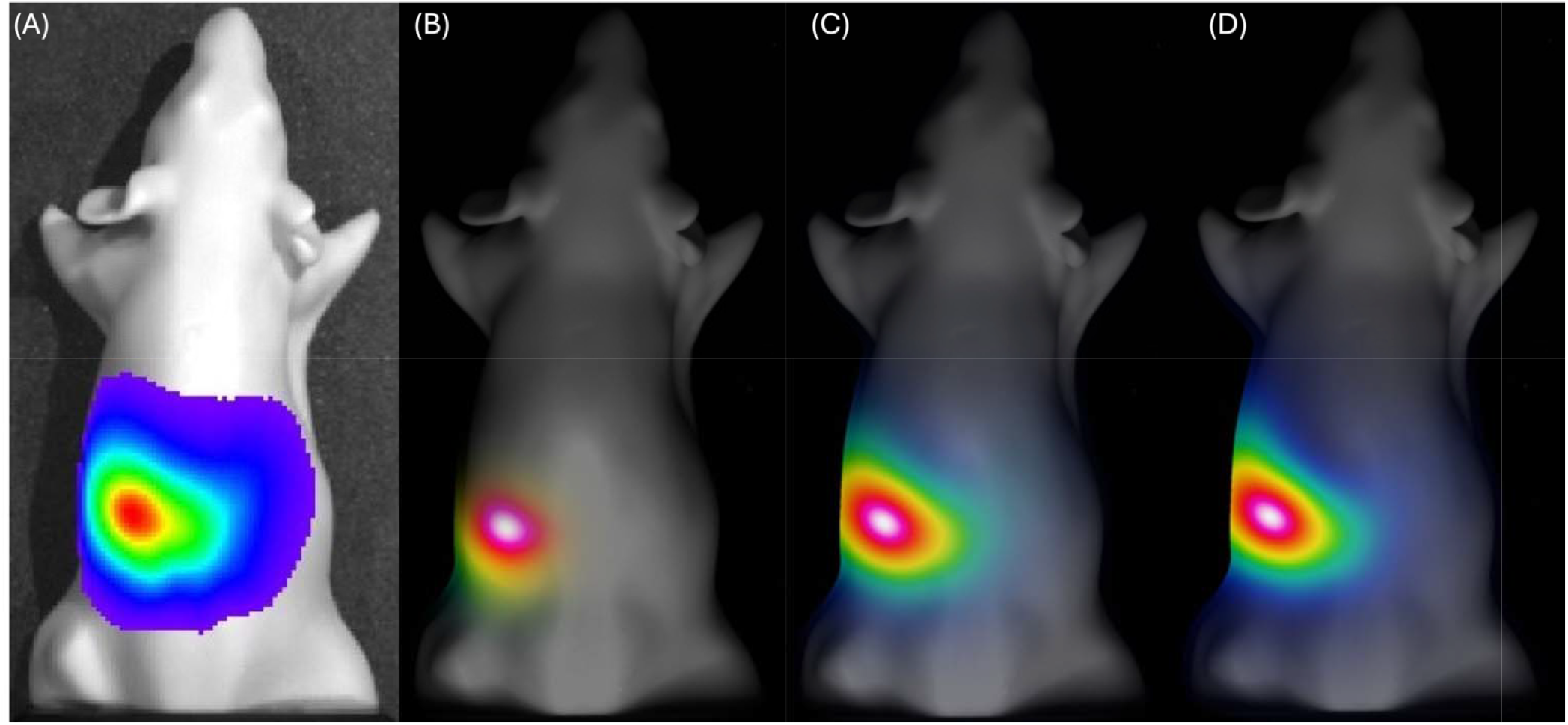
Imaging of the XPM-2 phantom with LED source “Mode A” active. (A) Reference image acquired with the Revvity IVIS Spectrum using the default autoexposure settings in Living Image software. (B-D) Corresponding AURORA-MSI **im**ages acquired with exposure times of 1 s, 10 s, and 30 s.

In both imaging systems, the source appears as a compact hotspot centered over the phantom’s midchest. As the AURORA-MSI exposure time increased, the signal-to-noise ratio of the central hotspot improved, and the surrounding diffuse emission halo expanded across a larger area of the phantom. This behavior is consistent with increased integration of multiply scattered photons within the tissue-mimicking phantom.^58,59^ This pattern, a compact high-intensity core surrounded by a broader, lower-intensity diffuse halo, is the expected signature of a buried optical source imaged through a scattering medium and is seen reproducibly in both the IVIS reference and the AURORA-MSI data.

As a self-contained biological luminescent source requiring no exogenous substrate administration, a commercially available bioluminescent petunia (*Firefly Petunia*, Light Bio, Inc) was imaged in complete darkness. The line carries a fungal bioluminescence gene cassette derived from *Neonothopanus nambi*, conferring continuous, metabolically driven green emission from leaves, stems, and immature blooms independent of external light cycling.^60,61^ Unlike luciferase-luciferin reporter systems, in which emission requires exogenous substrate delivery and follows characteristic post-injection decay kinetics,^57^ the fungal pathway sustains emission indefinitely via the plant’s endogenous caffeic acid cycle, making it a useful low-photon-flux benchmark for extended-integration imaging. A potted specimen was imaged in complete darkness across a series of exposures from 1 s to 1200 s (Figure 4). At short exposures, only the brightest flower centers were resolved above the read-noise floor; signal-to-noise ratio in the floral regions improved monotonically with integration time. Floral centers and petal margins showed elevated signal relative to petal interiors, a nonuniform luminescent distribution consistent with the known restriction of strong fungal bioluminescence pathway expression to actively metabolizing immature floral and stem tissue.^60,61^ A full-resolution detail of this fine spatial structure is provided in Figure S21. The *Firefly Petunia* dataset establishes the platform’s capacity to resolve fine spatial structure from a continuous, substrate-free biological luminescent source with full floral structure resolved at 300 s integration and negligible further gain beyond 600 s, independent of the injection-and-decay kinetics that constrain timelimited substrate-based assays.

**Figure 4.**
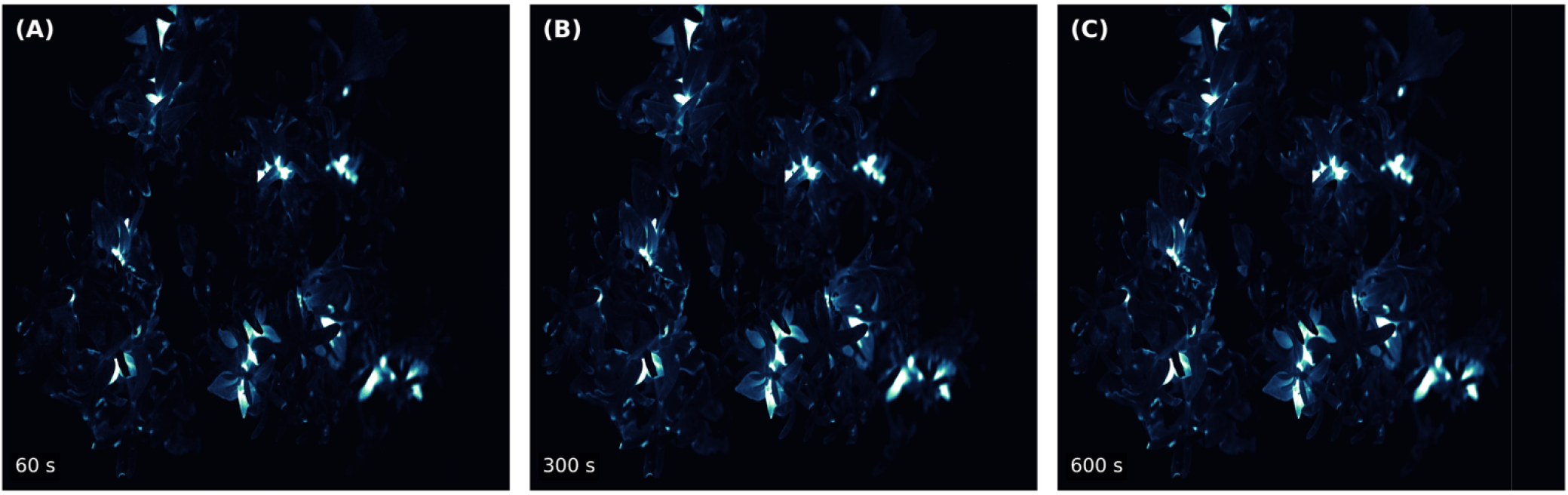
Bioluminescence imaging of a commercial *Firefly Petunia* (Light Bio, Inc.) acquired at increasing exposure times. (A) 60 s exposure, showing signal primarily in the brightest floral regions. (B) 300 s exposure, revealing the full luminescent structure of the plant. (C) 600 s exposure, producing little additional increase in the detected signal relative to (B).

To assess sensitivity at a biologically relevant scale closer to single-cell-population reporter assays, we imaged a 96-well plate containing a serial dilution of luciferase-expressing HSJD-GBM1-001 glioblastoma cells, with D-luciferin substrate added to one pair of replicate columns immediately before AURORA-MSI acquisition. The same plate was subsequently imaged on the IVIS Spectrum as a ground-truth reference, following addition of luciferin to a second, independent pair of replicate columns at the time of IVIS imaging (Figure S22). The IVIS system cleanly resolved all three cell-density tiers in both replicate wells. AURORA-MSI clearly resolved both replicate wells at the highest density but detected signal in only one of two replicate wells at each of the two lower densities. These results bracket the platform’s detection threshold for luciferase reporter signal under the acquisition conditions used: signal was detected in both replicate wells at 50,000 cells per well and in one of two wells at each lower density, whereas the IVIS Spectrum resolved all three tiers. This comparison is not instrument-matched since the two systems imaged different replicate columns of the same plate, dosed with luciferin at different times, and bioluminescent flux varies with time after substrate addition. Differences in detection outcome therefore reflect the combined contribution of instrument sensitivity, well-to-well biological variation, and substrate kinetics, and cannot be attributed to instrument sensitivity alone. With two replicate wells per density, these data bracket rather than quantify the detection threshold. A matched design with dosing-time-controlled acquisition on both instruments and n ≥ 6 wells per density is required to establish a limit of detection. The AURORA-MSI frame also contains features not corresponding to well positions, a diagonal cluster of small spots and one saturated high-intensity spot (Figure S22), whose origin has not been determined with candidate sources including hot-pixel events, plate-surface contamination, and stray reflection. These features are spatially disjoint from the analyzed wells and were excluded from ROI analysis, but their presence indicates that background rejection at the single-well detection limit is not yet fully characterized. AURORA-MSI is therefore suited to reporter detection at the higher end of this range, while quantitative discrimination near the threshold requires either longer integration or the cooled-camera optimizations discussed below.

Collectively, these studies demonstrate that AURORA-MSI supports bioluminescence imaging across phantom, cell-based, and biological specimen models spanning a broad range of emission intensities and source geometries. Concordance of the spatial emission profile with the IVIS Spectrum reference in phantom imaging, together with detection of luciferase reporter cells and of autonomous fungal-pathway emission in an intact plant, indicates that the platform provides sufficient sensitivity and spatial resolution for bioluminescence detection across phantom, in vitro, and whole-organism specimens. Extension to quantitative radiance measurement, and to longitudinal imaging in live animals, will require absolute radiometric crosscalibration against a certified luminescent standard and is not established here.

### Spectral Classification of Pharmaceutical and Chemically Similar Powders

To evaluate the platform’s capacity for material discrimination, ten pharmaceutical, food-grade, and household powders were imaged across all 14 excitation wavelengths. The panel deliberately includes two chemically overlapping pairs, two ibuprofen products of different formulation (Advil, generic ibuprofen) and two sucrose products of different particle size (granulated, powdered), to test whether excitation-resolved texture features respond to formulation and morphology as well as to composition. Multispectral image-texture features were analyzed via principal component analysis (PCA)^62^, a strategy previously validated for pharmaceutical identification, counterfeit detection, and illicit-substance screening^63–69^.

Each powder produces a distinctive intensity-wavelength fingerprint (Figure S23), with UV excitation (367 nm) yielding the greatest between-class contrast and NIR excitation (940 nm) encoding complementary diffuse-reflectance differences likely arising from crystal structure and organic versus inorganic composition.^70^ A suite of photometric and texture features, including mean ROI intensity, skewness, kurtosis, GLCM contrast^71^, and LBP entropy^72^, was extracted from each image (detailed in SI). Texture features proved orthogonal to mean intensity, carrying independent discriminating information linked to particle size distribution and surface morphology (Figure 5).

**Figure 5.**
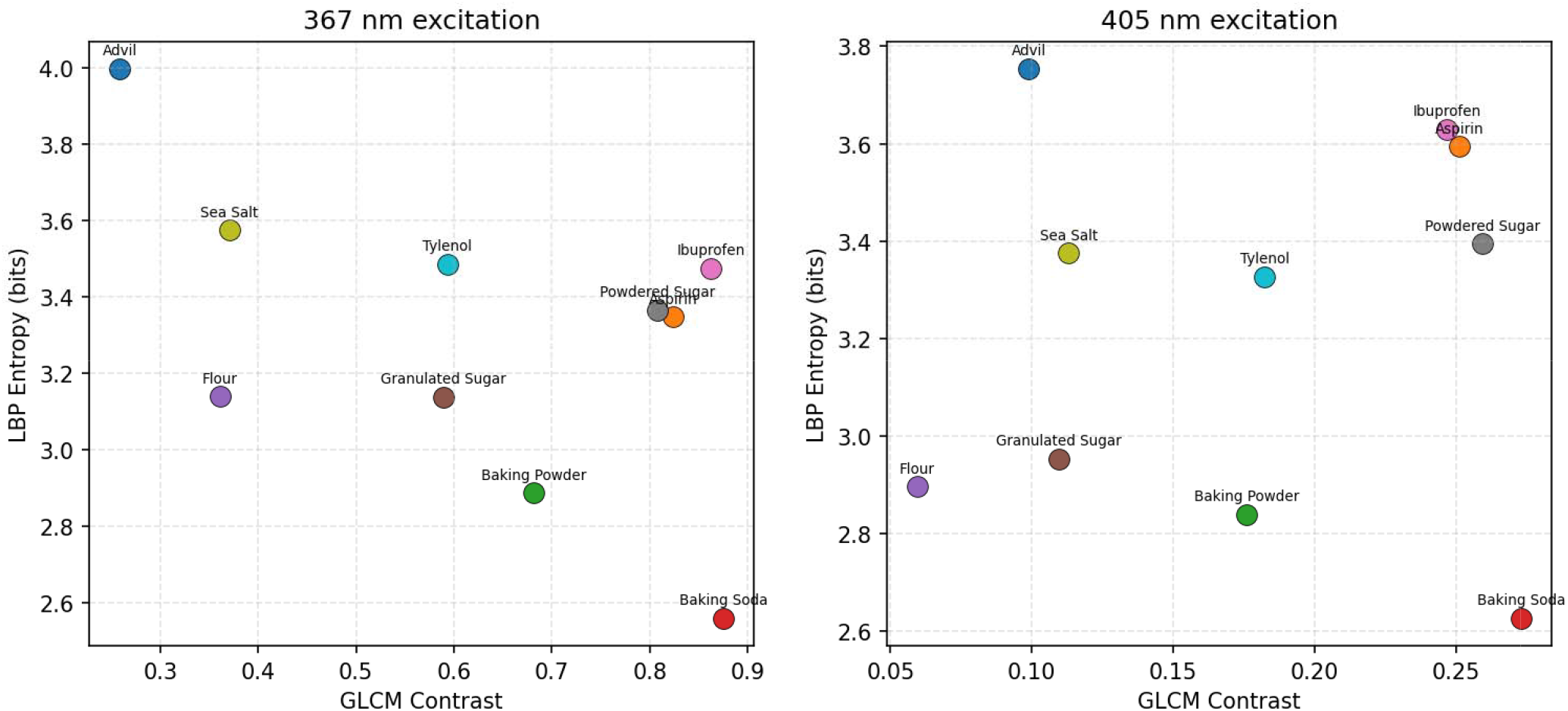
Texture feature space defined by gray-level co-occurrence matrix (GLCM) contrast and local binary pattern (LBP) entropy for powder ROIs acquired under 367 nm and 405 nm excitation. Each point represents the mean feature response of a powder class. The distributions indicate increased separation among powder classes in the two-dimensional texture feature space under short-wavelength excitation.

PCA of the full feature matrix resolved all ten powders in PC1-PC2 space, which together explain 88.4% of total variance (Figure S27). Wavelength-dependent higher-order intensity statistics (Figure S26) and supervised LDA projections (Figure S28) further support robust class discrimination. A Random Forest classifier trained on the multispectral feature matrix separated all ten powders without error under five-fold stratified cross-validation (Figure S29).^73^ Feature importance analysis (Figure S30) identifies skewness at 367 nm and mean intensity at 940 nm as the dominant discriminating features, consistent with UV excitation driving chromophore-specific absorption and fluorescence contrasts that differentiate aromatic API-containing tablets from inorganic and carbohydrate powders^74^, while NIR reflectance encodes bulk organic versus inorganic composition.^70^ These features must be interpreted with caution: saturation fractions of 0.19-0.24 were observed for several powders at 367 nm and 940 nm under the fixed exposure used (Table S4), and the higher-order statistics at these wavelengths partly reflect clipping of the intensity distribution at the sensor ceiling rather than intrinsic photophysical contrast (Figure S26).

Excitation wavelength modulates both the chromophoric absorption cross-section and fluorescence quantum yield of each compound, producing wavelength-dependent reflectance and emission signatures characteristic of chemical composition. For amorphous organic solids (Tylenol, ibuprofen), UV absorption by the active pharmaceutical ingredient is superimposed on broadband Mie scattering from the particle distribution, whereas for crystalline inorganic salts (NaCl, NaHCO□), the response is dominated by specular and diffuse reflectance with minimal UV absorption.^75^

Several limitations should be acknowledged. First, the dataset is small (10 powders, one batch per powder, one wellplate configuration), and performance on independent samples with different particle size distributions, polymorphic forms, or excipient blends is untested. Generalization across manufacturers and batches will require substantially expanded training sets. In addition, the absence of emission filters means that the detected signal is a convolution of reflected excitation, fluorescence emission, and phosphorescence; spectrally resolved emission would disambiguate these contributions and likely improve sensitivity.

### Wavelength-Selective Plant Imaging Anthocyanin Stress Monitoring

Anthocyanins are photoprotective vacuolar pigments and sensitive biomarkers of postharvest physiological decline in leafy crops.^76,77^ Since conventional quantification by pH-differential spectrophotometry or HPLC is destructive,^78,79^ we applied dual-wavelength gray-normalized reflectance imaging to monitor anthocyanin accumulation nondestructively in postharvest Lactuca sativa over seven days. The anthocyanin index (AI = log[R□/R□]^80,81^) was computed from spatially registered image pairs; gray-standard normalization corrected for inter sweep illumination drift, and leaf segmentation was performed by morphological image processing.

The spatially averaged anthocyanin index,, exhibited a systematic increase across the full seven-day postharvest monitoring period. Representative daily time points (Figure 6) displayed a progressive shift in the spatial AI distribution toward higher values. At day 1, values were relatively low and spatially uniform across the leaf lamina, consistent with physiologically intact tissue. By day 7, had increased substantially, with the spatial maps revealing a heterogeneous distribution and localized high-AI foci at leaf margins and midrib-adjacent zones, regions known to exhibit preferential anthocyanin accumulation under stress. ^76,81^

**Figure 6.**
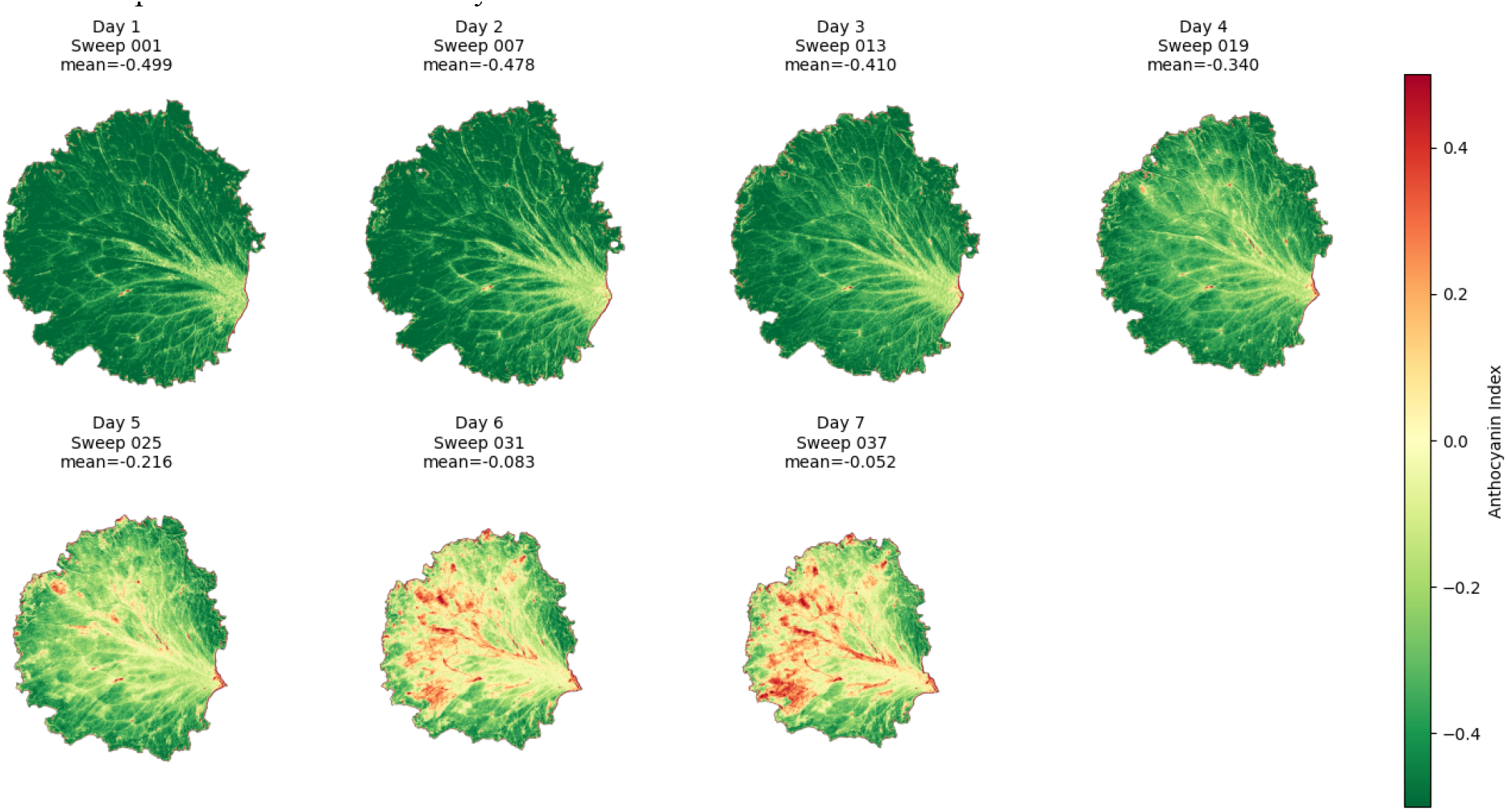
Temporal evolution of spatially averaged anthocyanin index (AI). Spatial maps and corresponding mean values (*Al* _*mean*_) are shown for representative daily time points over a seven-day monitoring period. *Al*_*mean*_ increased systematically with time, consistent with progressive postharvest changes in the tissue.

The full 42-sweep temporal series (Figures S31-S32) confirmed that the increase was not restricted to the daily representative images but occurred progressively across within-day acquisitions as well, indicating that anthocyanin accumulation was a continuous rather than threshold-triggered process under the experimental conditions. The monotonic increase in observed across the seven-day monitoring period is aligned with the documented stress-responsive anthocyanin biosynthesis in postharvest leafy vegetables. This observed AI increase is empirical, while the biosynthetic interpretation is mechanistic inference supported by prior literature. Because leaves were held without supplemental water, progressive dehydration co-occurred with any pigment change, and the two contributions are not separated by these measurements. In the absence of destructive reference quantification (pH-differential or HPLC) at matched time points, the measured index is reported as an empirical optical correlate of post-harvest decline rather than as a calibrated measure of anthocyanin content.

The spatial AI maps generated by this study reveal interleaf heterogeneity that is not captured by bulk extraction methods. The observed gradient, with higher AI values at leaf margins and midrib-adjacent tissues in later sweeps, agrees with the well-documented pattern of preferential anthocyanin localization in adaxial epidermal cells exposed to incident radiation, as well as with the spatial distribution of dehydration-induced water loss in leaf tissue.^76,82^ This capacity to map, rather than merely average, anthocyanin distribution provides a basis for region-specific quality scoring relevant to consumer-facing produce assessment.

### Depth-Dependent Optical Contrast

The broad spectral range of the illumination system permits separation of surface-localized and subsurface optical contrast within a single imaging geometry, exploiting the strong wavelength dependence of effective penetration depth in plant tissue. ^83–89^ Succulent specimens imaged under UV-A excitation (Figure 7A) showed emission concentrated at leaf surfaces and margins. Ratio contrast was concentrated at leaf edges and on leaf surfaces rather than in the leaf interior. Because 367 nm is more strongly absorbed by epidermal phenolics and by the cuticle than 405 nm,^83,84^ the 367/405 ratio is expected to be sensitive to spatial variations in the thickness or phenolic loading of the cuticle-epidermis complex.

**Figure 7.**
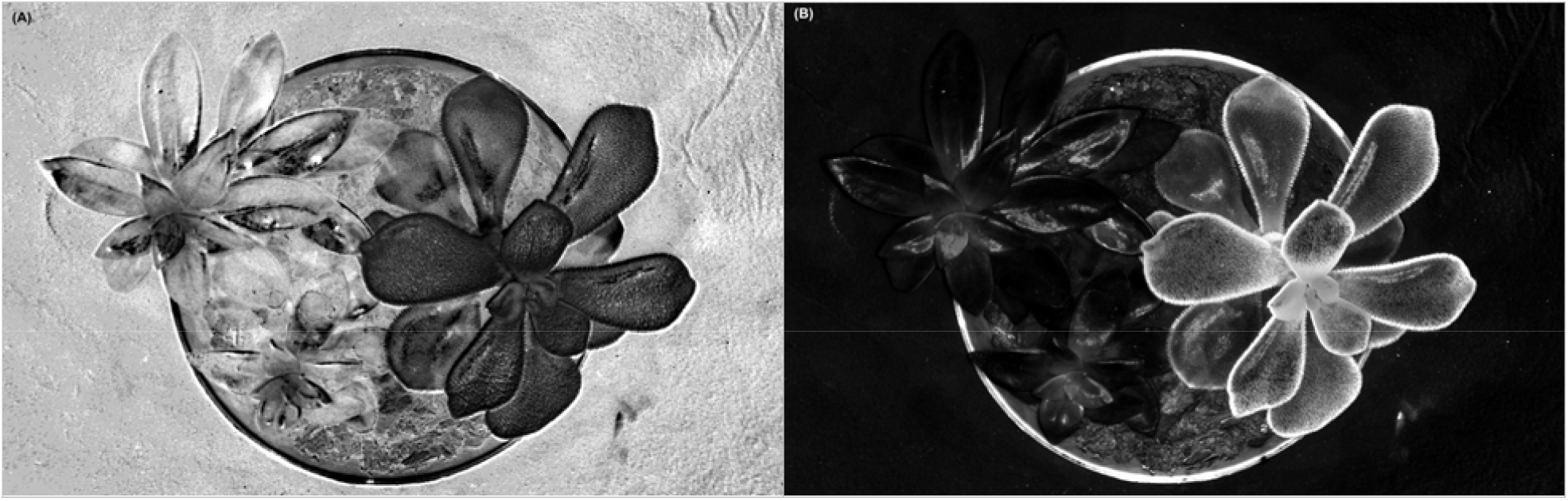
Excitation-ratio imaging reveals surface fluorescence localization and depth-dependent light transport. (A) Ratiometric images acquired under ultraviolet excitation: 367/405 nm. Based on the shallow optical penetration depth of these wavelengths, the measured signals are expected to be dominated by surface and near-surface fluorophores (e.g., the cuticle, phenolics, and epidermal layers). Ratioing suppresses geometric and illumination heterogeneity while enhancing contrast consistent with differences in relative excitation efficiency and surface chemical composition. (B) Ratiometric images combining visible and near-infrared excitation: 530/850 nm. Because optical penetration depth varies substantially with excitation wavelength, these ratios are expected to sample different tissue depths, with NIR excitation contributing relatively greater sensitivity to subsurface structure and bulk scattering. Accordingly, the resulting ratios are interpreted as reflecting wavelength-dependent light transport rather than absolute reflectance. This interpretation is based on established leaf optics literature and the qualitative appearance of the images and was not independently validated here.

VIS/NIR ratio images (Figure 7B) displayed large spatial dynamic range, with high-ratio zones co-localizing with visually dense or pigmented surface features and low-ratio zones co-localizing with translucent tissue consistent with strong NIR backscattering from spongy mesophyll.^86,90^ We emphasize that the surface-to-subsurface depth-contrast interpretation is supported by the leaf optics literature and the qualitative appearance of the images; it is not validated here by crosssectional imaging. Inverted renderings (Figure S33) are provided for perceptual comparison; it encodes no additional physical information.

Multispectral imaging of cilantro (Coriandrum sativum) seedlings grown in soil provided a direct test of our system’s capacity to resolve spectrally distinct biological and pedogenic materials within a single field of view. The grayscale meanintensity image (Figure 8A) resolved root architecture structurally but with limited root-soil spectral contrast, consistent with the comparable scattering coefficients of root cortical tissue and mineral soil particles. ^91^

**Figure 8.**
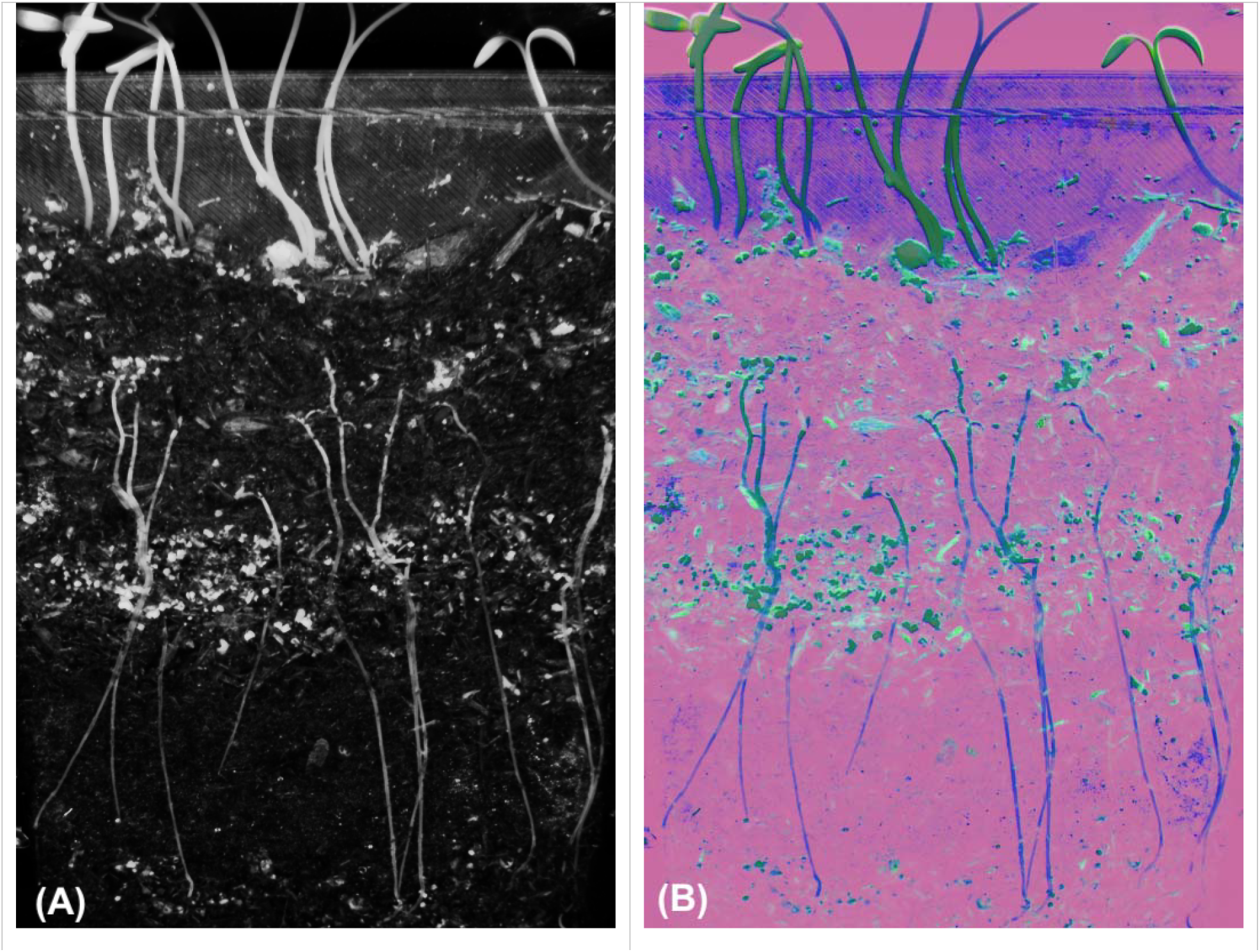
Multispectral imaging of cilantro (Coriandrum sativum) roots in soil. (A) Grayscale mean-intensity image generated by averaging all wavelength bands in the multispectral image stack, providing a structural overview of the sample. (B) PCA false-color composite constructed from the first three principal components (PC1 = red, PC2 = green, PC3 = blue). The PCA representation highlights spectral variation within the dataset, increasing visual contrast among regions with differing spectral characteristics.

PCA of the full multispectral stack (Figure 8B) achieved unsupervised separation of three material classes without supervised training or spectral library input. In the false-color composite rendered from the first three principal components (PC1 = red, PC2 = green, PC3 = blue), the bulk soil mineral matrix is represented as a uniform magenta field, root and hypocotyl tissue in dark blue^92,93^), and chlorophyll-bearing shoot tissue above the soil interface in saturated green.^94^ Cyan and teal speckles distributed throughout the subsurface zone are interpreted as spectrally intermediate features suggestive of organic matter fragments or moisture-laden aggregates. Collectively, these results demonstrate that PCA-based dimensionality reduction, applied to a multispectral stack acquired with the spectral illumination system described here, achieves unsupervised separation of biologically and pedogenically distinct material classes without any supervised training or prior spectral library input.

### Platform Capabilities and Future Development

AURORA-MSI provides a unified framework for calibrated spectral excitation, large-field-of-view mapping, angular reflectance, and time-resolved imaging within a single opensource instrument. Inverting the conventional multispectral paradigm by resolving spectral content through excitation rather than emission enhances the ability to characterize excitation-specific photophysical responses compared with approaches employing broadband illumination. The combination of ultraviolet (367, 405 nm), visible (448-655 nm), and nearinfrared (740, 850, 940 nm) excitation channels span a spectral range that encompasses electronic, vibrational, and combination absorption features relevant to both biological chromophores and synthetic organic compounds. The azimuthally sectorized ring illumination geometry allows, within a single instrument, both the directional illumination required for photometric-stereo reconstruction (Figure S34-S35) and the near-symmetric illumination required for quantitative flat-field-corrected reflectance mapping.

Several practical limitations of the current implementation motivate near-term development. In the present configuration, emission-filter exchange and the acquisition of excitation-emission matrices or time-resolved filter sequences require manual intervention between steps, constraining acquisition throughput and reproducibility. A motorized filter wheel, controlled through the existing Python software stack, will enable fully automated acquisition of excitation-emission matrices and time-resolved filter sequences without manual intervention. In addition, the present emission-filter set is limited to long-pass cutoffs; as noted in the powder classification results, the absence of narrower emission filtering means the detected signal in that demonstration is a convolution of reflected excitation, fluorescence emission, and phosphorescence, which likely reduces achievable classification sensitivity. Expanded emission-filtering options, including narrow-bandpass filters, will disambiguate these contributions and extend the system to Raman-shifted and anti-Stokes detection applications.

LED channel addressing and camera exposure triggering are currently coordinated through software polling rather than hardware interrupts, introducing frame-to-frame timing jitter and requiring conservative software-timed delays between LED activation and image acquisition to ensure the illumination source has stabilized before each exposure, adding unnecessary acquisition overhead. This is adequate for the multisecond-to-multi-minute decay kinetics characterized in this work, but it constrains temporal precision and throughput for faster luminescence processes. Tighter synchronization between LED channel addressing and camera exposure triggering, implemented through hardware trigger lines and interrupts rather than software polling, will reduce interframe timing jitter and improve temporal precision in fast phosphorescence decay measurements.

Specular reflectance from reflective sample containers (e.g., glossy or reflective petri dishes, well-plate walls) and specimen surfaces (e.g., powder surfaces, leaf cuticle) can superimpose a directly reflected excitation component on the diffuse-reflectance and fluorescence signal of interest, potentially biasing measurements in the affected modalities. Using crossed linear polarizers (one over the LED illumination sources, combined with one in front of the camera) can help to suppress this reflection (Figure S36).

## Conclusion

We have designed, built, and validated AURORA-MSI, a multispectral imaging platform that integrates fourteen-wavelength narrowband excitation spanning 367-940 nm, eight-sector azimuthal angular illumination control, and a thermoelectrically cooled monochrome detection pathway within a fully 3D-printed, reproducible mechanical assembly. The platform supports reflectance, fluorescence, bioluminescence, and time-resolved phosphorescence imaging through interchangeable long-pass emission filters. Its quantitative performance was established through systematic radiometric calibration across all wavelength and sector configurations, spatial uniformity mapping, and characterization of detector noise and linearity as functions of sensor temperature. The platform was subsequently evaluated in complementary experiments spanning chemical and biomedical imaging: bioluminescence detection benchmarked against a commercial IVIS Spectrum across phantom, cell-based, and whole-organism specimens; time-resolved phosphorescence decay imaging; and excitation-resolved spectral classification of visually similar pharmaceutical powders. Two further experiments, dual-excitation fluorescence fingerprinting of mineral specimens and wavelength-selective plant-tissue contrast imaging, established generality across specimen classes.

By inverting the conventional detection-side multispectral paradigm, AURORA-MSI enables excitation-specific photophysical measurements across diverse application domains, from bioluminescence detection and phosphorescence lifetime imaging to pharmaceutical powder classification and mineral fluorescence fingerprinting. This approach allows information complementary to that obtained with conventional detection-side multispectral systems, and the two modalities can be combined when both excitation and emission spectral resolution are required. The ability to discriminate ten visually indistinguishable white powders using multispectral texture analysis without chemical preparation or reagents indicates the potential applicability of the platform to rapid, noncontact screening in forensic, pharmaceutical, and food-safety contexts. Likewise, benchmarking against a commercial IVIS system demonstrated comparable localization of bioluminescent sources in a tissue-mimicking phantom, supporting the feasibility of open-source instrumentation for preclinical optical imaging applications.

## ASSOCIATED CONTENT

### Supporting Information

The Supporting Information is available free of charge on the ACS Publications website.

## AUTHOR INFORMATION

### Author Contributions

Conceptualization: J.M.B., N.J.S., A.L.G., K.J.C.

Methodology: J.M.B., C.L.L., I.A.H., T.W., G.L., K.J.C. Software: J.M.B., C.L.L., K.J.C.

Validation: J.M.B., C.L.L., I.A.H., K.J.C.

Formal Analysis: J.M.B., C.L.L., K.J.C. Investigation: J.M.B., C.L.L., I.A.H., T.W., G.L.

Resources: J.M.B., T.W., G.L., N.J.S., A.L.G., K.J.C.

Data Curation: J.M.B.

Writing - Original Draft: J.M.B., C.L.L.

Writing - Review & Editing: J.M.B., C.L.L., T.W., G.L., N. J.S., A.L.G., K.J.C.

Visualization: J.M.B., C.L.L., T.W., K.J.C.

Supervision: N.J.S., A.L.G., K.J.C.

Project Administration: N.J.S., A.L.G., K.J.C. Funding Acquisition: N.J.S., A.L.G., K.J.C.

All authors have given approval to the final version of the manuscript

## ACKNOWLEDGMENT

The authors thank Logan Ruthardt for valuable discussions and feedback on experimental design throughout this work, and Adrian Mendonsa for helpful discussions and constructive feedback that helped ground the conceptual framework of this study.

