## Supporting Information for "An Open-Source, Excitation-Resolved UV-NIR Imaging Platform for Bioluminescence, Luminescence-Decay, and Chemical Discrimination Measurements"

2. The MITRE Corporation, Bedford, Massachusetts, USA. The author’s affiliation with The MITRE Corporation is for identification purposes only and is not intended to convey or imply MITRE’s concurrence with, or support for, the positions, opinions, or viewpoints expressed by the author. Approved for Public Release, Distribution Unlimited. Public Release Case Number 26-0987. ©2026 The MITRE Corporation. All Rights Reserved.

3. Mechanical Engineering Department, Colorado School of Mines, Golden, CO, USA.

4. Animal Imaging Irradiation Shared Resource, University of Colorado Comprehensive Cancer Center, University of Colorado Anschutz, Aurora, CO, USA

5. Morgan Adams Foundation Pediatric Brain Tumor Research Program, Department of Pediatrics, University of Colorado School of Medicine, Aurora, Colorado.

6. Department of Radiology, University of Colorado Anschutz, Aurora, CO, USA

7. Center for Cancer and Blood Disorders, Children’s Hospital Colorado, Aurora, Colorado

8. Chemical and Biological Engineering Department, Colorado School of Mines, Golden, CO, USA.

****1. Instrumentation****

Figure S1. AURORA-MSI platform and hardware components

Figure S2. Excitation LED illumination patterns

2. System Calibration

2.1 Spectral Sensitivity and Exposure Linearity

Figure S3. Representative calibration image

Figure S4. Gray step-wedge intensity profiles vs. wavelength and exposure time

Figure S5. Sensor response characterization from step-wedge data

2.2 Optical Power Linearity and Angular Uniformity

Figure S6. Optical power spectral distribution by wavelength at maximum drive intensity

Figure S7. Spectral power response of individual optical enable zones vs. drive intensity

Figure S8. Linearity (R²) of optical power response across wavelength-zone combinations

3. Mineral Fluorescence Fingerprinting

Table S1. Representative fluorescence intensities of mineral specimens

Figure S9. Effect of excitation wavelength on mineral fingerprint separability

Figure S10. Normalized fluorescence emission fingerprints of mineral specimens

Figure S11. Per-sample band-differenced emission spectra

Figure S12. Excitation-dependent emission spectral signatures

4. Time-Resolved Phosphorescence Decay Imaging

Figure S13. Normalized tri-exponential phosphorescence decay curves

Figure S14. Tri-exponential lifetime components

Table S2. Tri-exponential decay parameters by phosphor channel

Figure S15. Phosphorescence decay curves

Table S3. Comparison of phosphorescence decay models

Figure S16. Comparison of phosphorescence decay models

Figure S17. Goodness-of-fit comparison across decay models

Figure S18. Residual analysis of phosphorescence decay models

Figure S19. Temporal signal-to-noise ratio of phosphorescence measurements

5. Bioluminescence Imaging

Figure S20. Imaging of the IVIS XPM-2 phantom with LED source "Mode B"

Figure S21. Full-resolution detail of Firefly Petunia bioluminescence image

Figure S22. Bioluminescence imaging of a luciferase-expressing HSJD-GBM1-001 cell dilution series

6. Spectral Classification of Pharmaceutical and Chemically Similar Powders

Figure S23. Excitation-dependent intensity fingerprints of powder samples

Table S4. Frame-level image descriptors for representative powder samples

Figure S24. Repeatability of powder measurements across excitation wavelengths

Figure S25. System-level summary of signal magnitude and repeatability

Figure S26. Higher-order intensity statistics (skewness/kurtosis) of powder images

Figure S26. Principal component analysis of multispectral powder features

Figure S28. Linear discriminant analysis of multispectral powder features

Figure S29. Random Forest classification performance (confusion matrix)

Figure S30. Random Forest feature importance

7. Wavelength-Selective Plant Imaging

7.1 Anthocyanin Stress Monitoring

Figure 31. Temporal evolution of the anthocyanin index in lettuce

Figure 32. Temporal progression of the anthocyanin index

7.2 Wavelength-Selective Plant-Tissue Imaging

Figure 33. Inverted and supplementary excitation-ratio renderings

Figure 34. Spatial spectral response of a succulent plant across eight zones

Figure 35. Spatial spectral response of 3D-printed mouse phantom across eight zones

8. Light Polarization Effects

Figure S36. Cross-polarization suppression of specular glare

9. References

### Instrumentation

The AURORA-MSI system comprises a thermoelectrically cooled monochrome camera, a fourteen-wavelength LED excitation array (367-940 nm), eight independently addressable illumination sectors, interchangeable emission-filter mounts, and a height-adjustable sample platform (Figures S1 and S2). The complete optomechanical assembly, electronics, and illumination hardware are shown in Figures S1-S2 and included as additional supporting information files.

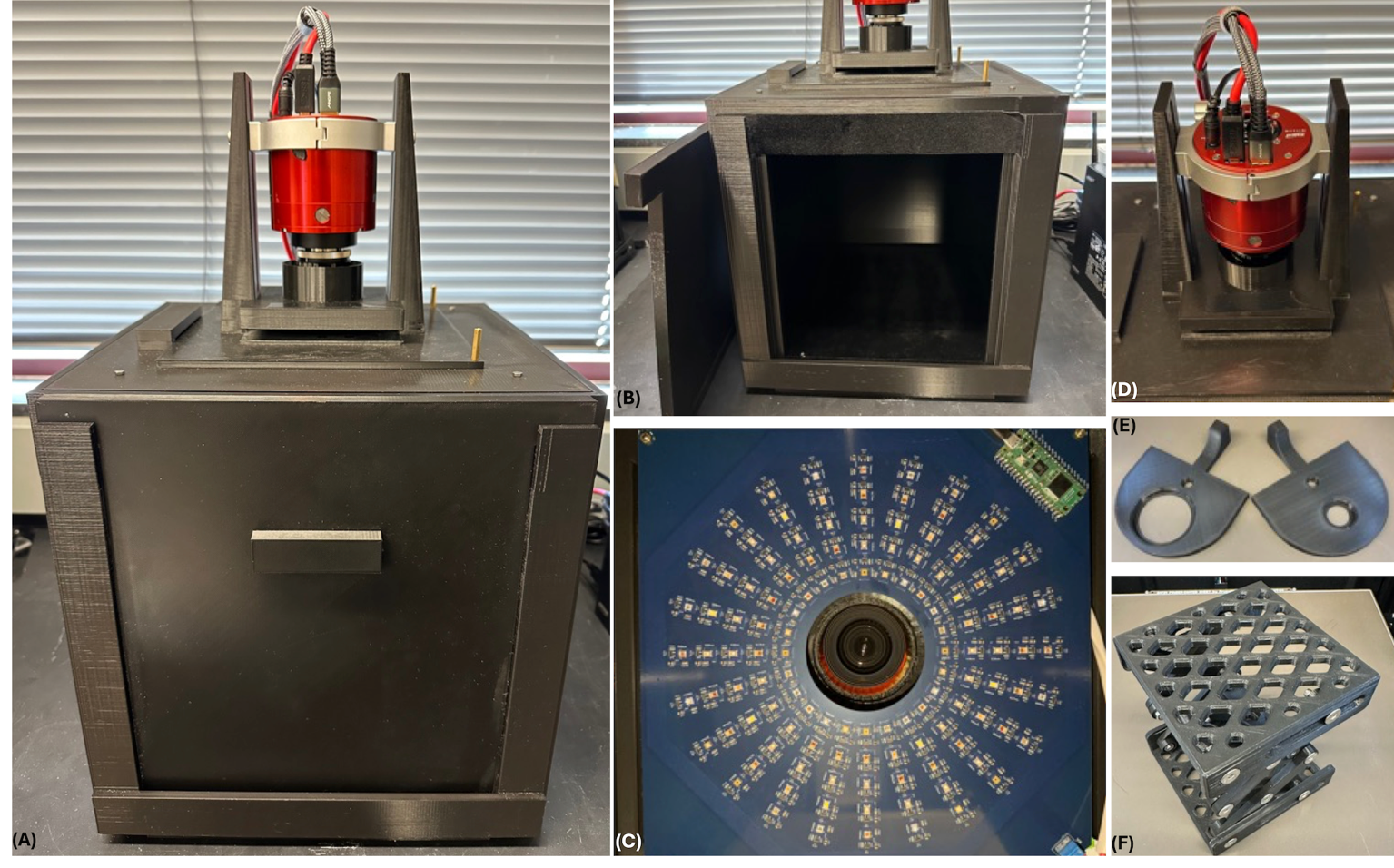

Figure S1. AURORA-MSI platform and hardware components. (A) Complete imaging system with top-mounted camera. (B) Imaging enclosure with the front panel removed. (C) LED control electronics and microcontroller. (D) Internal optomechanical assembly. (E) Interchangeable 3D-printed filter holders for 58 mm threaded and 25 mm unmounted optical filters. (F) Height-adjustable 3D-printed sample platform.

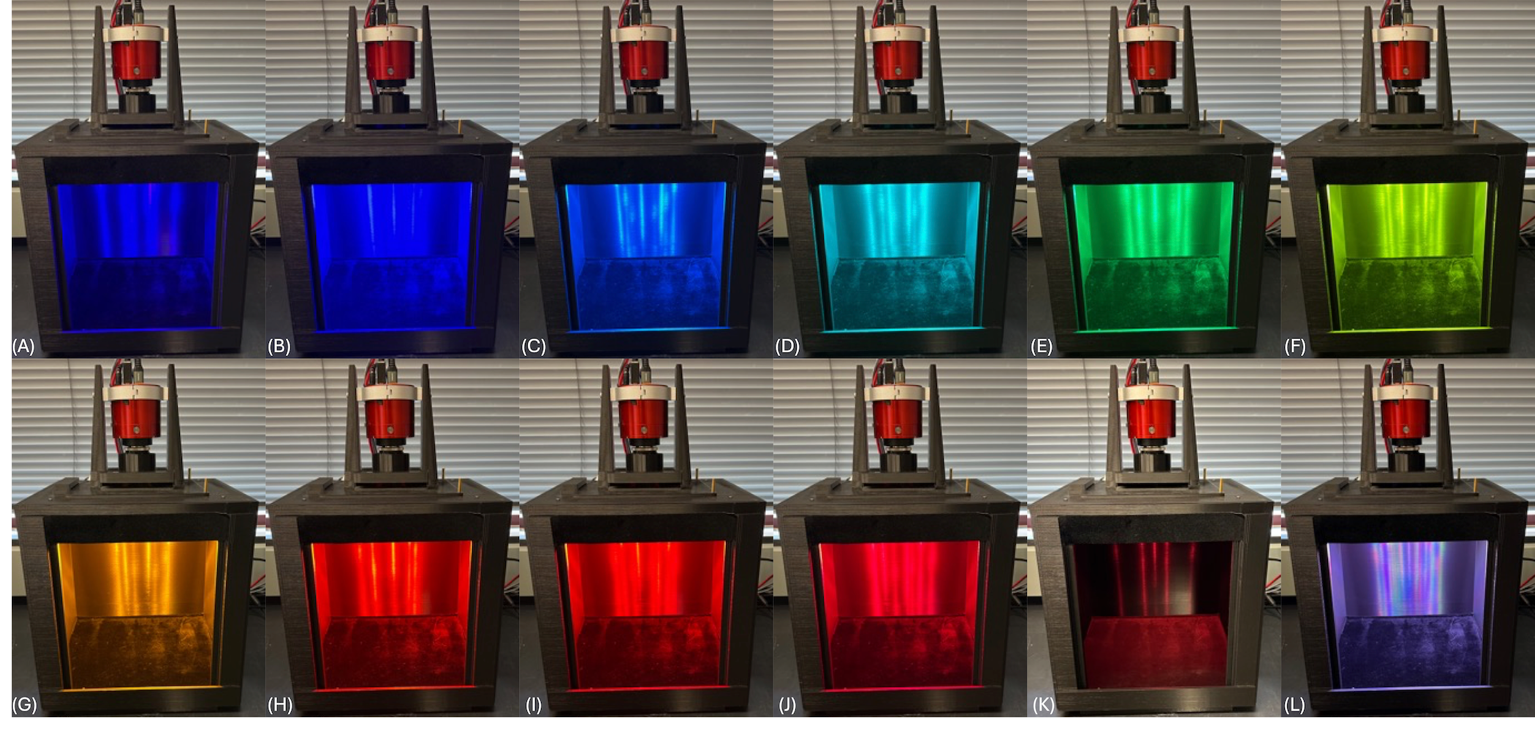

Figure S2. The LEDs can illuminate the sample across the UV-VIS-NIR spectral range. Shown here are illumination patterns acquired with individual LEDs at full output for (A) 405 nm, (B) 448 nm, (C) 470 nm, (D) 505 nm, (E) 530 nm, (F) 568 nm, (G) 591 nm, (H) 617 nm, (I) 627 nm, (J) 655 nm, and (K) 740 nm. (L) Composite image with the 405-740 nm LEDs operated simultaneously at 20% output. Images of the 280 nm, 367 nm, 850 nm, and 940 nm LEDs are not shown because their emission lies largely outside the spectral sensitivity of the camera used to document the array.

### System Calibration

#### Spectral Sensitivity and Exposure Linearity

A DGK Color Tools High Resolution SD Professional Lens Test Chart was imaged at 14 excitation wavelengths (367-940 nm) using five exposure times (1-20 ms) with fixed camera gain (100) and full LED output. Mean digital number (DN) was measured for each grayscale step and used to assess detector linearity over the unsaturated exposure range and to determine image dynamic range.

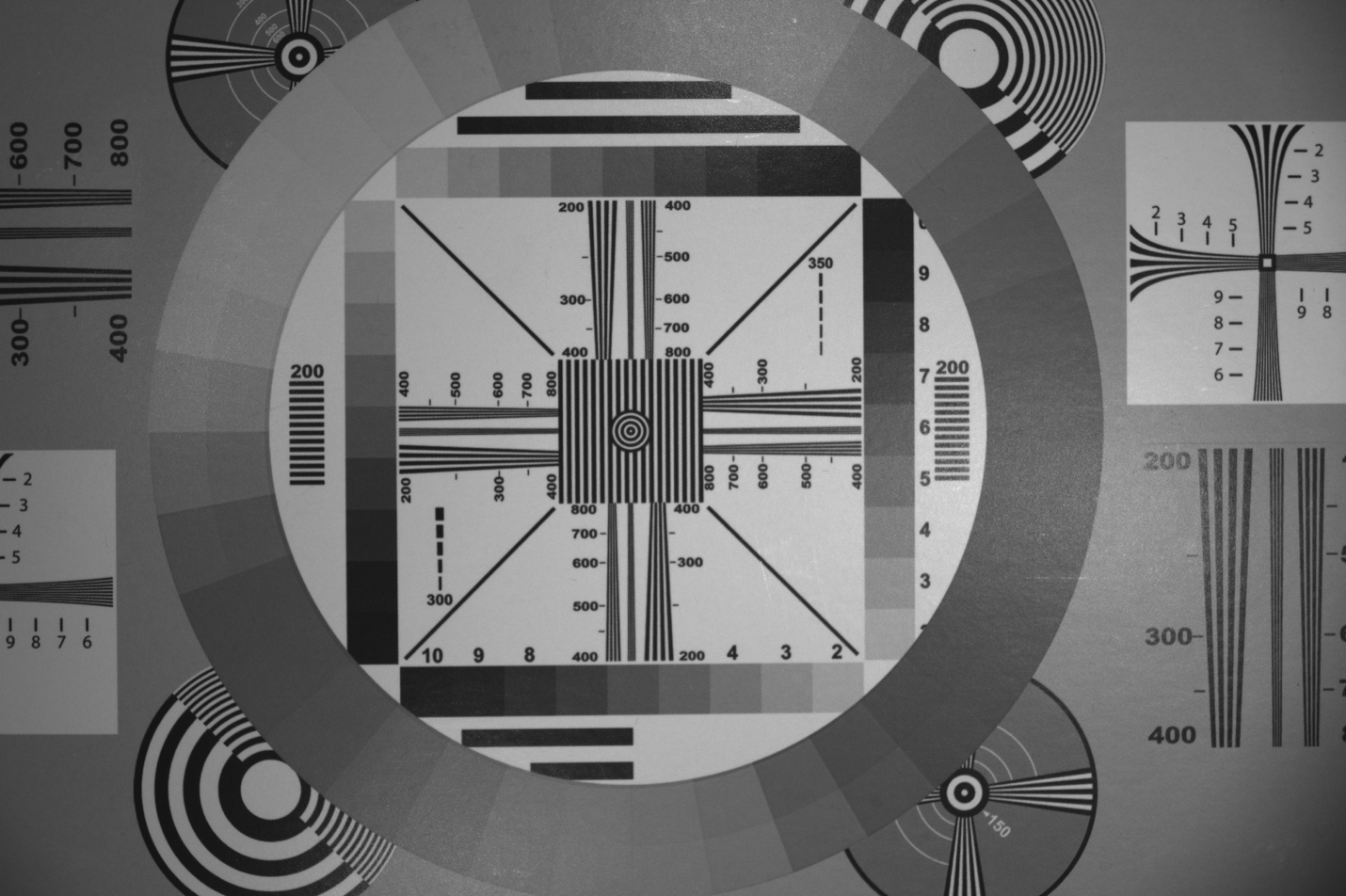

Figure S3. Representative calibration image. Image of the DGK Color Tools High Resolution SD Professional Lens Test Chart acquired under simultaneous 448, 530, and 627 nm illumination. The calibration target provides reference features for assessing spatial resolution, grayscale response, and illumination uniformity.

Measured system optical response varied with excitation wavelength and remained linear throughout the unsaturated operating range (Figures S4-S5). Response compression was observed only at the longest exposure times for the brightest grayscale targets under visible illumination.

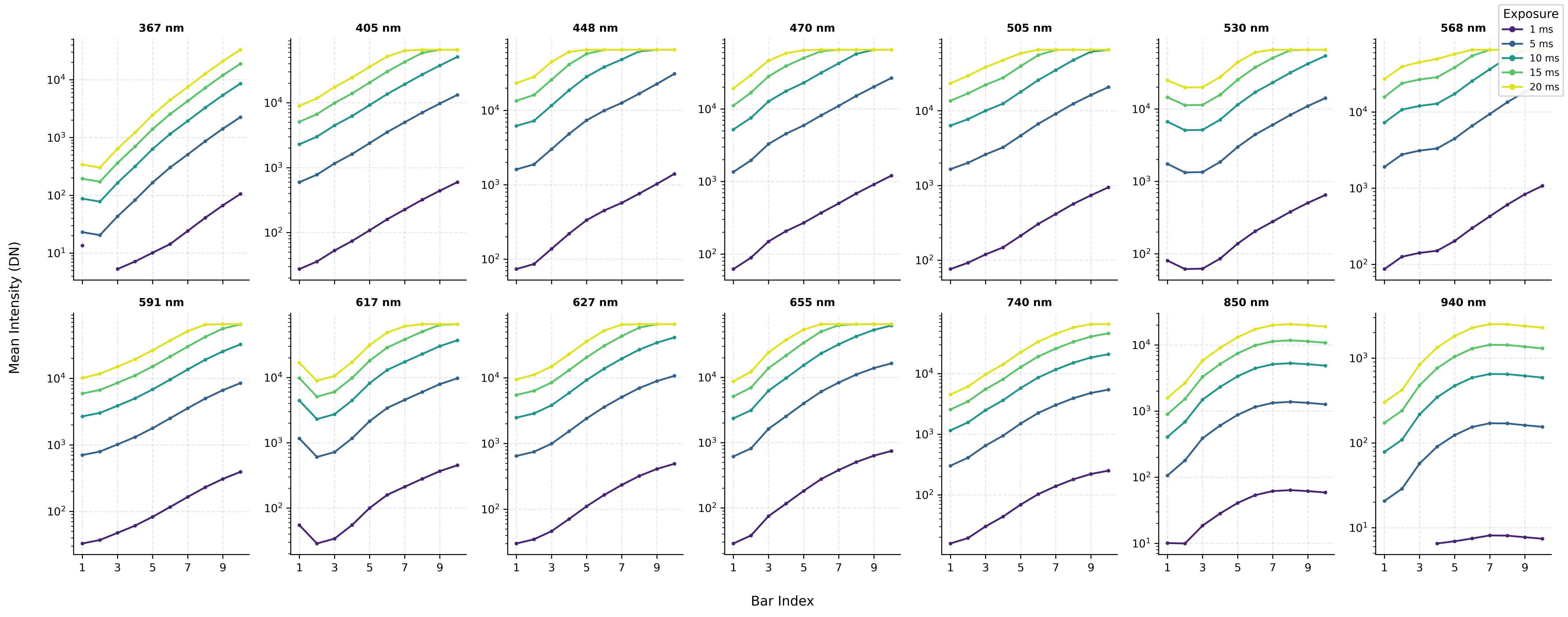

Figure S4**. Gray step-wedge intensity profiles across wavelength and exposure time.** Each panel plots the mean pixel intensity (DN, logarithmic scale) for each of the 10 step-wedge bars, averaged across four perimeter ROIs (top, bottom, left, and right strips), with each ROI independently rotated to a canonical dark-to-bright left-to-right gradient orientation prior to averaging. Line color encodes exposure time from dark blue (1 ms) to yellow (20 ms). Monotonically increasing profiles are consistent with the known geometric reflectance progression of the step wedge. Flattening of profiles at high bar indices and long exposures in the visible range indicates ADC saturation near the 16-bit ceiling.

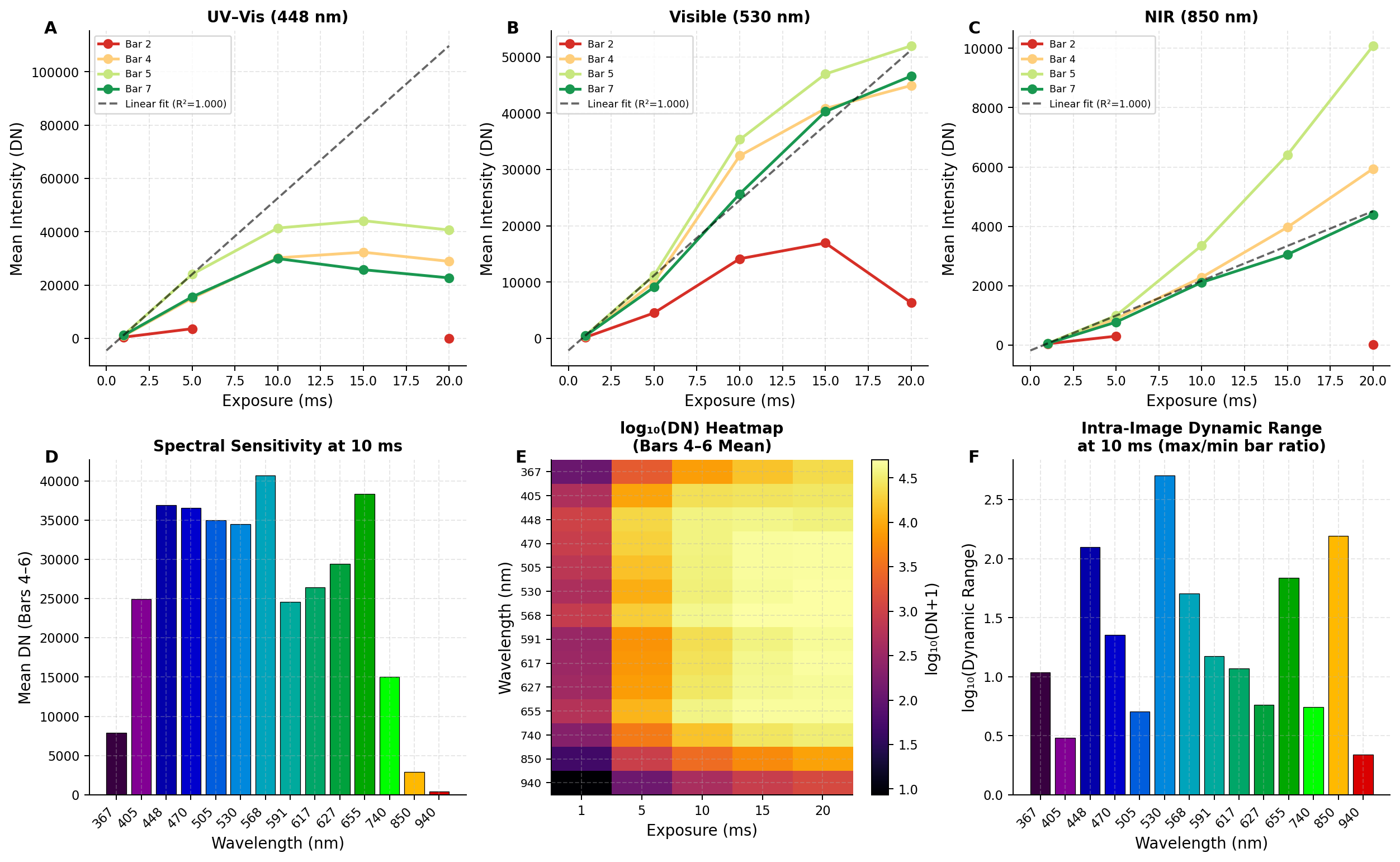

Figure S5. Sensor response characterization derived from gray step-wedge measurements at 14 illumination wavelengths and five exposure times (1-20 ms). (A-C) Mean intensity as a function of exposure time for four representative step-wedge bars (2, 4, 5, 7) at 448 nm, 530 nm and 850 nm. The dashed line shows a linear fit extrapolated from the two shortest exposures (1 and 5 ms). Departure of brighter bars from this extrapolation at exposures indicates ADC saturation. (D) Spectral sensitivity at 10 ms, quantified as the mean intensity of bars 4-6. Peak sensitivity occurs at 448-470 nm, with a secondary visible peak near 655 nm. Sensitivity at 367 nm and 940 nm is reduced by approximately one and two orders of magnitude consistent with the quantum efficiency of the camera. (E) log₁₀(intensity + 1) as a function of wavelength and exposure time, confirming that signal scales monotonically with exposure across all wavelengths and that spectral sensitivity variations span more than two decades within the 14-channel set. (F) Intra-image dynamic range at 10 ms, expressed as log₁₀(max intensity / min intensity). The elevated value at 367 nm (~130:1) reflects noise-floor proximity of the darkest bars at this wavelength rather than a genuine increase in chart contrast and should be interpreted with caution. Most visible channels yield intra-image dynamic ranges of 10-25:1 under this exposure condition.

#### Optical Power Linearity and Angular Uniformity

The AURORA-MSI optical excitation subsystem was characterized across its complete full wavelength range (280-940 nm) to establish absolute irradiance values and validate radiometric assumptions underlying quantitative imaging protocols. At maximum driver intensity (100% PWM), absolute optical power exhibited a 2.8-fold spectral variation, with peak output at 405 nm and secondary peaks at 448 nm and 568 nm, reflecting inherent differences in narrowband LED source efficiency and optical coupling design (Figure S6). Excellent linearity was maintained across all eight individual optical enable zones and across the complete wavelength set, enabling straightforward exposure-time normalization without correction factors (Figure S7). The spectral response was highly linear across the full driver intensity range (10-100% PWM), with mean coefficient of determination R² = 0.974 ± 0.081 across all 120 wavelength-enable zone combinations, indicating that optical power scales proportionally with PWM intensity in 88% of measured conditions (Figure S8). Low linearity regions (R² < 0.85, ~3% of conditions) were localized to detector saturation zones at high-power short wavelengths (405, 448 nm) and incomplete measurement sweeps at low-power long wavelengths (940 nm). These measurements were excluded from quantitative analysis. These absolute irradiance values and validated linearity characteristics form the foundation for radiometric normalization in all subsequent fluorescence lifetime decay and reflectance-based sample classification analyses.

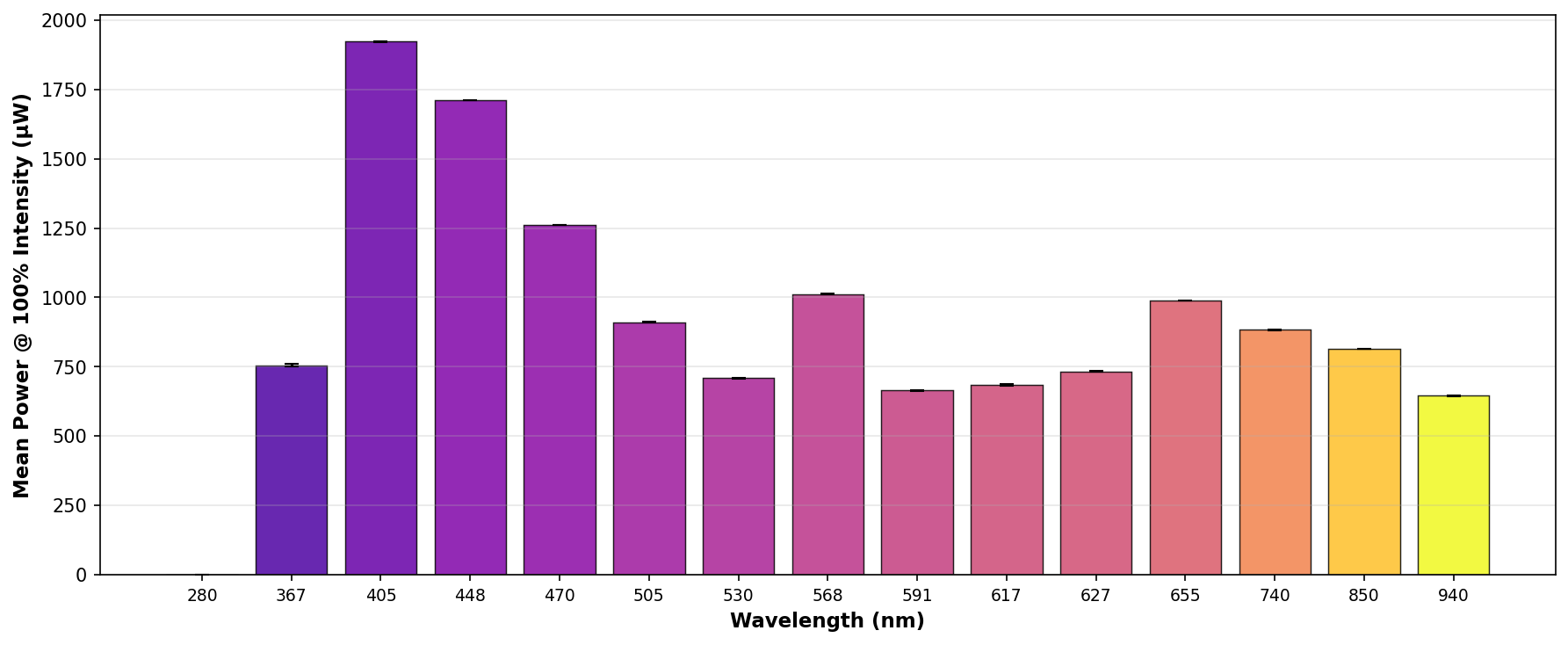

Figure S6. Optical Power Spectral Distribution by Wavelength. Absolute optical power (microwatts) delivered by each of the narrowband LED illumination modules, measured at maximum driver intensity (100% PWM). Wavelength-dependent power variation (2.8-fold range) reflects differences in LED source efficiency and optical coupling design. These absolute irradiance values are used for radiometric normalization in all quantitative fluorescence lifetime and reflectance analyses presented herein.

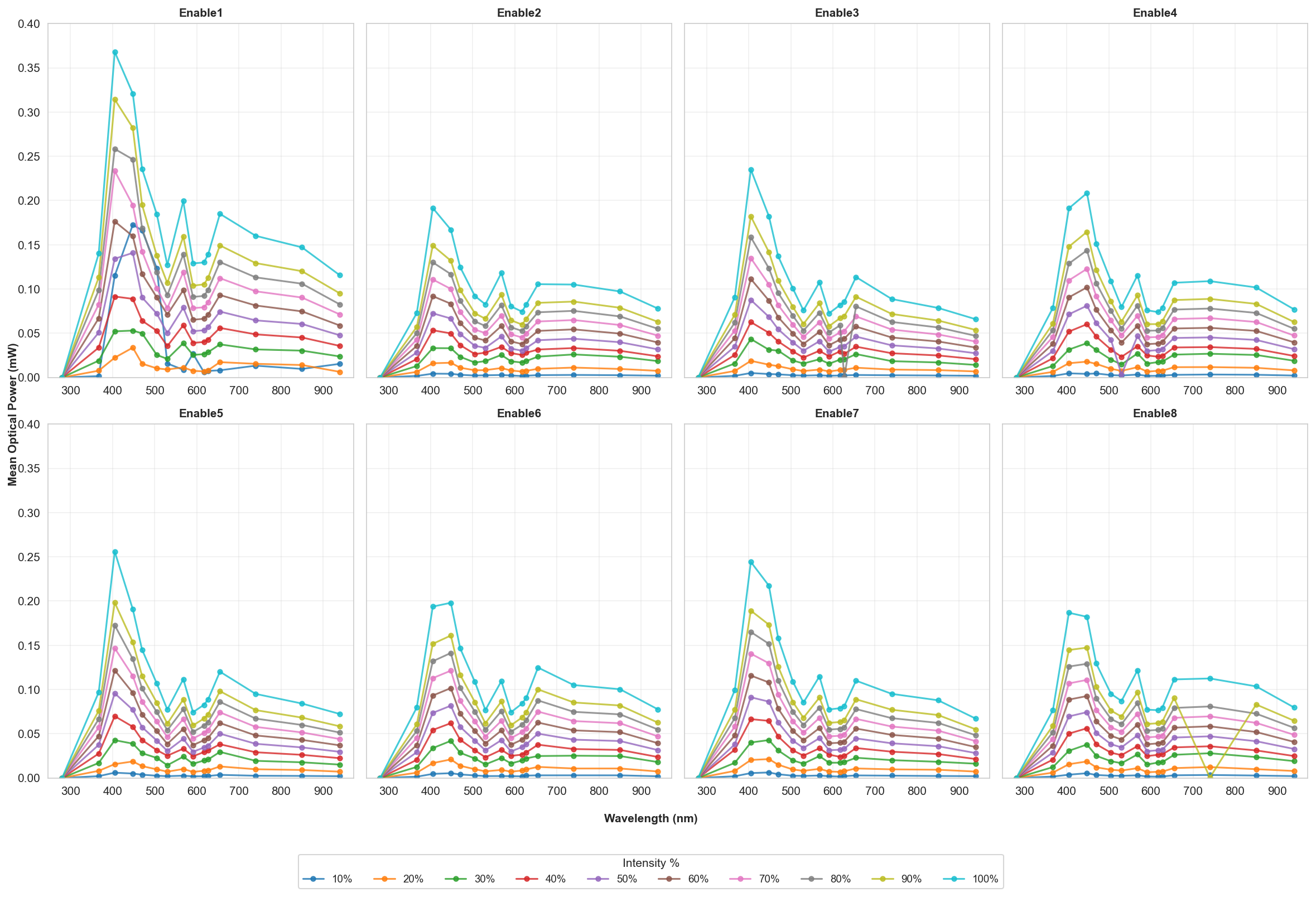

Figure S7. Spectral Power Response of Individual Optical Enable Zones. Optical power output across the full wavelength range (367-940 nm) for each of the eight individual LED enable zones (Enable1-Enable8). Mean power (±SD from n=5 replicates) is plotted against wavelength at ten intensity levels (10%-100%), shown as distinct curves per subplot. Power was measured in µW across the full intensity range (10-100%). Most wavelengths exhibited near-linear increases with drive intensity. These data confirm scalable radiant flux control across the wavelength range and enable zones for quantitative imaging experiments.

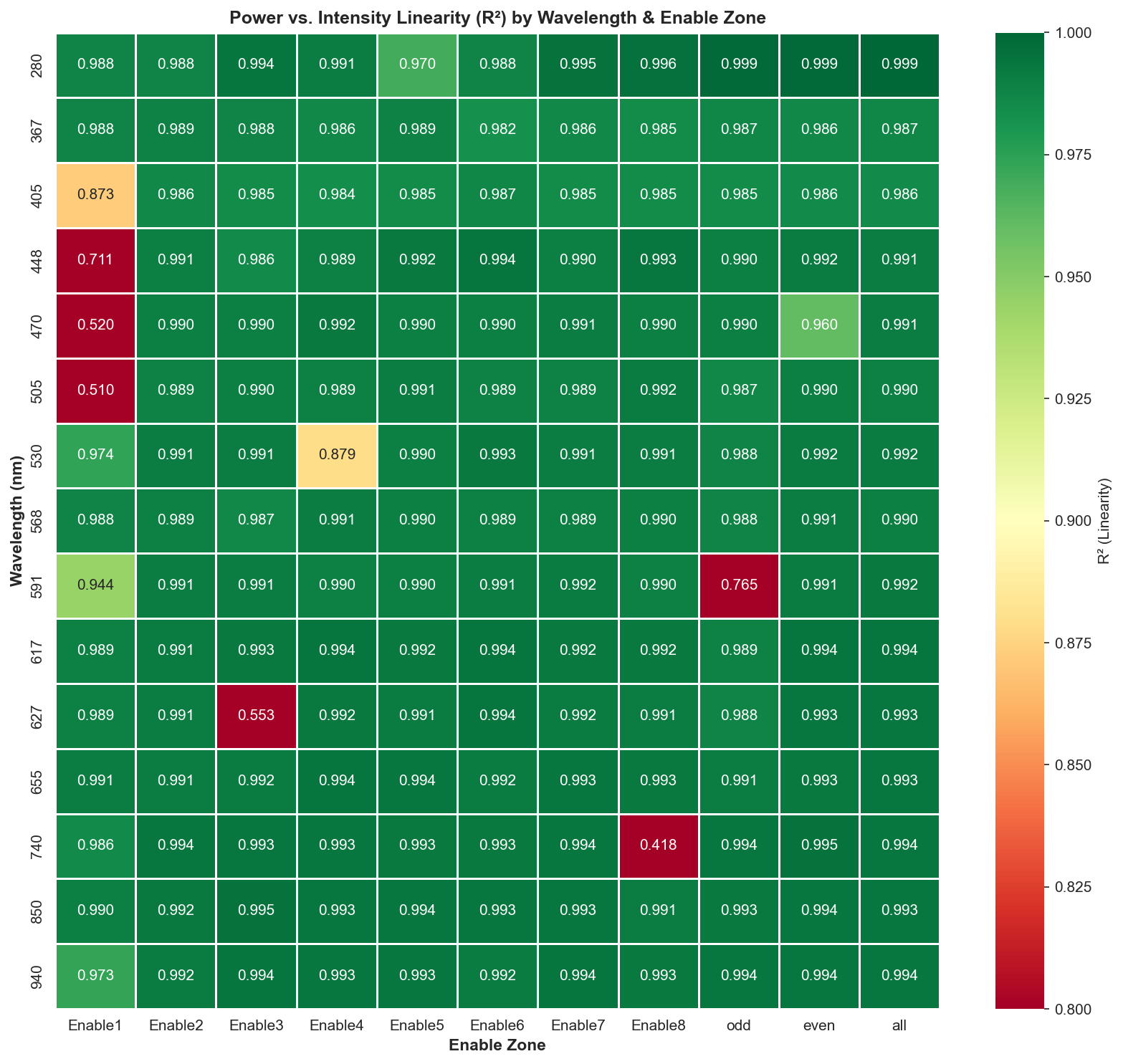

Figure S8. Linearity assessment of optical power response across wavelengths and enable zones. Heatmap of coefficient of determination (R²) values quantifying the goodness-of-fit between measured optical power and driver intensity (10%-100% PWM) for each wavelength-enable zone combination. Colors indicate R² values (green: R² > 0.95, excellent linearity; yellow: 0.85-0.95, acceptable; red: R² < 0.85, poor linearity). Mean R² = 0.974 ± 0.081 across 120 wavelength-zone pairs. Low R² regions (marked red) indicate nonlinear response, saturation, or incomplete measurement sweeps and excluded.

### Mineral Fluorescence Fingerprinting

Fifteen mineral specimens were selected to span the range of fluorescence intensity and peak position commonly reported for UV-excited fluorescent minerals (Frey Scientific and Amazon). Three calcite specimens from different sources were included as independent classes to assess the ability of the platform to distinguish nominally identical mineral types.

Images were acquired for each excitation wavelength and long-pass emission filter over multiple exposure times (50-300 ms) and camera gains (100-300). Three fixed ROIs were analyzed for each specimen. Mean ROI intensity was normalized by exposure time and gain to permit comparison across acquisition settings. Unsaturated measurements were combined using the median response for each ROI. Normalized spectral feature vectors were analyzed using cosine similarity, principal component analysis (PCA), hierarchical clustering, and silhouette analysis to assess spectral reproducibility and class separability.

Excitation-dependent fluorescence fingerprints are summarized in Figures S9-S12. The 367 and 405 nm excitation wavelengths were selected because they span the principal near-UV excitation range for fluorescent minerals while providing sufficient optical power and detector sensitivity for quantitative imaging. Combining 367 and 405 nm excitation improved class separability relative to either wavelength alone (Figure S9). Several minerals exhibited distinct excitation-dependent emission profiles, whereas others retained similar spectral shapes across both excitation wavelengths (Figures S10-S12).

Table S1. Representative fluorescence intensities of mineral specimens. Mean ROI intensity (mean ± SD) measured under representative excitation and acquisition conditions.

| Mineral | 367 nm, 50 ms, Gain 100 (mean ± std) | 367 nm, 50 ms, Gain 300 (mean ± std) | 405 nm, 50 ms, Gain 100 (mean ± std) | 405 nm, 50 ms, Gain 300 (mean ± std) |
| --- | --- | --- | --- | --- |
| Hackmanite | 9.84 ± 5.08 | 97.9 ± 41.5 | 16.3 ± 9.28 | 147 ± 63.5 |
| Apatite | 1.64 ± 2.99 | 14.7 ± 14.5 | 3.61 ± 7.11 | 56.4 ± 84.1 |
| Fluorite | 1.60 ± 2.95 | 13.4 ± 13.1 | 3.15 ± 7.95 | 43.0 ± 79.5 |
| Selenite | 1.66 ± 4.31 | 12.6 ± 23.8 | 2.10 ± 5.64 | 33.3 ± 60.6 |
| Chalcedony | 1.55 ± 3.81 | 12.7 ± 19.2 | 2.93 ± 6.93 | 40.6 ± 71.9 |
| Willemite | 0.66 ± 2.22 | 2.62 ± 4.43 | 0.81 ± 2.54 | 9.30 ± 17.0 |
| Tremolite | 0.64 ± 2.22 | 3.06 ± 4.31 | 0.80 ± 2.49 | 11.7 ± 19.5 |
| Resinous Coal | 0.66 ± 2.22 | 2.51 ± 4.36 | 0.80 ± 2.56 | 10.3 ± 17.4 |

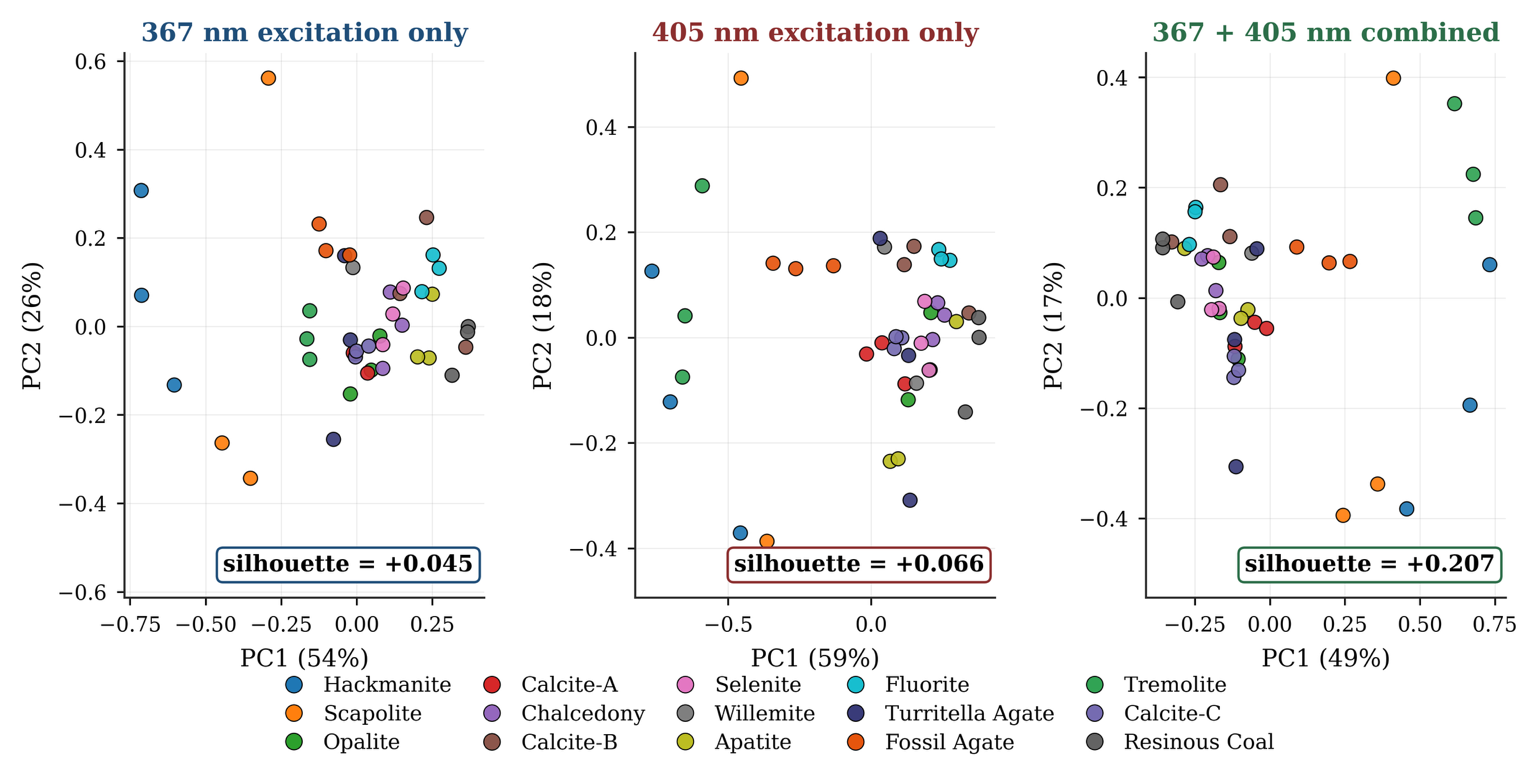

Figure S9**.** Effect of excitation wavelength on mineral fingerprint separability. PCA projections of feature vectors acquired under 367 nm excitation (left), 405 nm excitation (center), and the combined excitation dataset (right). Silhouette scores computed from the full feature space are indicated for each analysis.

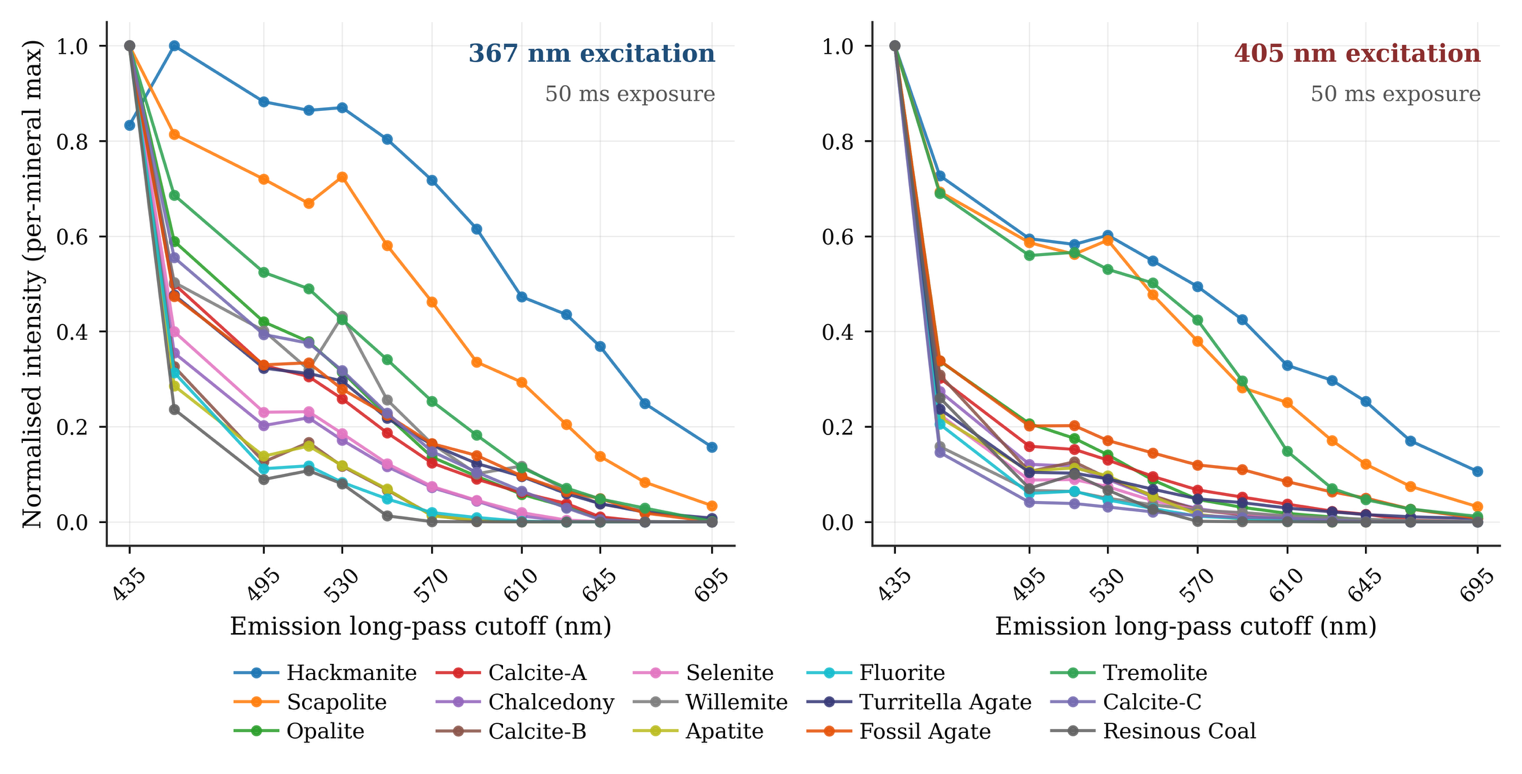

Figure S10. Normalized fluorescence emission fingerprints of mineral specimens. Normalized emission intensity measured through the long-pass filter series under 367 nm (left) and 405 nm (right) excitation for 15 mineral specimens. Each curve represents the normalized median emission measured across 2-3 ROIs for each mineral under the indicated excitation wavelength.

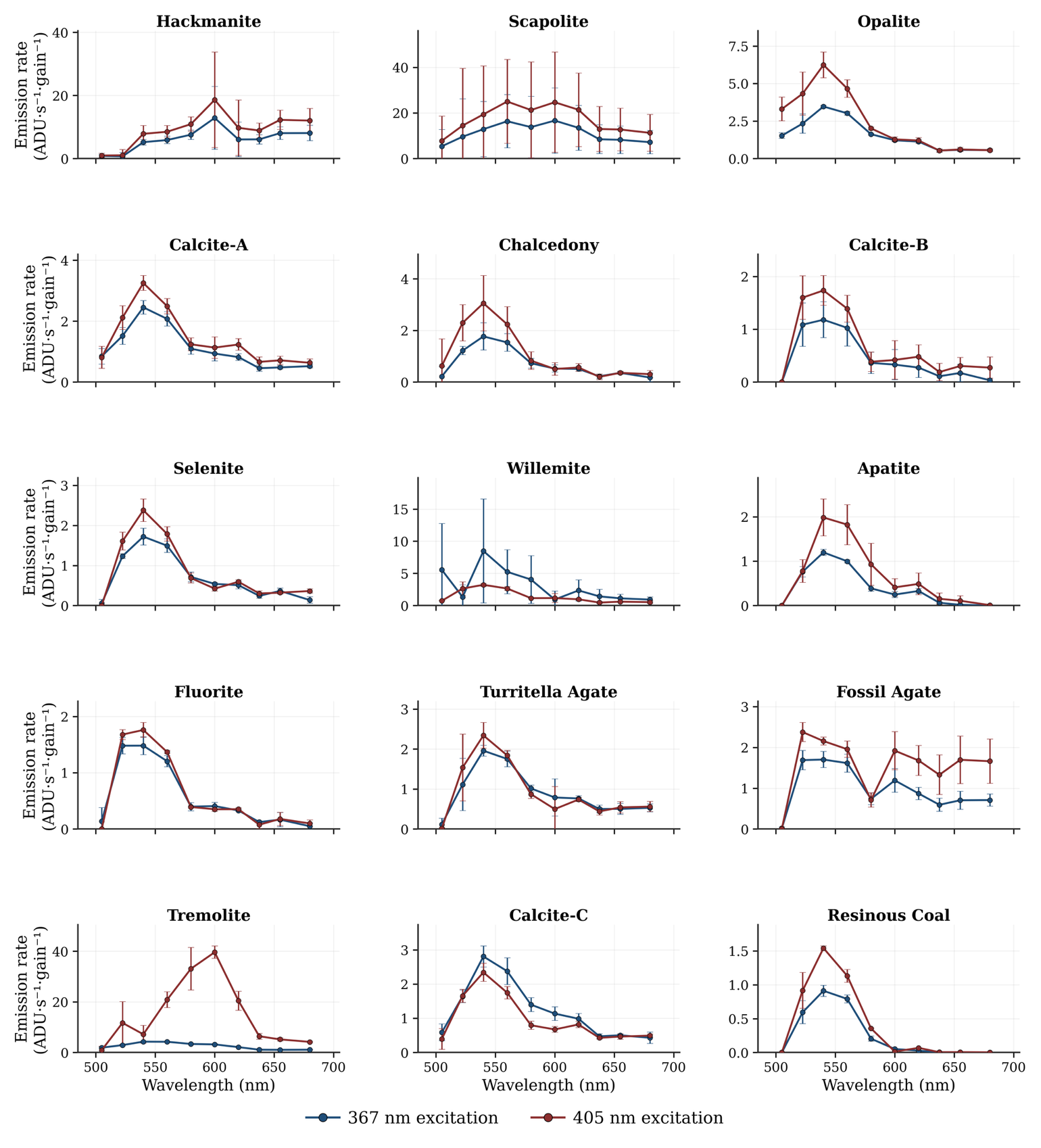

Figure S11. Per-sample band-differenced emission spectra for 15 mineral samples. Each panel shows mean emission rate at 367 nm (blue) and 405 nm (red) excitation across mineral samples obtained by successive differencing of the long-pass filter stack. Since each panel is independently y-axis-scaled to reveal spectral shape, absolute emission rates are not directly comparable between panels and readers should compare spectral shape. Several minerals exhibit strongly excitation-dependent emission: Tremolite shows a 405 nm-excited peak at ~590-600 nm that is nearly absent at 367 nm; Willemite shows the inverse pattern with a 367 nm-excited peak near 540 nm. The three Calcite specimens (A, B, C) share a primary emission peak at 515-550 nm but differ in long-wavelength tail behavior.

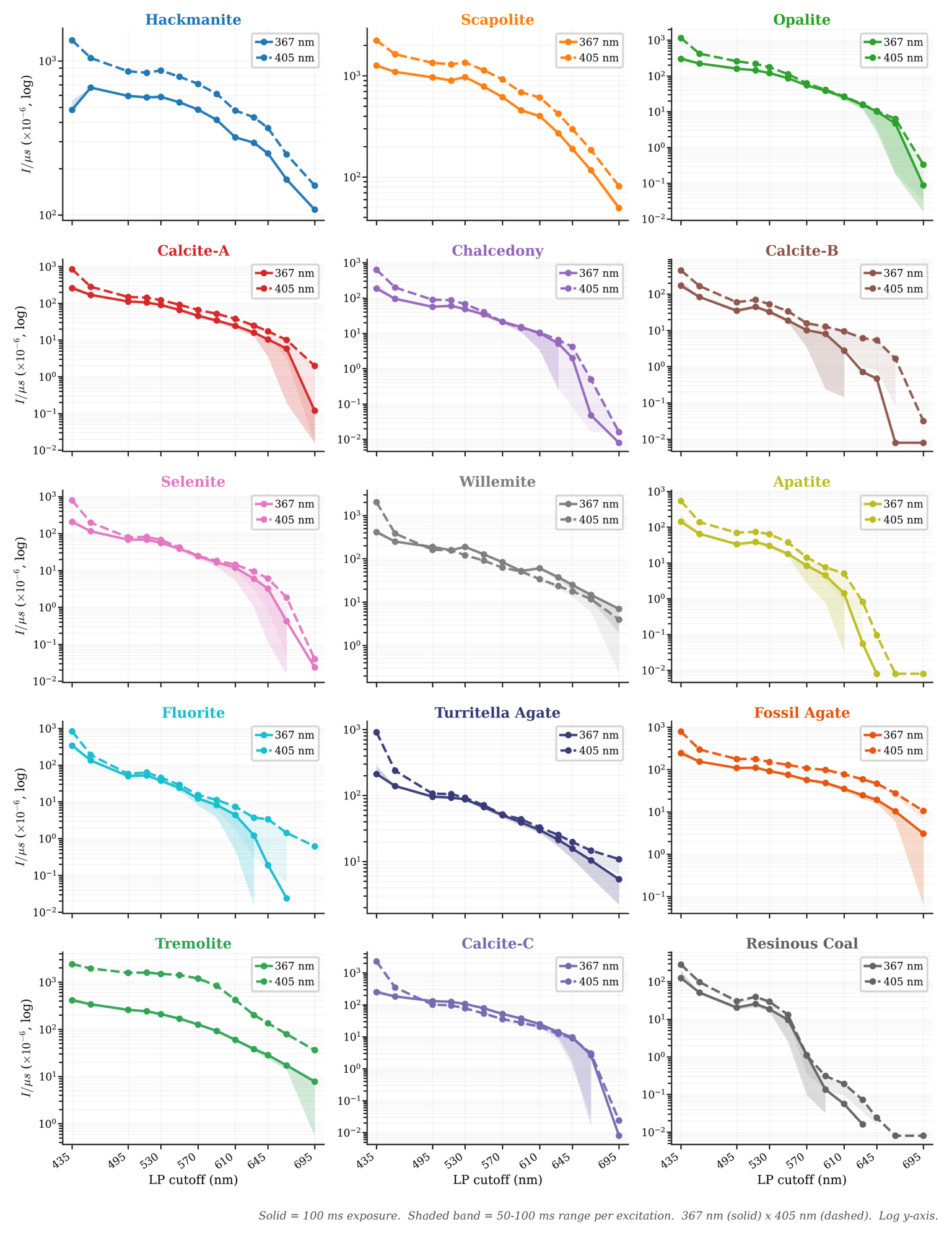

Figure S12. Excitation-dependent emission spectral signatures for all 15 mineral specimens. Emission intensity normalized per microsecond of integration time measured across the long-pass filter series under 367 nm (solid) and 405 nm (dashed) excitation. The solid marker trace for each excitation wavelength corresponds to the spectrum at 100 ms exposure; the shaded band spans the range of normalized values obtained at 50 ms and 100 ms exposure for that excitation channel, illustrating the consistency of the normalized signal across this portion of the exposure sweep described in the Methods.

### Time-Resolved Phosphorescence Decay Imaging

Six commercial phosphor powders (TechnoGlow Professional Sample Pack) were excited with 405 nm illumination for 300 s under identical conditions. Following excitation, phosphorescence images were acquired at 1 frame s⁻¹ for 1,980 s using a thermoelectrically cooled monochrome camera operated at −10 °C and gain 100.

At each frame, the background level was estimated as the pooled mean of six background ROIs of identical dimensions, sampled from phosphor-free regions of the sensor. This background estimate was subtracted from the ROI mean to obtain the background-corrected signal. The signal for each phosphor was computed independently at each timepoint from the spatial statistics of pixel values from three replicate regions of interest (ROI). Decay curves were fit using mono-exponential, bi-exponential, tri-exponential, stretched exponential (KWW), power-law, log-normal, gamma-distribution, and Tsallis-q models. Model performance was evaluated using the coefficient of determination (R²) and the Akaike Information Criterion (AIC). Residual analysis and temporal signal-to-noise ratio were used to assess fit quality and measurement stability.

Representative phosphorescence decay curves and corresponding tri-exponential fits are shown in Figures S13-S15. Comparison of candidate decay models demonstrates that multi-parameter and continuous-distribution models provide substantially better agreement with the measured decays than mono- or bi-exponential models (Table S2; Figures S16-S18). Signal-to-noise ratio remained sufficient for quantitative analysis throughout most of the acquisition period for the longer-lived phosphors (Figure S19).

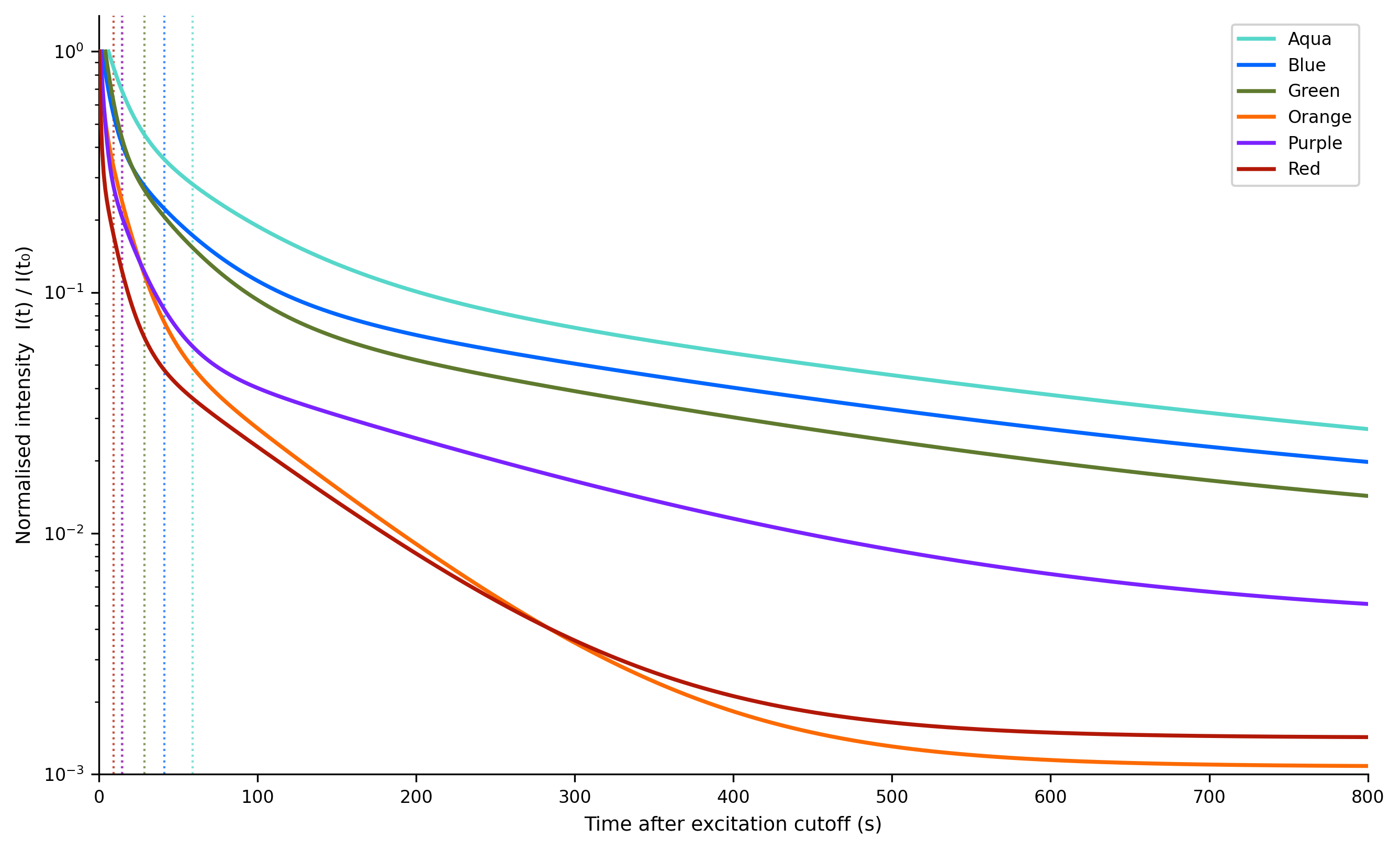

Figure S13. Normalized tri-exponential phosphorescence decay curves for six phosphor samples. Decay curves are shown as normalized intensity, I(t)/I(t₀), on a semi-logarithmic axis, where I(t) is the tri-exponential fit to the mean ROI decay signal derived from each phosphor and I(t₀) is the fitted intensity at the first valid time point after excitation cutoff. Normalization isolates the shape of the decay kinetics from differences in absolute phosphorescence intensity across samples. Each phosphor (Aqua, Blue, Green, Orange, Purple, Red) is plotted as a solid line in its corresponding color for t ≤ 800 s after excitation cutoff. Dotted vertical lines indicate the amplitude-weighted mean lifetime, τ_mean_, for each phosphor, calculated from the time constants (τ₁, τ₂, τ₃) and their corresponding amplitudes (A₁, A₂, A₃) of the tri-exponential fit

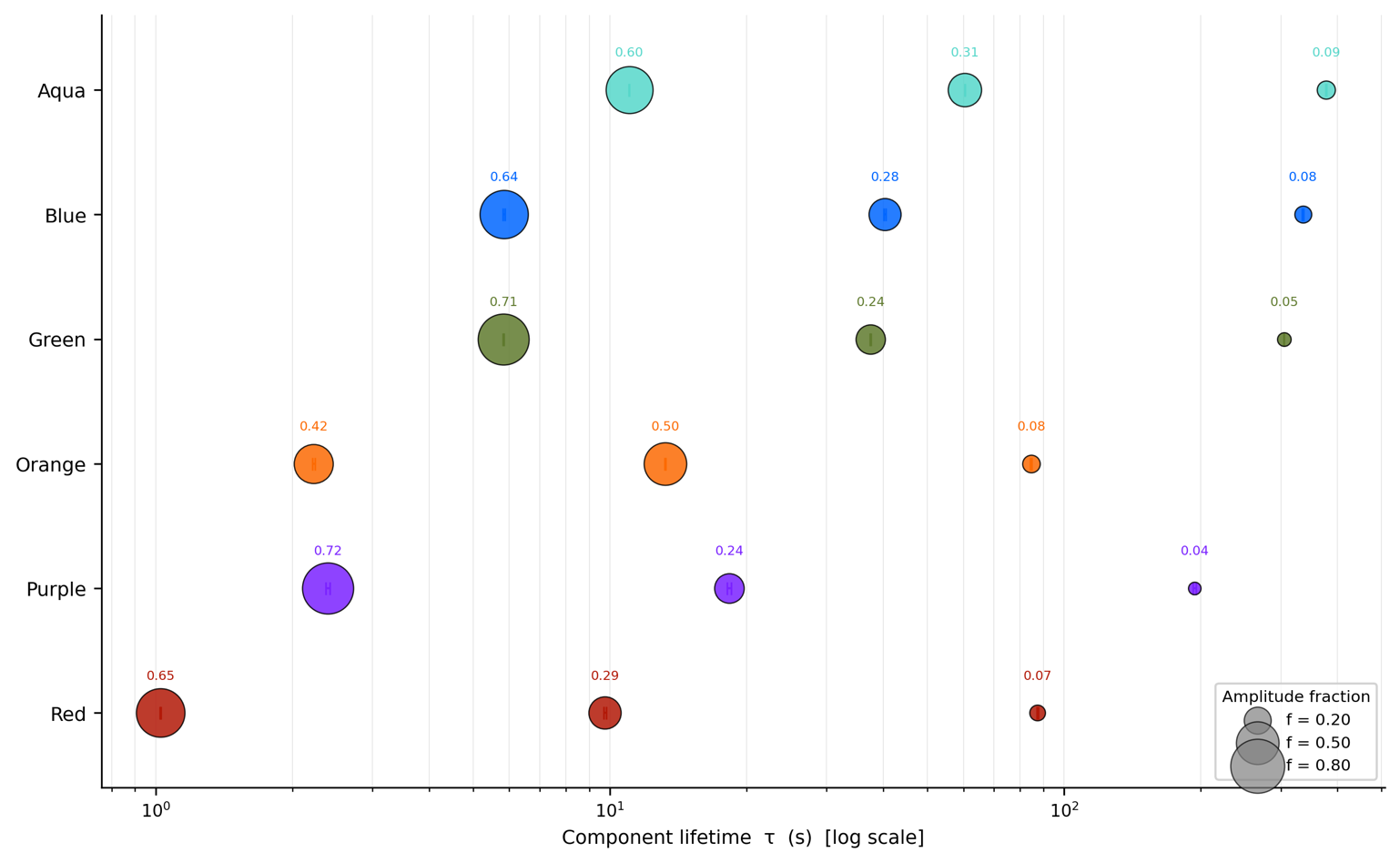

Figure S14**.** Tri-exponential lifetime components. Lifetime measurements for six phosphor samples. Bubble position indicates decay lifetime, and bubble area is proportional to the corresponding amplitude fraction.

Table S2. Tri-exponential decay parameters by phosphor channel Tri-exponential decay parameters extracted from phosphorescence lifetime measurements for the six phosphor channels shown in Figure S14. Amplitude fractions correspond to the bubble diameters in Figure S14, and component lifetimes correspond to bubble positions along the log-scaled abscissa. Reported uncertainties are 1σ parameter standard errors from the nonlinear least-squares fit. τ_amp_ is the amplitude-weighted mean lifetime and t_1/2_ is the corresponding half-life.

| Phosphor | τ₁ (s) | f₁ | τ₂ (s) | f₂ | τ₃ (s) | f₃ | R² | ⟨τ⟩ₐₘₚ (s) | t₁/₂ (s) |
| --- | --- | --- | --- | --- | --- | --- | --- | --- | --- |
| Red | 1.02 ± 0.01 | 0.649 | 9.66 ± 0.09 | 0.284 | 86.9 ± 0.9 | 0.067 | 0.9985 | 9.20 | 1.28 |
| Orange | 2.21 ± 0.02 | 0.419 | 13.22 ± 0.09 | 0.496 | 84.5 ± 0.8 | 0.085 | 0.9994 | 14.66 | 4.28 |
| Aqua | 11.02 ± 0.07 | 0.604 | 60.58 ± 0.38 | 0.305 | 378.3 ± 2.3 | 0.091 | 0.9997 | 59.53 | 13.96 |
| Green | 5.85 ± 0.04 | 0.712 | 37.61 ± 0.28 | 0.237 | 305.4 ± 2.5 | 0.051 | 0.9993 | 28.66 | 6.15 |
| Blue | 5.88 ± 0.04 | 0.639 | 40.51 ± 0.32 | 0.283 | 336.7 ± 2.6 | 0.079 | 0.9992 | 41.72 | 7.24 |
| Purple | 2.42 ± 0.02 | 0.715 | 18.55 ± 0.16 | 0.242 | 195.4 ± 2.0 | 0.044 | 0.9985 | 14.76 | 2.58 |

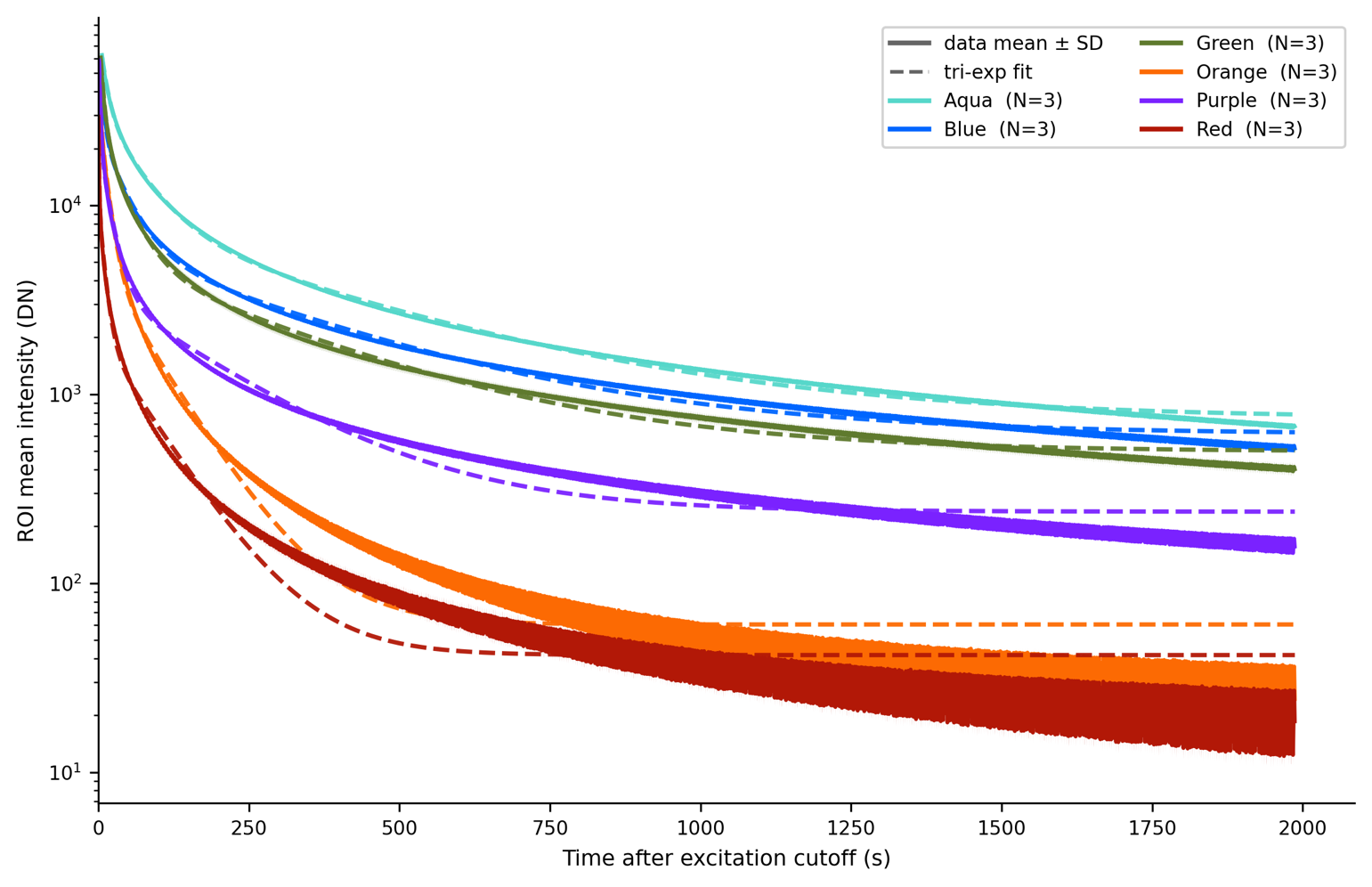

Figure S15. Phosphorescence decay curves. Mean phosphorescence intensity (solid lines; mean ± SD, n = 3) and corresponding tri-exponential fits (dashed lines) measured over the 2000 s acquisition period.

Table S3. Comparison of phosphorescence decay models. Model fit statistics, coefficient of determination (R²), Akaike Information Criterion (AIC), for the different decay models.

| Phosphor | R² / ΔAIC | | | | |
| --- | --- | --- | --- | --- | --- |
|  | Bi-exp | Tri-exp | Power law | KWW | Log-normal |
| Aqua | 0.9957 / 10279 | 0.9997 / 4816 | 0.9994 / 6314 | 0.9996 / 5542 | 0.99998 / 0 |
| Blue | 0.9906 / 6920 | 0.9992 / 2068 | 0.9993 / 1822 | 0.9969 / 4679 | 0.99971 / 0 |
| Green | 0.9917 / 8147 | 0.9993 / 3242 | 0.99986 / 0 | 0.9924 / 7961 | 0.99882 / 4273 |
| Orange | 0.9951 / 5839 | 0.9994 / 1642 | 0.9986 / 3319 | 0.9960 / 5406 | 0.99974 / 0 |
| Purple | 0.9865 / 8283 | 0.9985 / 3955 | 0.99979 / 0 | 0.9857 / 8398 | 0.99748 / 4947 |
| Red | 0.9869 / 5863 | 0.9985 / 1572 | 0.9944 / 4156 | 0.9940 / 4293 | 0.99931 / 0 |

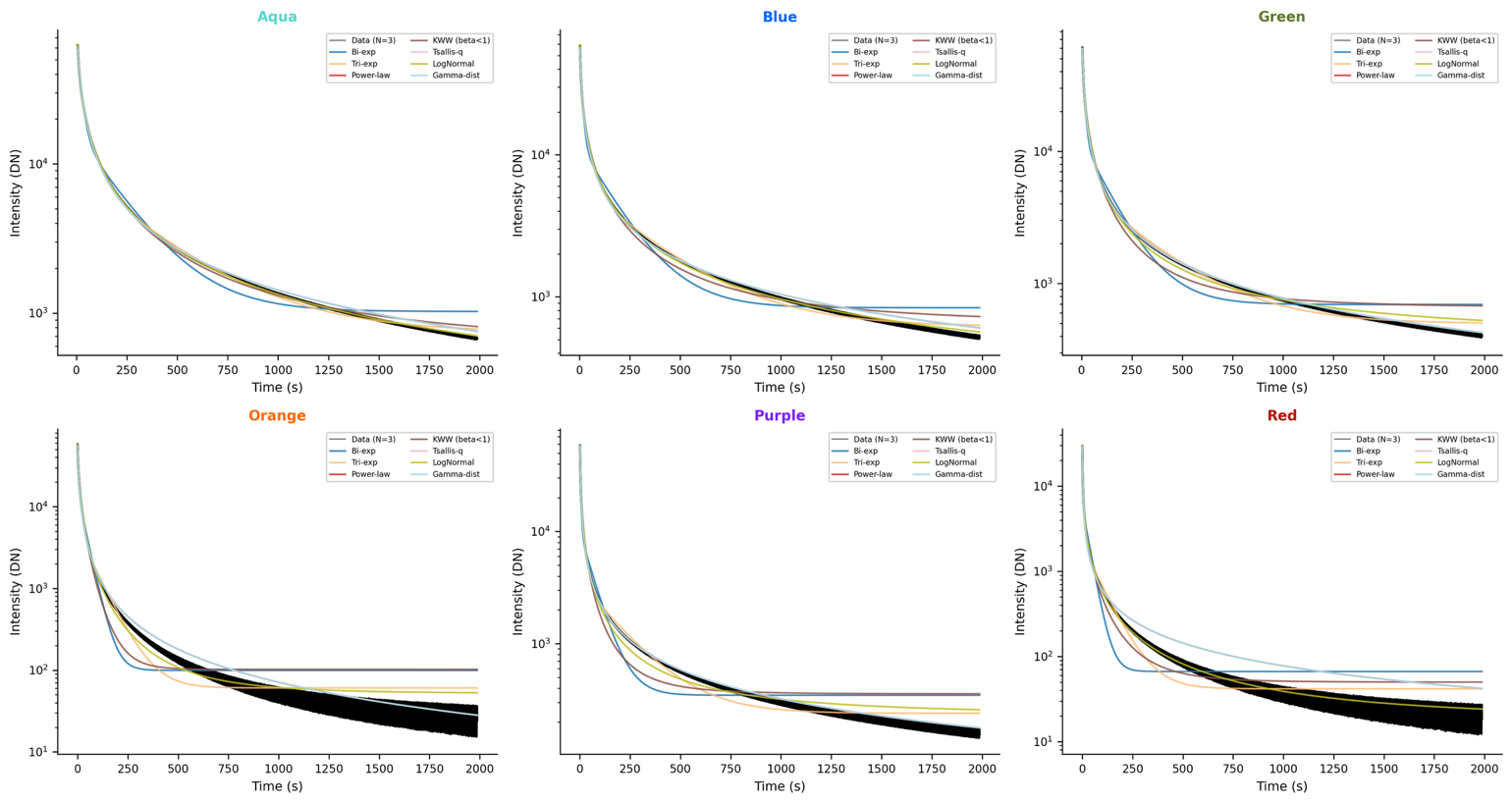

Figure S16. Comparison of phosphorescence decay models. Representative fits of the candidate decay models to each phosphor sample.

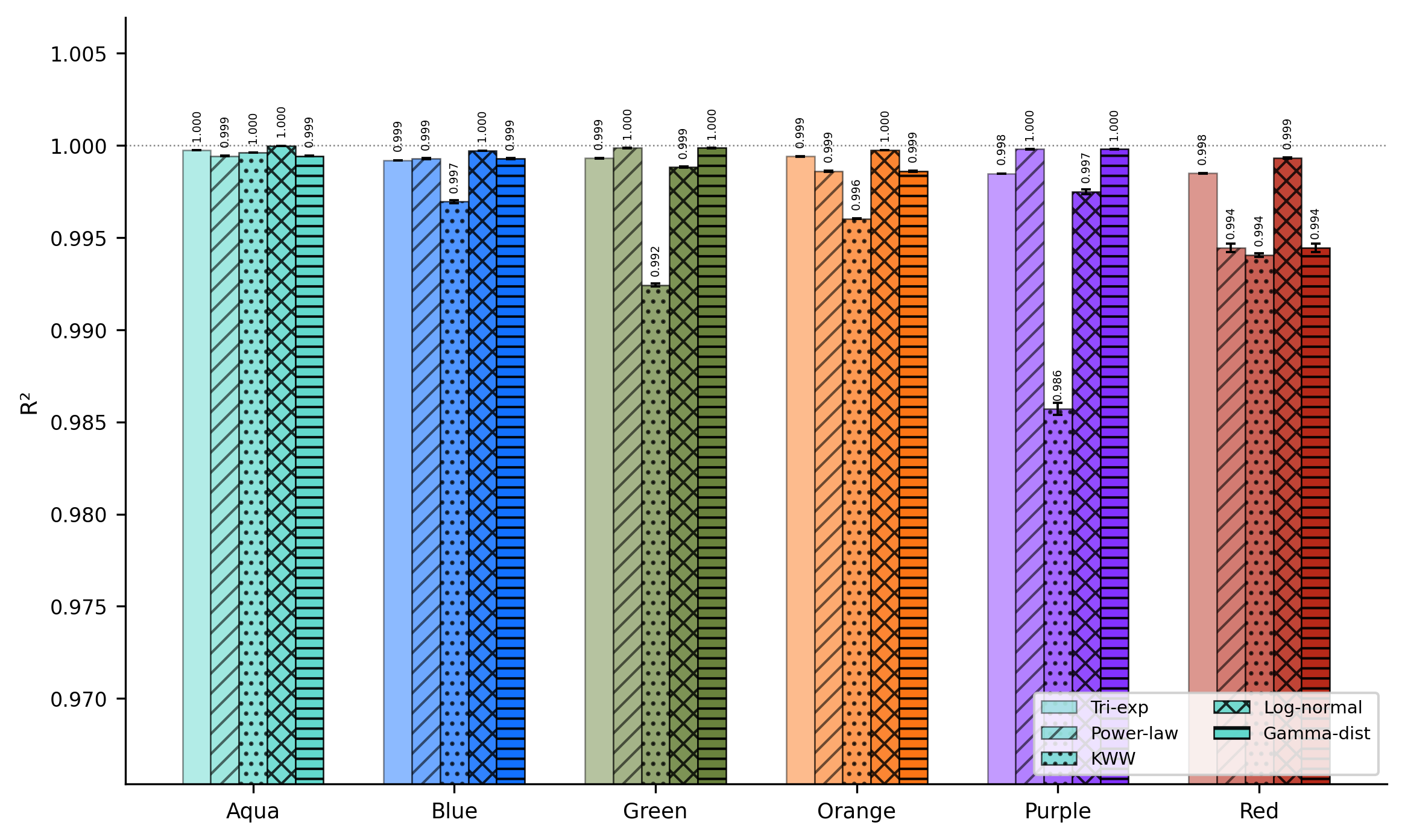

Figure S17. Goodness-of-fit comparison across decay models. Mean coefficient of determination (R²) for candidate phosphorescence decay models fitted to each phosphor sample. Error bars represent ±SD (n = 3).

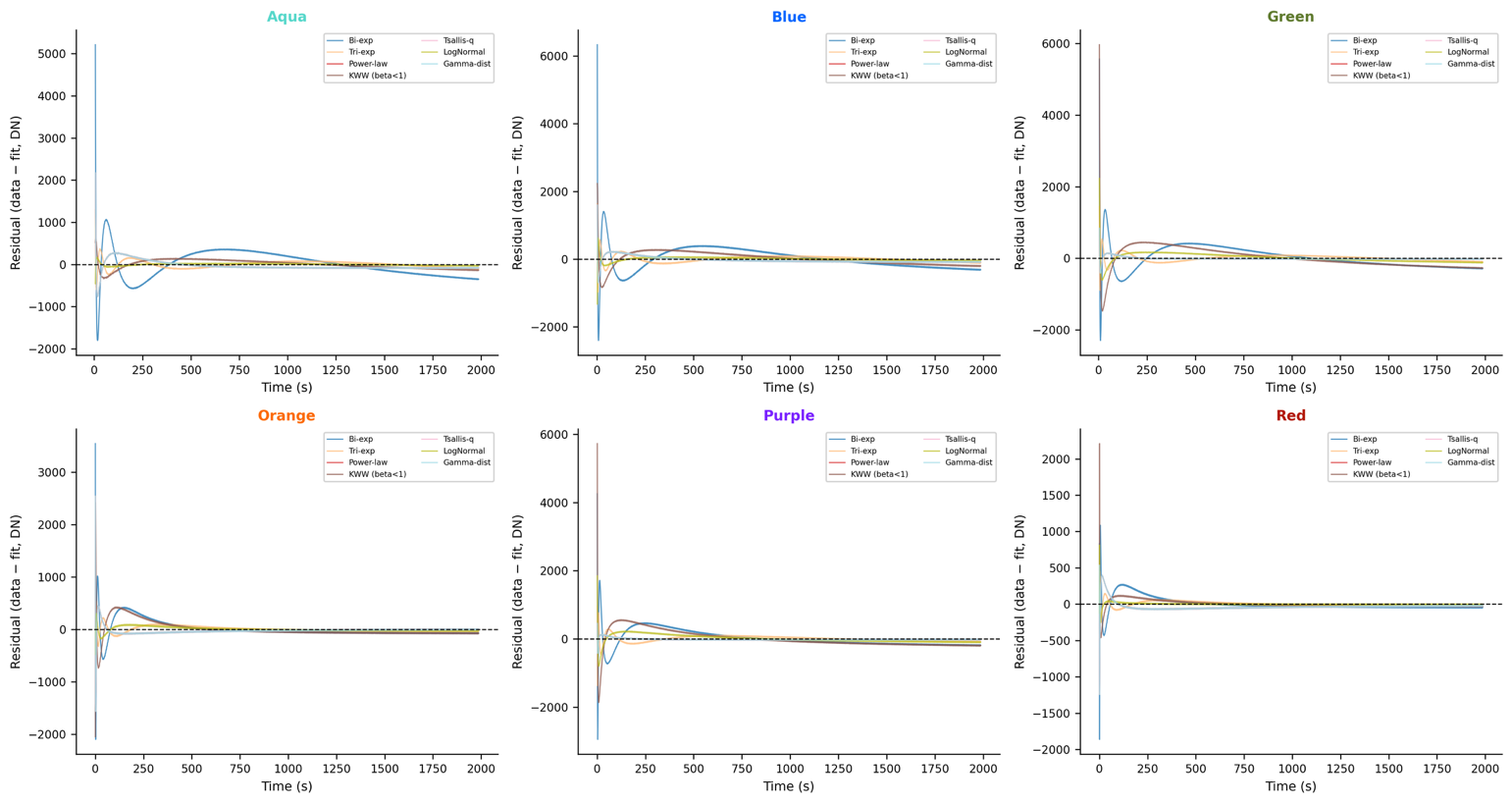

Figure S18. Residual analysis of phosphorescence decay models. Residuals (measured minus fitted intensity) for each decay model evaluated over the full acquisition period.

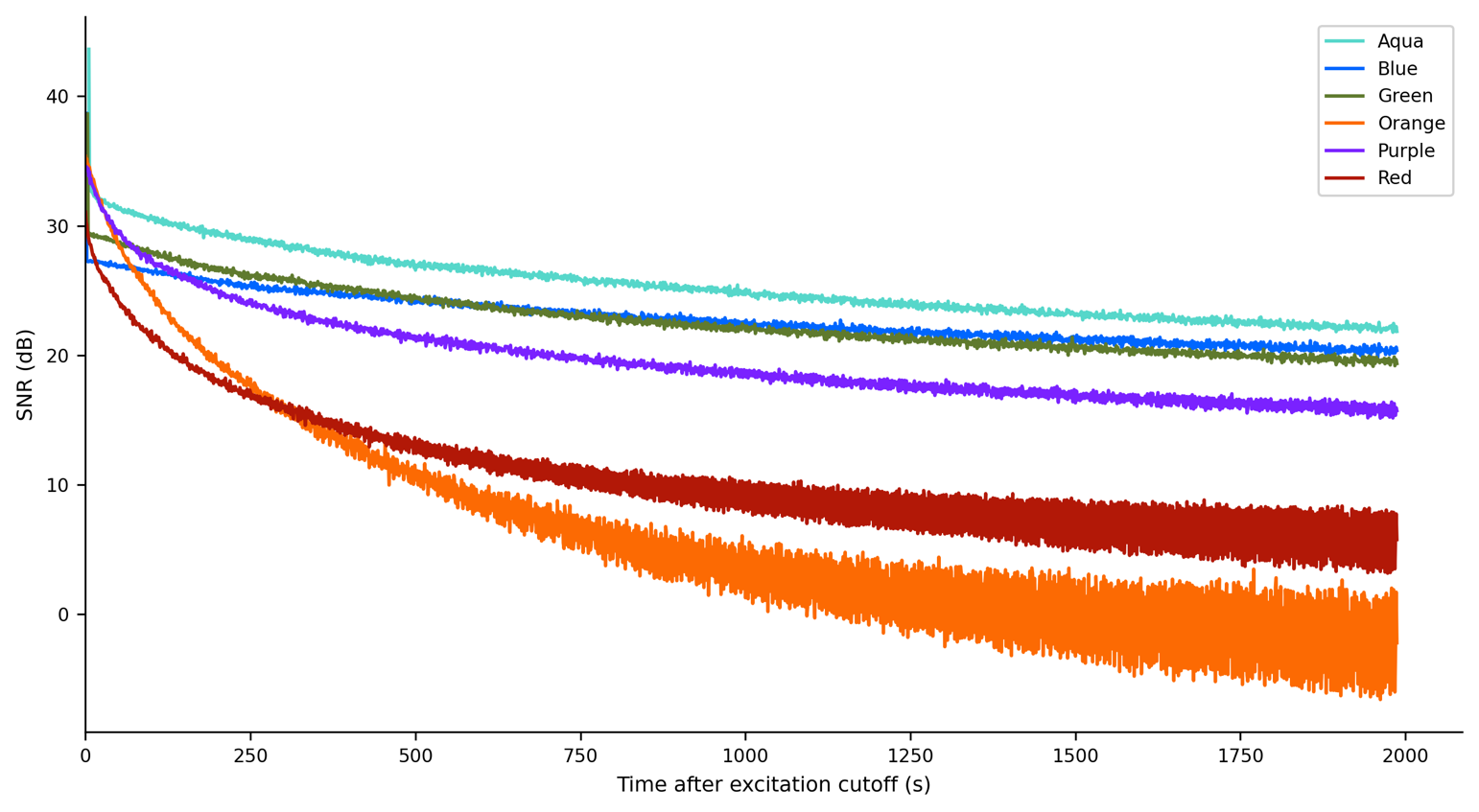

Figure S19. Temporal signal-to-noise ratio of phosphorescence measurements. Signal-to-noise ratio (SNR) was computed at each timepoint as the background-subtracted spatial mean divided by the spatial standard deviation within a ROI (N=3 per phosphor), rather than from a temporal window of adjacent frames. This background estimate was subtracted from the ROI mean to obtain the background-corrected signal, and the noise term was taken as the spatial sample standard deviation, of pixel values within the same signal ROI at the same timepoint.

### Bioluminescence Imaging

A XPM-2 calibrated bioluminescent phantom (Revvity) containing two independently addressable embedded light sources was imaged in both illumination modes and orientations. Reference images were acquired using a Revvity IVIS Spectrum under standard autoexposure conditions. Corresponding AURORA-MSI images were acquired in complete darkness at exposure times of 1, 10, and 30 s. Reference datasets were acquired using an IVIS Spectrum system at Anschutz Medical Center with manufacturer-recommended auto-exposure settings to generate standard pseudocolor overlays. Representative phantom and cell-line imaging results are shown in Figures S20 and S22. AURORA-MSI reproduced the spatial localization of the embedded phantom sources. In the cell dilution series, signal was detected in both replicate wells at the highest density and in one of two replicate wells at each of the two lower densities (Figure S22).

For each exposure condition, calibrated frames were bias-subtracted, dark-subtracted, and flat-fielded. A smooth background pedestal was estimated by Gaussian-blurring a median-filtered copy of each frame and subtracted to isolate the bioluminescent signal. The isolated signal was spatially smoothed for display, and a soft-edged alpha mask was generated from a percentile-based presence threshold. For pseudocolor rendering, the smoothed signal in each frame was independently rescaled by its own percentile range and mapped through a 16-node rainbow colormap, then alpha-composited over a dimmed grayscale reference. Because this normalization is performed independently per frame rather than against a common absolute scale, the resulting overlays are expected to appear similar in spatial extent and peak color across exposure times even where underlying photon counts and signal-to-noise ratio differ. Unlike the IVIS Spectrum reference, which reports the manufacturer's calibrated radiance (photons s⁻¹ cm⁻¹ sr⁻¹) under autoexposure, the AURORA-MSI overlays are uncalibrated, relative-intensity renderings intended to validate spatial source localization rather than absolute photon flux.

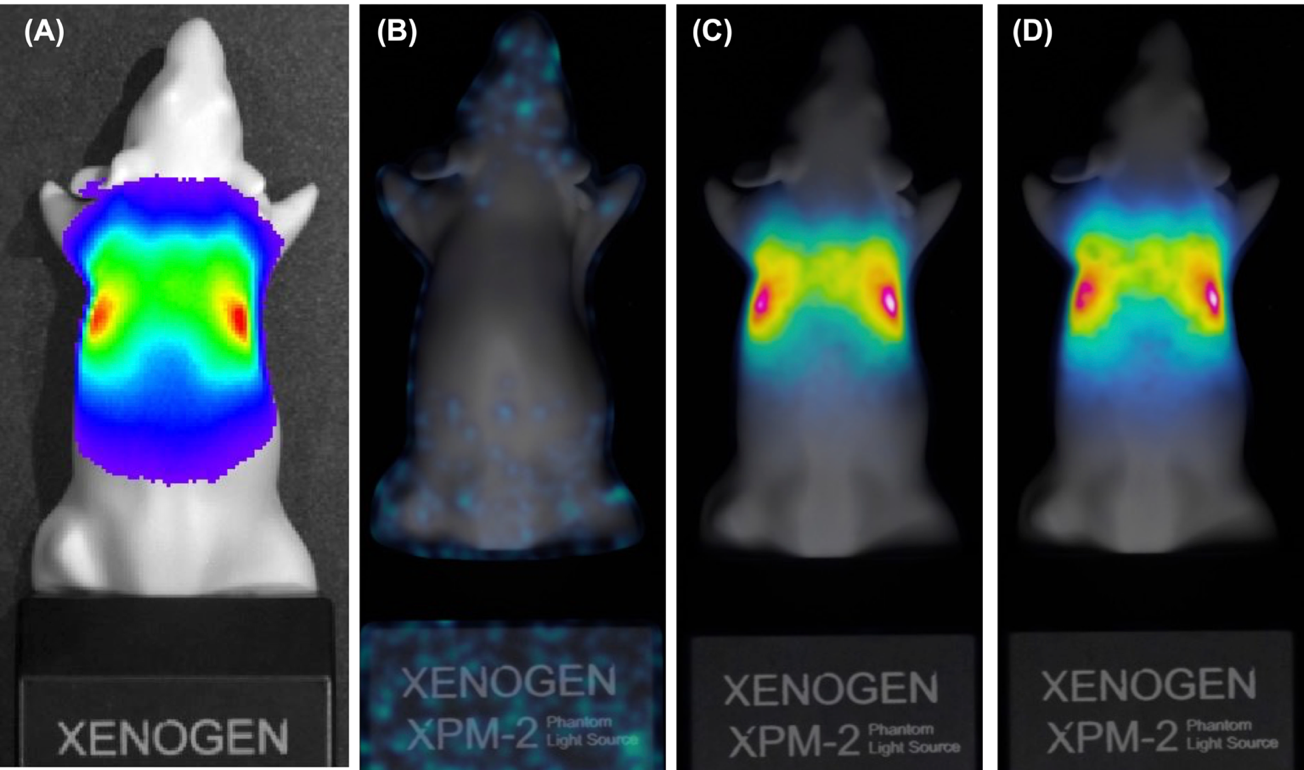

Figure S20. Imaging of the XPM-2 phantom with LED source “Mode B” active. (A) Reference image acquired with the Revvity IVIS Spectrum. (B-D) Corresponding AURORA-MSI images acquired with exposure times of 1, 10, and 30 s. For pseudocolor rendering, the smoothed signal in each frame was independently rescaled by its own percentile range and mapped through a 16-node rainbow colormap, then alpha-composited over a dimmed grayscale reference. Because this normalization is performed independently per frame rather than against a common absolute scale, the resulting overlays are expected to appear similar in spatial extent and peak color across exposure times. Panels B-D therefore illustrate spatial concordance with the reference, not a calibrated intensity comparison across exposures or against panel A.

| 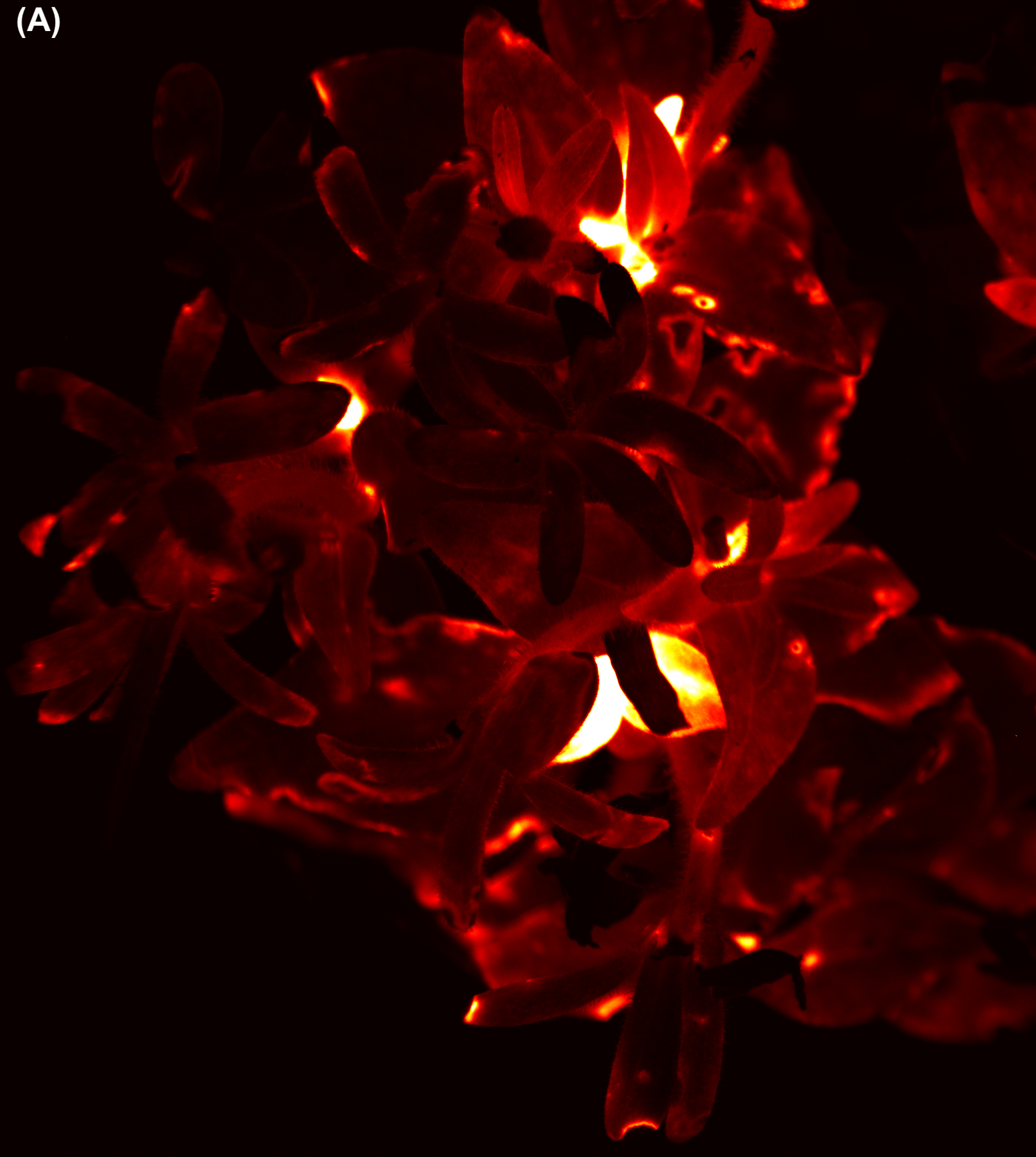 | 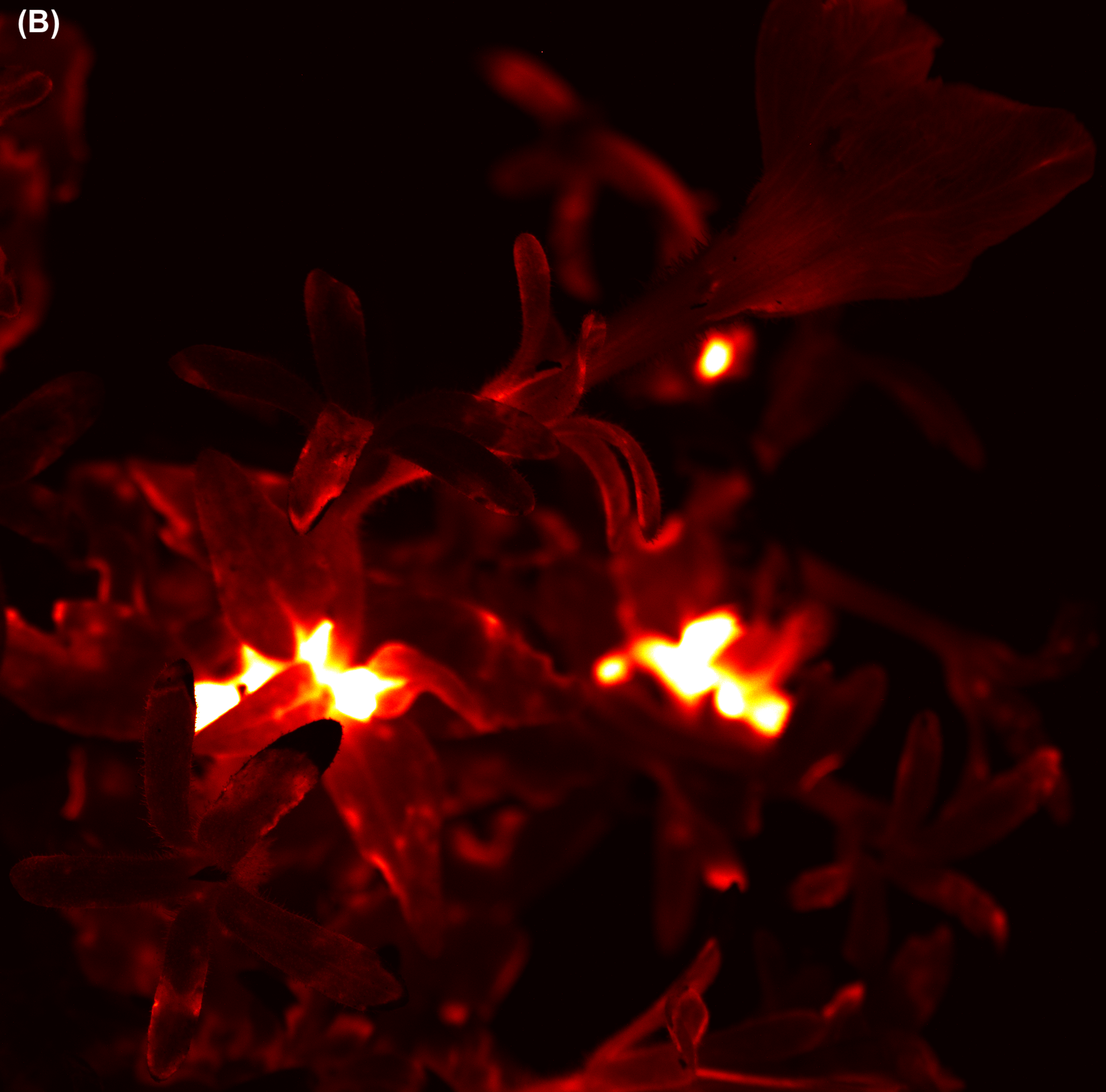 |
| --- | --- |

Figure S21. Full-resolution detail of the Firefly Petunia bioluminescence image (1200 s exposure) showing fine spatial structure. (A) Floral centers and petal margins exhibit elevated signal relative to petal interiors, consistent with the restriction of fungal bioluminescence pathway expression to actively metabolizing floral tissue described in the main text.

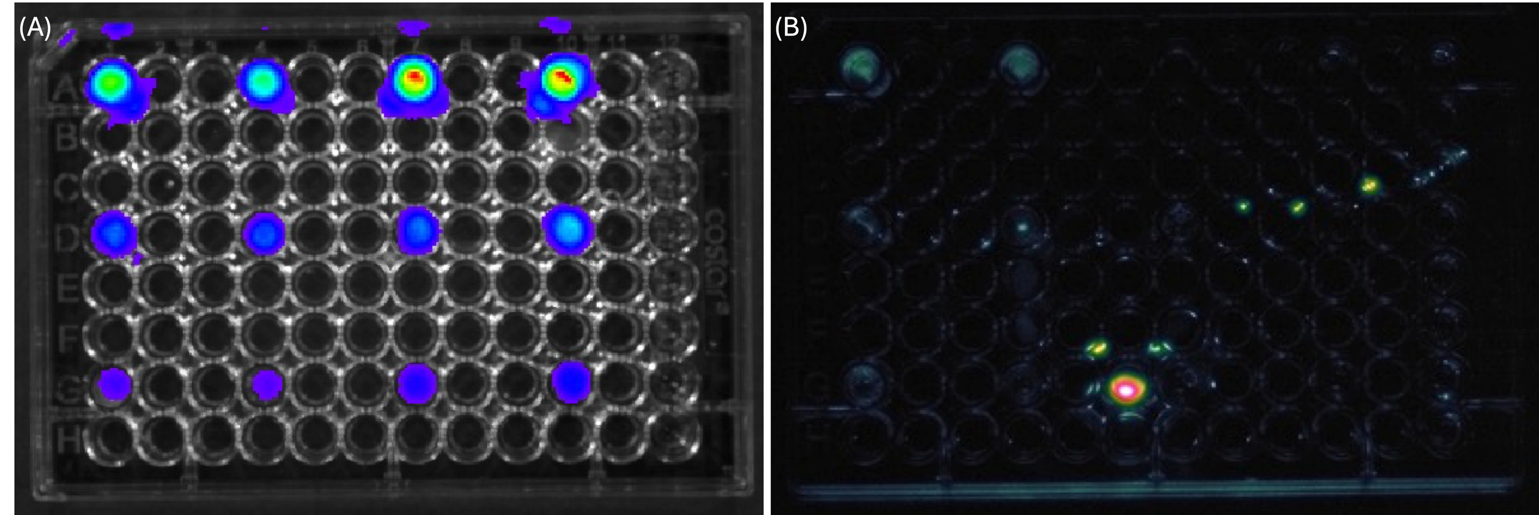

Figure S22. Bioluminescence imaging of a luciferase-expressing HSJD-GBM1-001 cell dilution series. (A) Reference image acquired with the Revvity IVIS Spectrum, following luciferin addition to a dedicated pair of replicate columns; all three cell-density tiers (50,000, 25,000, and 12,500 cells, top to bottom) are clearly resolved in both replicate wells. (B) Corresponding AURORA-MSI image of the same plate acquired using a separate pair of replicate columns dosed with luciferin prior to AURORA-MSI imaging (before the IVIS-session columns were dosed). The diagonal cluster of small spots in the upper right and the saturated high-intensity spot in the lower plate region do not correspond to expected well positions and are artifacts of undetermined origin rather than luciferase signal.

### Spectral Classification of Pharmaceutical and Chemically Similar Powders

Ten visually similar powders representing pharmaceutical, food, and household materials were loaded into individual wells of a standard 12-well plate and imaged under fixed geometry. Each well contained sufficient material to fully obscure the well bottom. Circular ROIs were automatically positioned within each well, excluding the well walls and specular reflections. Fifty images were acquired at each excitation wavelength. A sample size of fifty images per wavelength was selected to provide an adequate number of observations for both descriptive statistical characterization (mean, variance, texture features) and for constructing training/test splits for the classification analyses. Bootstrap resampling of the frame-level measurements shows that the relative standard error of ROI mean intensity decreases from 0.19% at N=5 to 0.06% at N=50, with diminishing marginal improvement beyond N≈30-35 frames. Critically, frame-to-frame measurement noise (mean CV = 0.43% across all powder-wavelength combinations) is more than two orders of magnitude smaller than between-powder variability (mean CV = 19.96%), confirming that N=50 provides measurement precision well below the discriminative signal exploited by the classification analyses.

For each ROI, first-order intensity statistics, gray-level co-occurrence matrix (GLCM) texture features, and local binary pattern (LBP) entropy were computed. Multispectral feature vectors were analyzed using principal component analysis (PCA), linear discriminant analysis (LDA), and Random Forest classification. Excitation-dependent intensity profiles for all powder classes are shown in Figure S23. Ultraviolet excitation produced the greatest variation among materials, whereas near-infrared excitation provided complementary reflectance contrast. Frame-to-frame repeatability is summarized in Figures S24 and S25. Measurement variability remained low across most powder-wavelength combinations. Higher-order intensity statistics are summarized in Figure S26 and contributed to the multispectral feature set used for classification. PCA and LDA projections are shown in Figures S27 and S28. Both analyses demonstrated clear separation of the powder classes in the multispectral feature space. Random Forest classification achieved complete separation of the powder classes (Figure S29). Feature importance rankings are summarized in Figure S30.

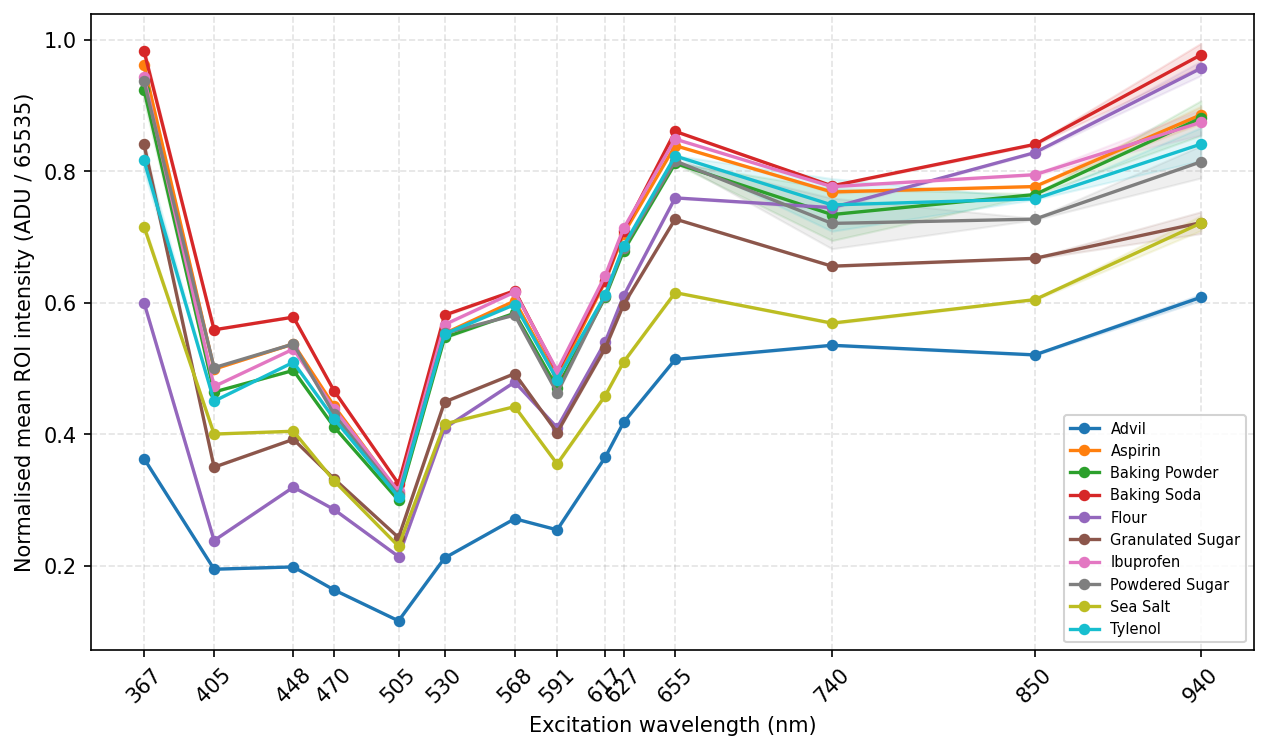

Figure S23. Excitation-dependent intensity fingerprints of powder samples. Mean ROI intensity as a function of excitation wavelength for each powder class. Curves are normalized to the maximum response of each material; shaded regions indicate ±SD (n = 50).

Table S4. Frame-level image descriptors for representative powder samples. Mean intensity, standard deviation, GLCM contrast, LBP entropy, and saturation fraction measured for each ROI.

| Powder | Mean | Std | Skew | Kurt | GLCM-C | LBP-H | Sat. frac. |
| --- | --- | --- | --- | --- | --- | --- | --- |
| Baking Soda | 239.4 | 18.7 | -1.29 | 1.07 | 0.91 | 2.73 | 0.20 |
| Ibuprofen | 224.3 | 35.4 | -1.43 | 2.25 | 0.79 | 3.42 | 0.19 |
| Powdered Sugar | 209.4 | 26.4 | -0.55 | 0.43 | 0.70 | 3.65 | 0.00 |
| Baking Powder | 198.9 | 29.5 | -2.10 | 6.76 | 0.49 | 3.01 | 0.02 |
| Granulated Sugar | 193.2 | 25.4 | -0.93 | 1.24 | 0.50 | 3.22 | 0.00 |
| Tylenol | 171.2 | 35.1 | -1.43 | 2.52 | 0.40 | 3.58 | 0.00 |
| Aspirin | 161.6 | 106.3 | -0.51 | -1.55 | 0.58 | 3.48 | 0.24 |
| Sea Salt | 109.6 | 79.0 | -0.23 | -1.60 | 0.31 | 3.73 | 0.00 |
| Flour | 108.9 | 64.8 | -0.49 | -0.90 | 0.40 | 3.58 | 0.00 |
| Advil | 54.7 | 46.0 | 0.28 | -1.42 | 0.17 | 3.98 | 0.00 |

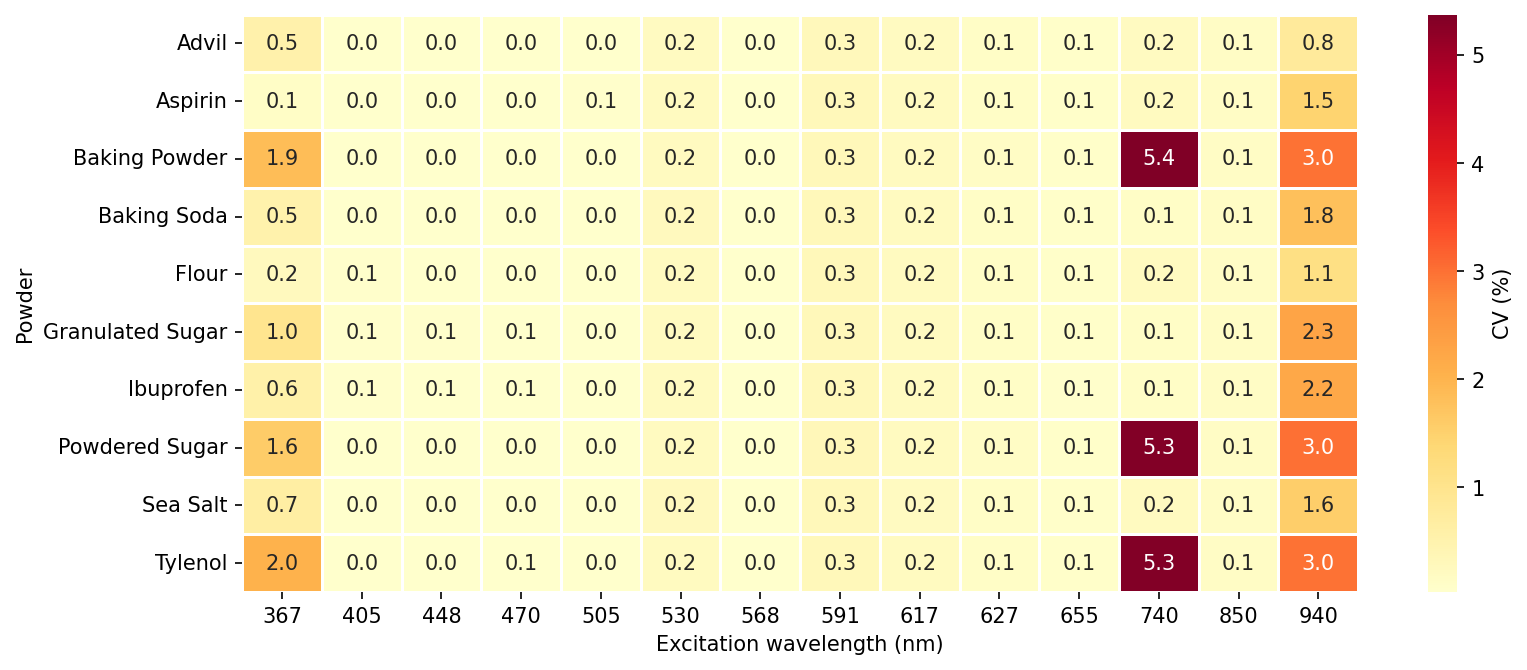

Figure S24. Repeatability of powder measurements across excitation wavelengths. Heatmap of the frame-to-frame coefficient of variation (CV, %, n = 50) of mean ROI intensity for each powder and excitation wavelength.

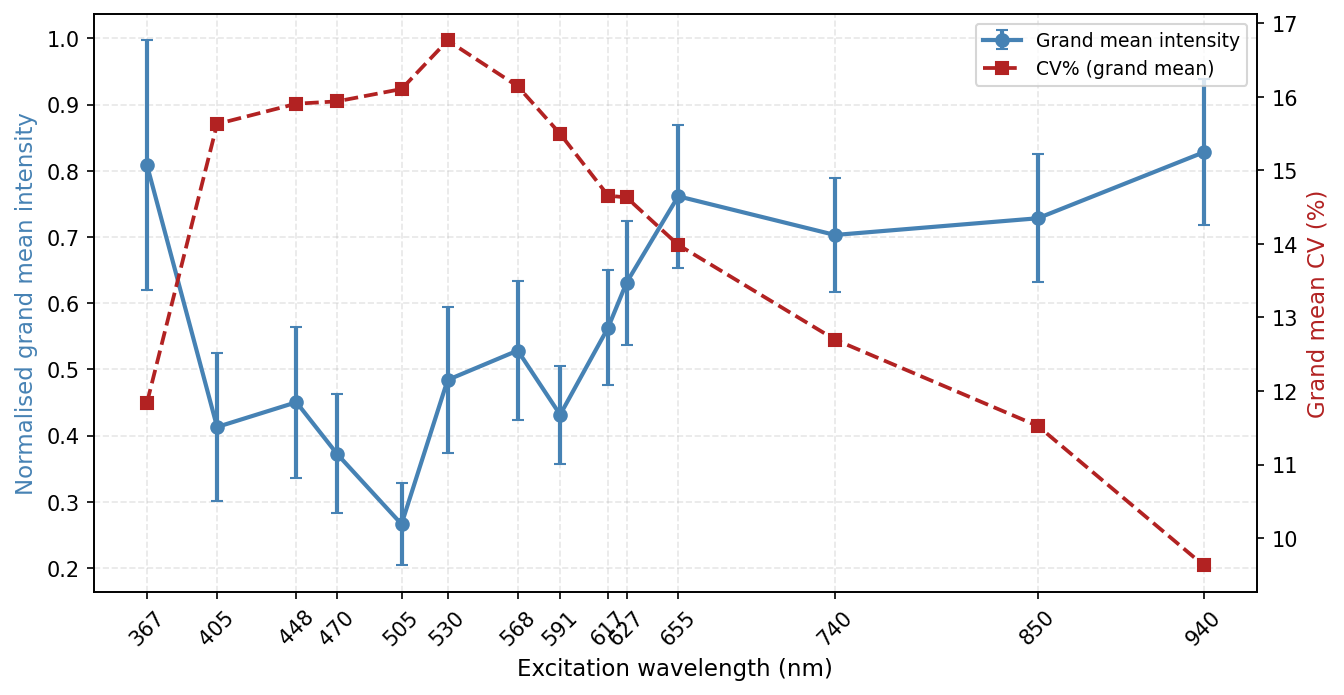

Figure S25. System-level summary of signal magnitude and repeatability. Mean normalized ROI intensity and corresponding coefficient of variation across excitation wavelengths, averaged over all powder samples. Signal intensity and cross-powder variability are lowest near 505 nm and increase toward both spectral extremes. This figure characterizes the excitation-wavelength dependence of two system-level quantities averaged across all ten powders: overall signal magnitude, and intra-sample spatial heterogeneity (texture contrast) of the granular powder surface. Since GLCM/LBP texture features derive from this same spatial heterogeneity, the figure implicitly indicates where in the spectrum texture-based discrimination has the rawest material to work with.

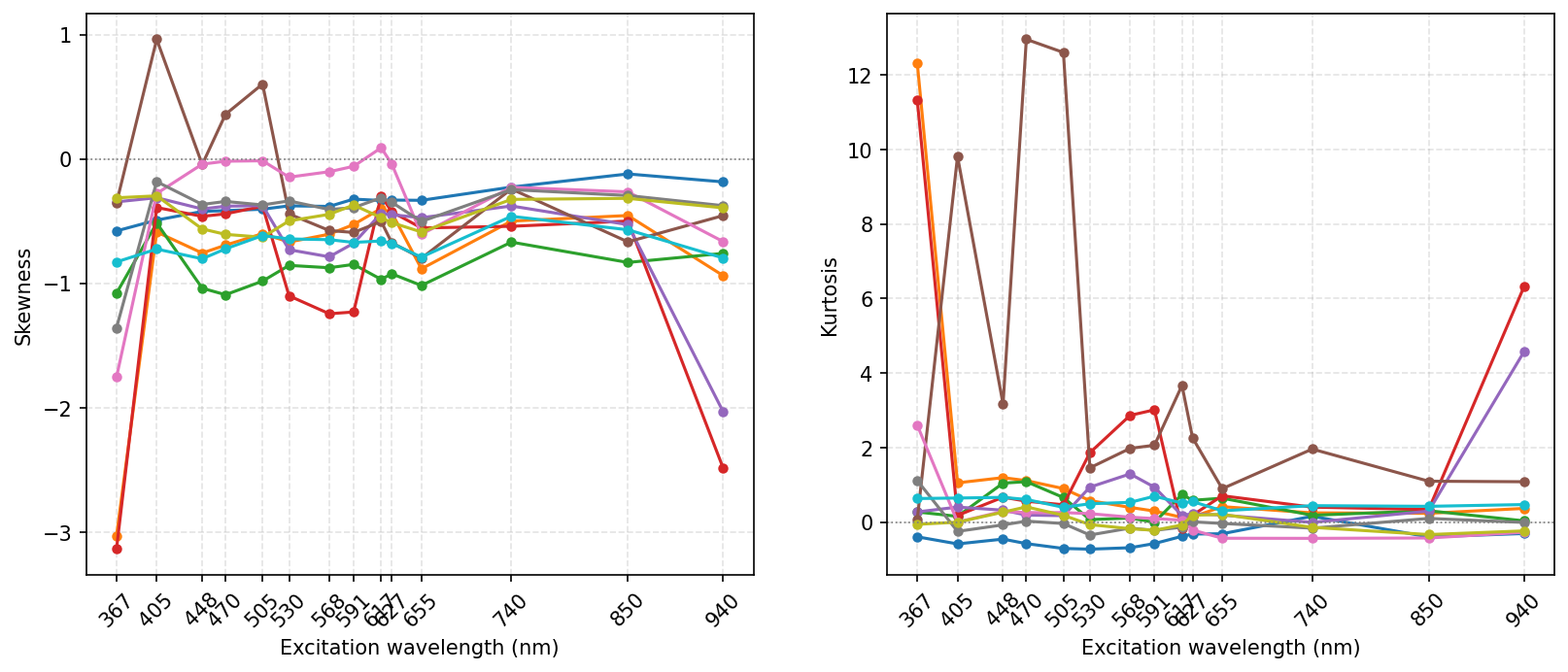

Figure S26. Higher-order intensity statistics of powder images. Skewness and kurtosis of ROI pixel-intensity distributions as a function of excitation wavelength for each powder class. The pronounced negative skewness and elevated kurtosis observed for several powders at 367 nm and 940 nm (e.g., Baking Soda, Aspirin, Ibuprofen, Flour) are consistent with clipping of the intensity distribution at the sensor ceiling rather than an intrinsic higher-order property of the reflected/emitted signal at these wavelengths. Skewness and kurtosis at intermediate wavelengths (405-655 nm), where saturation fractions were negligible, are interpretable as genuine descriptors of intensity-distribution shape

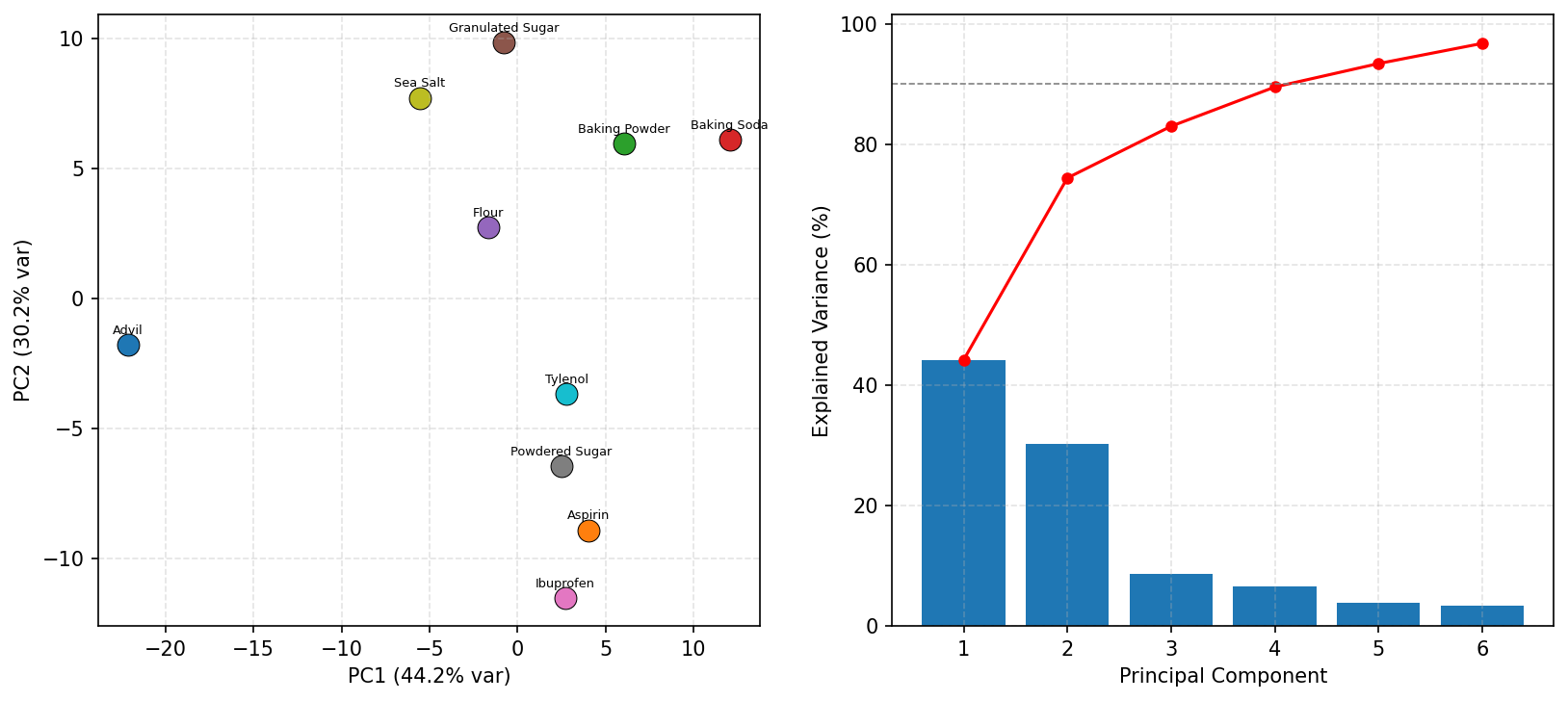

Figure S27. Principal component analysis of multispectral powder features. PCA scores for the concatenated multi-wavelength feature matrix. The first two principal components account for 88.4% of the total variance.

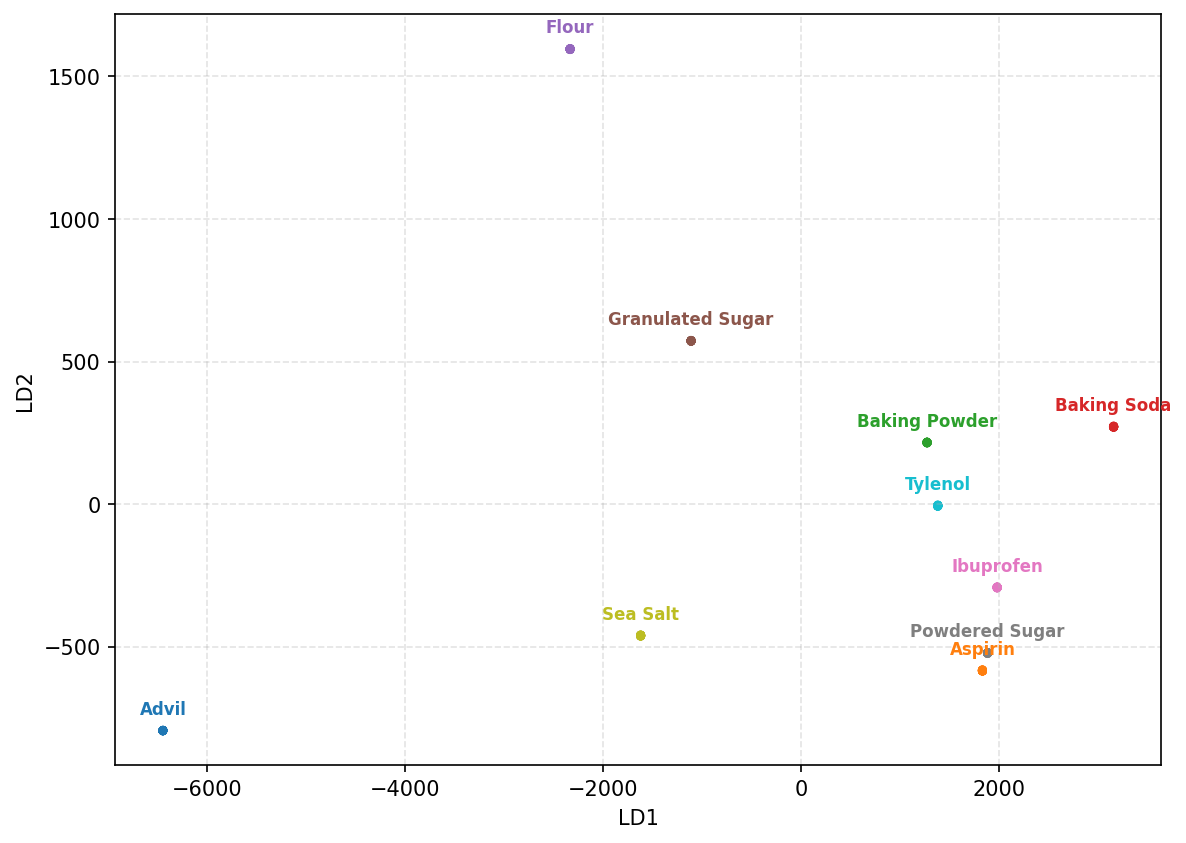

Figure S28. Linear discriminant analysis of multispectral powder features. Two-dimensional projection of the frame-level feature vectors using linear discriminant analysis.

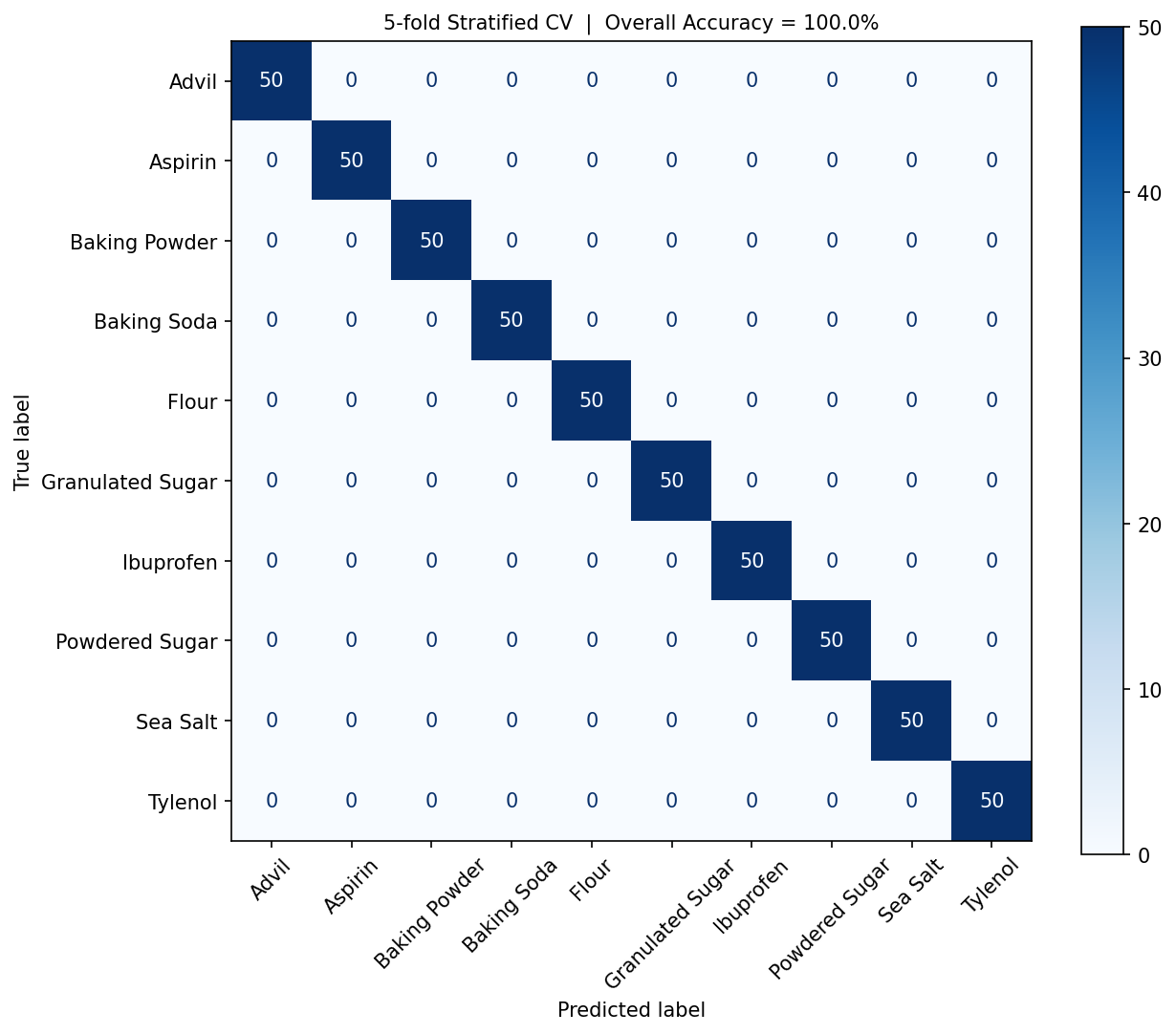

Figure S29. Random Forest classification performance. Confusion matrix obtained from a Random Forest classifier (200 trees) evaluated using five-fold stratified cross-validation. Random Forest classification achieved 100% accuracy under 5-fold stratified cross-validation. Each class was represented by a single physical specimen imaged repeatedly (50 technical replicate frames per well), this result demonstrates that frame-to-frame measurement noise is small relative to between-class spectral/textural differences under fixed acquisition conditions, rather than validating generalization to independent specimens, container placements, or acquisition sessions. Classification performance on independent physical replicates (distinct lots, wells, or imaging sessions) has not yet been evaluated and is identified as a priority for future validation.

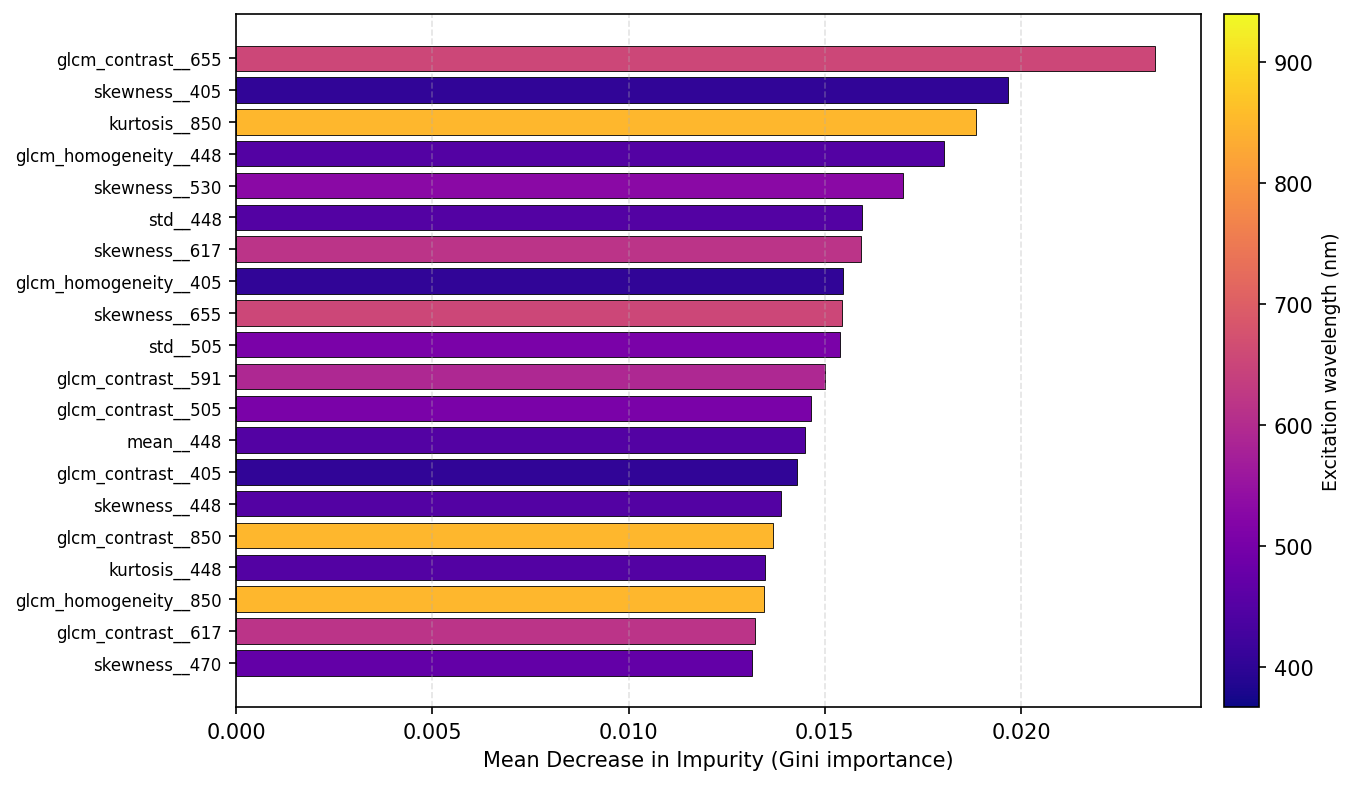

Figure S30. Random Forest feature importance. Relative importance of wavelength-specific photometric and texture features contributing to classification performance.

### Wavelength-Selective Plant Imaging

#### Anthocyanin Stress Monitoring

Whole lettuce (Lactuca sativa) leaves were imaged over seven days under ambient laboratory conditions without supplemental water to induce post-harvest dehydration. Images were acquired six times per day using 530 nm and 627 nm excitation. A 627 nm reference wavelength was used in place of the more commonly reported 665 nm channel because it matches the available hardware configuration. Although absolute AI values differ from conventional implementations, the underlying index formulation is preserved.

Images were normalized using a fixed gray reflectance standard included in each acquisition. Leaf masks were generated from the normalized 530 nm images using thresholding and morphological filtering. The anthocyanin index (AI) was calculated from the normalized 530 and 627 nm images using $AI=\log\left( {I_{norm-627}+\varepsilon}/{I_{norm-530}}+\varepsilon\right)$ where $I_{norm-530}$ and $I_{norm-627}$ denote gray-normalized pixel intensities at the respective wavelengths and $\varepsilon={10}^{-6}$ is a small regularization constant added to prevent logarithmic singularities at zero-valued pixels. The mean AI was calculated over the segmented leaf area. Representative anthocyanin index maps and the corresponding temporal progression are shown in Figures S31 and S32. Mean anthocyanin index increased throughout the monitoring period.

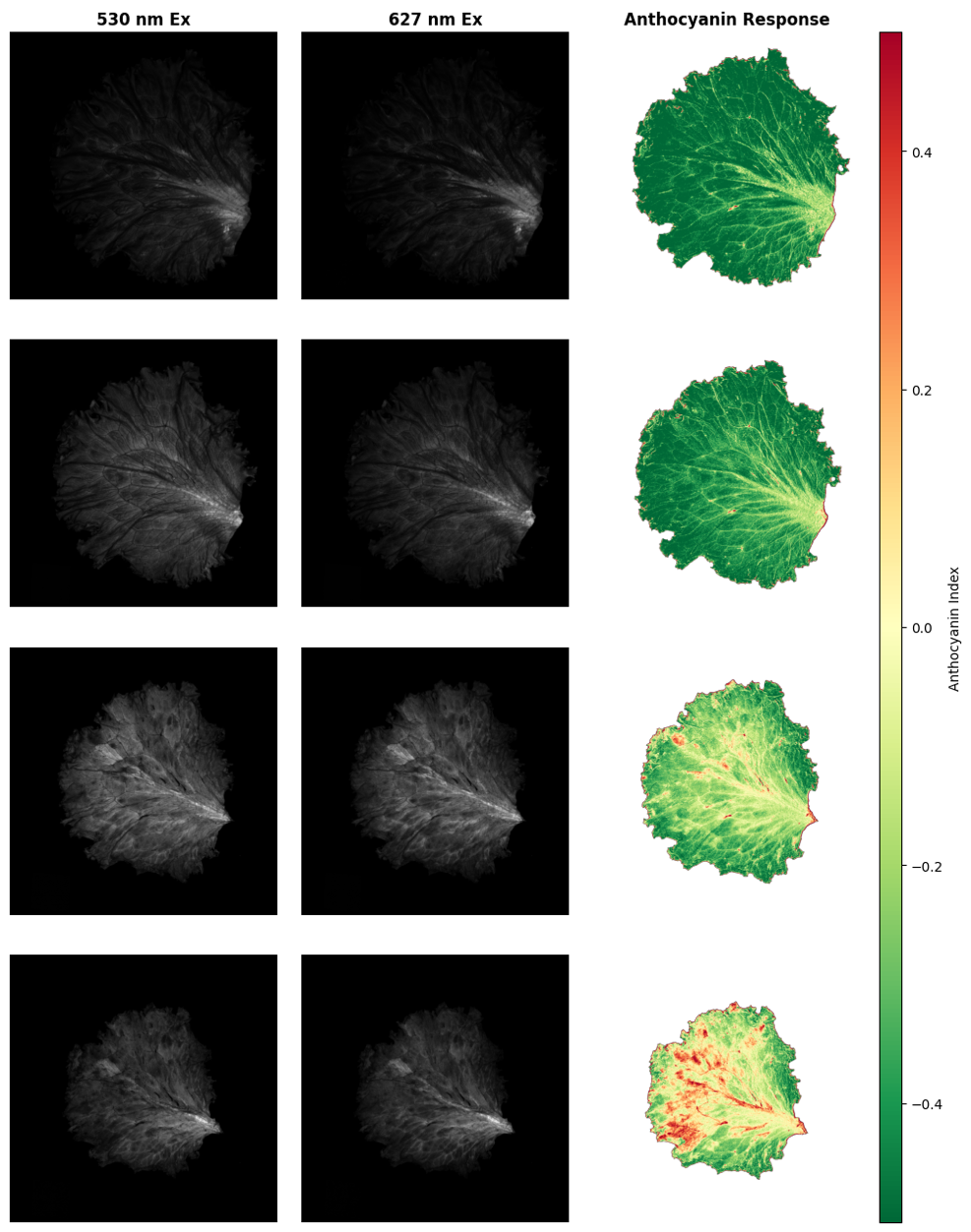

Figure S31. Temporal evolution of the anthocyanin index in lettuce. Representative grayscale images acquired under 530 and 627 nm excitation together with the corresponding anthocyanin index maps for four samples monitored over seven days.

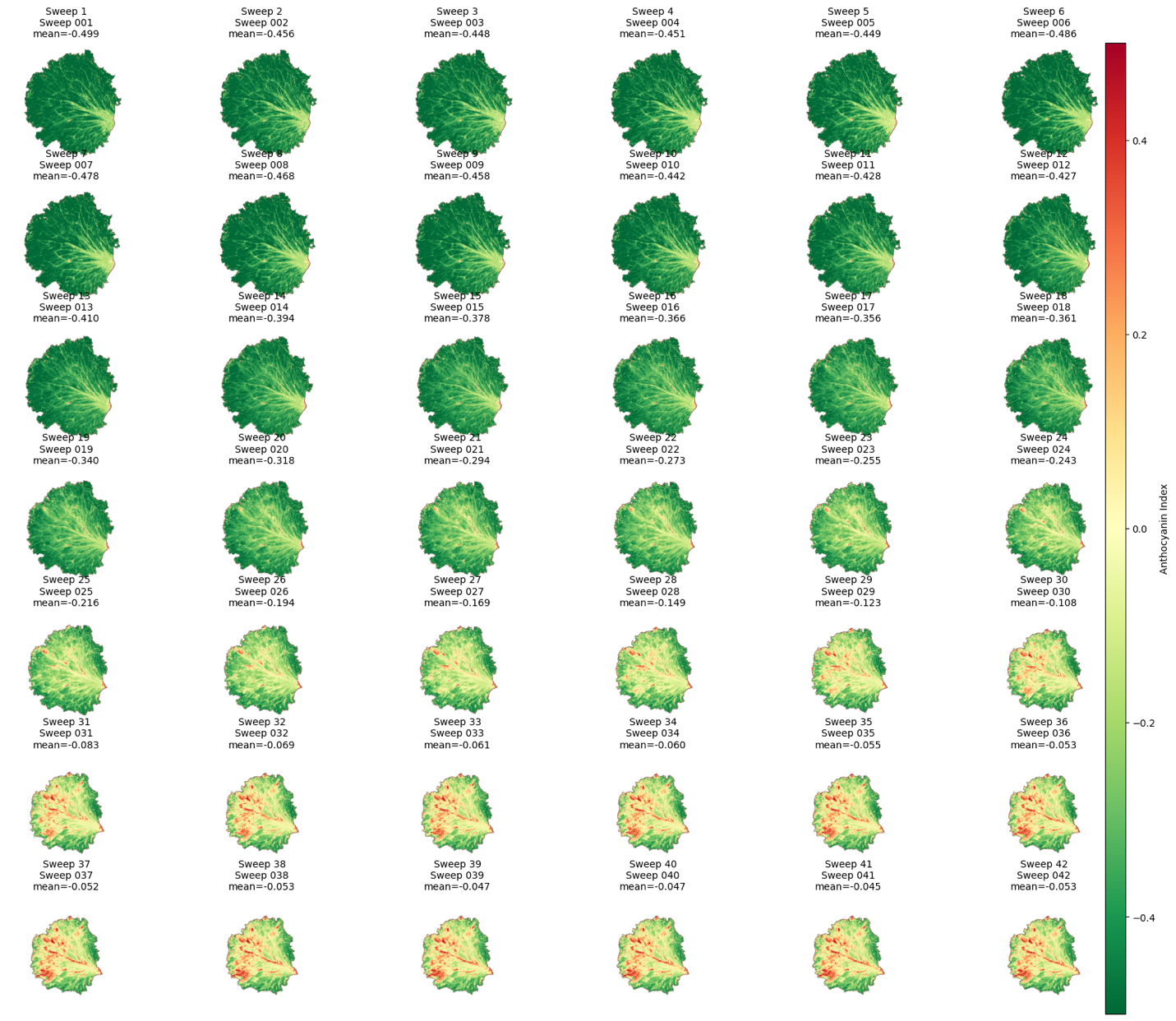

Figure S32. Temporal progression of the anthocyanin index. Anthocyanin index measured over the complete 42-acquisition time series for all samples.

#### Wavelength-Selective Plant-Tissue Imaging

Succulent plants were imaged under ultraviolet (367 and 405 nm) and visible-near-infrared (530, 850, and 940 nm) excitation using a fixed imaging geometry. Pixel-wise ratio images were computed for each wavelength pair after addition of a small numerical offset to prevent division by zero. Cilantro (*Coriandrum sativum*) seedlings were imaged under sequential excitation from 405 to 940 nm. Images were assembled into a multispectral stack, normalized independently by wavelength, and analyzed by principal component analysis (PCA). A false-color composite was generated by assigning the first three principal components to the red, green, and blue display channels. Representative ratio images are shown in Figure S33. Principal component analysis improved visual separation of roots and surrounding soil relative to broadband intensity images.

| 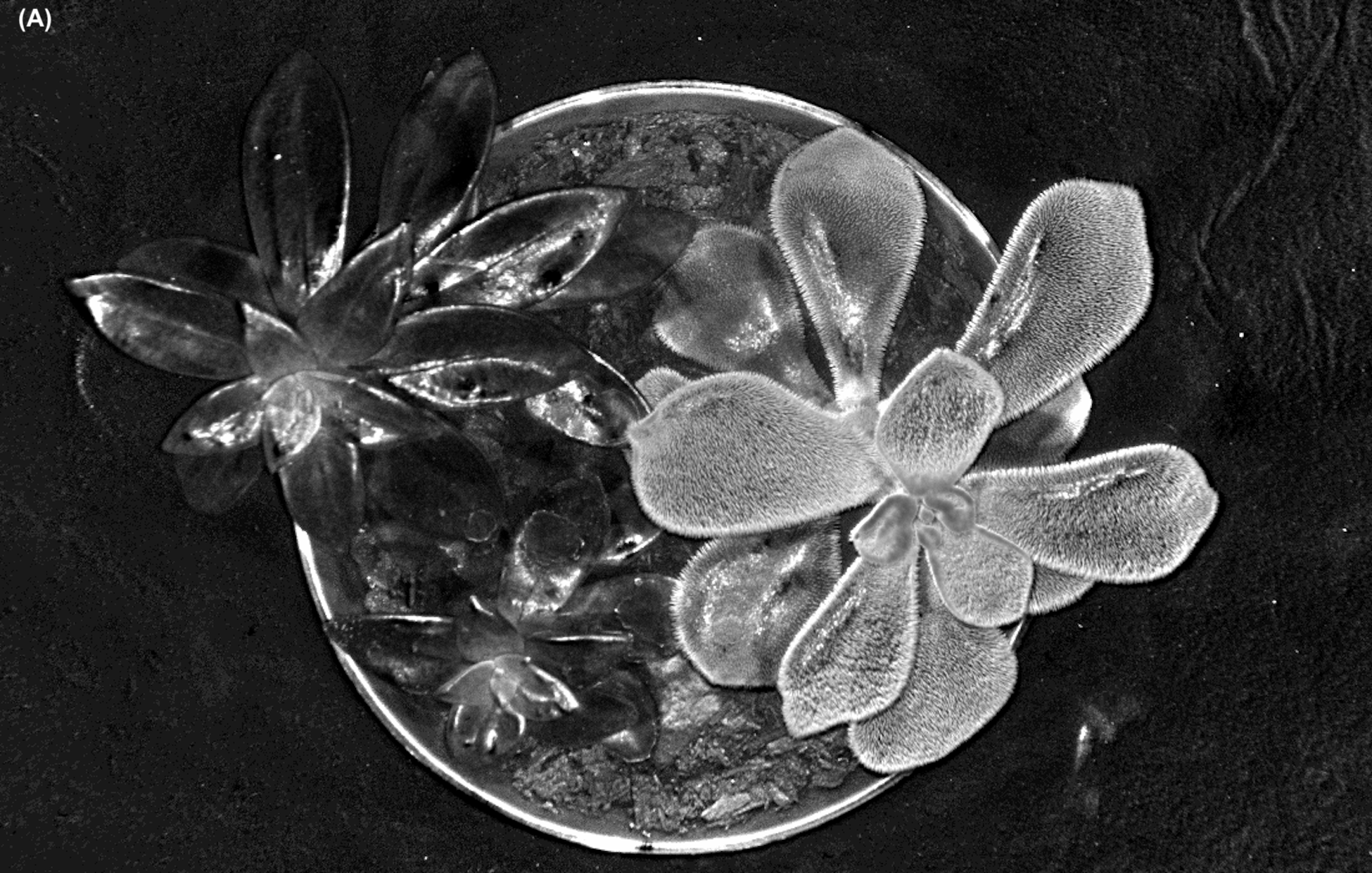 | 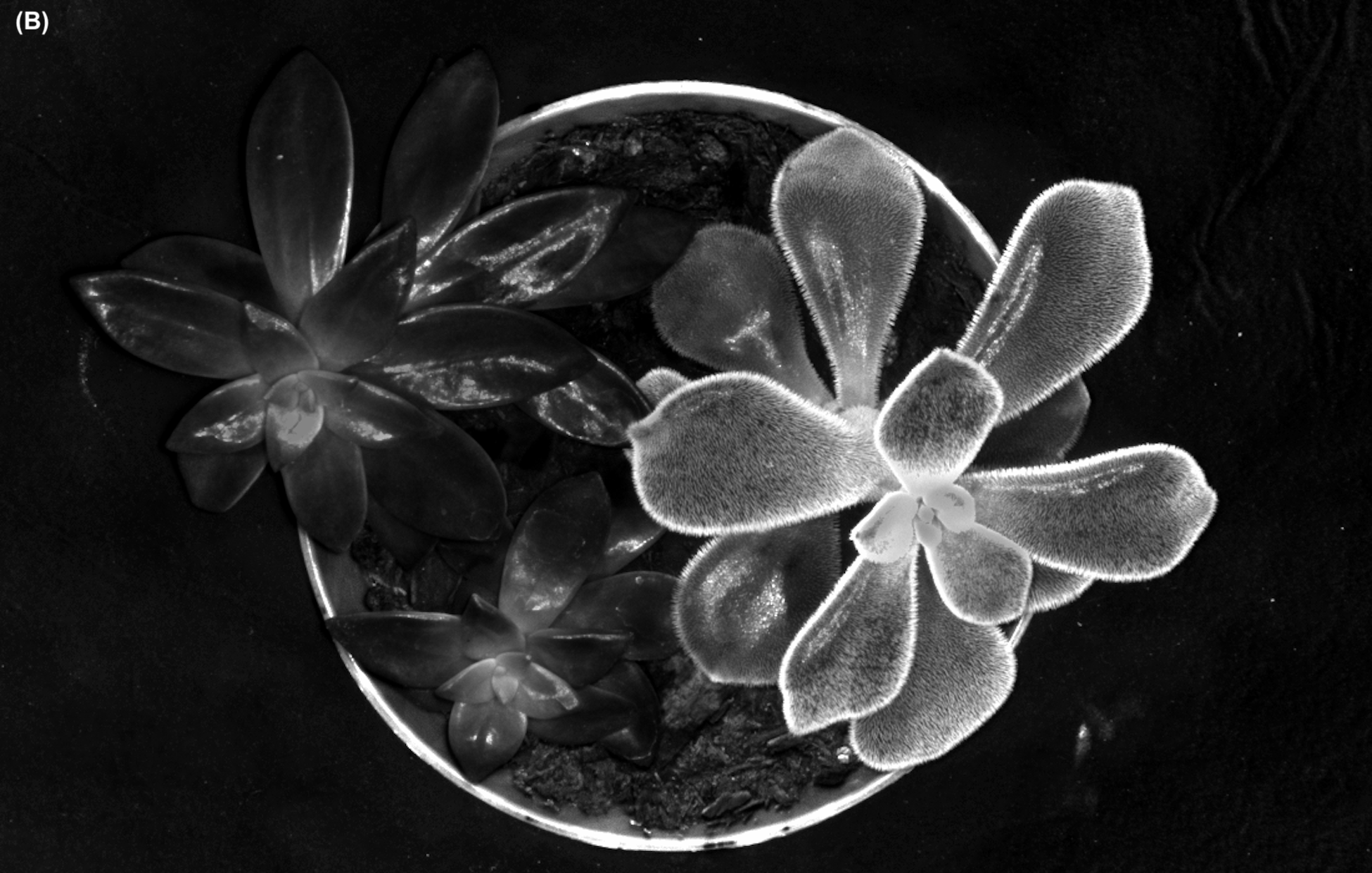 |
| --- | --- |
| 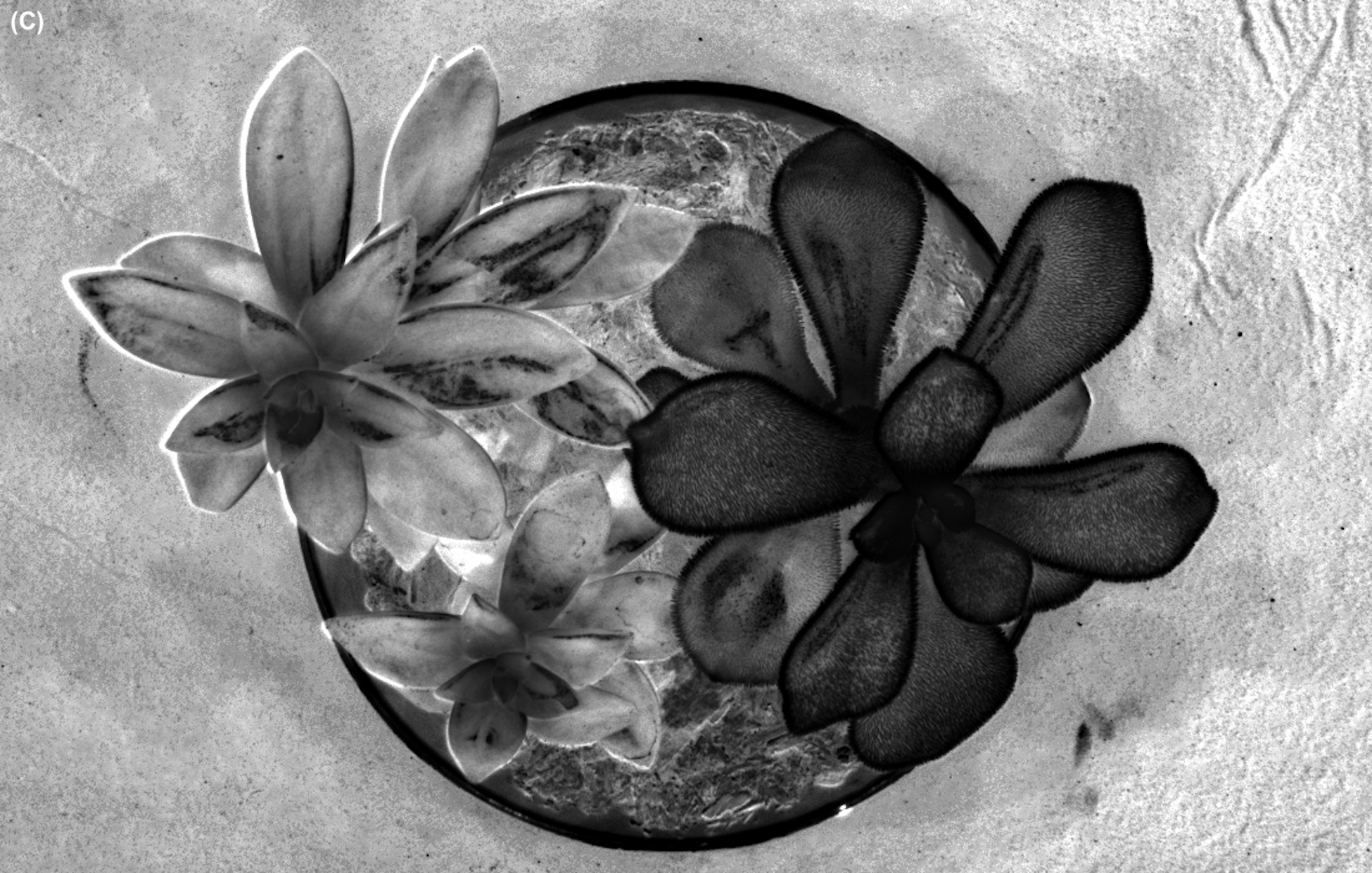 | 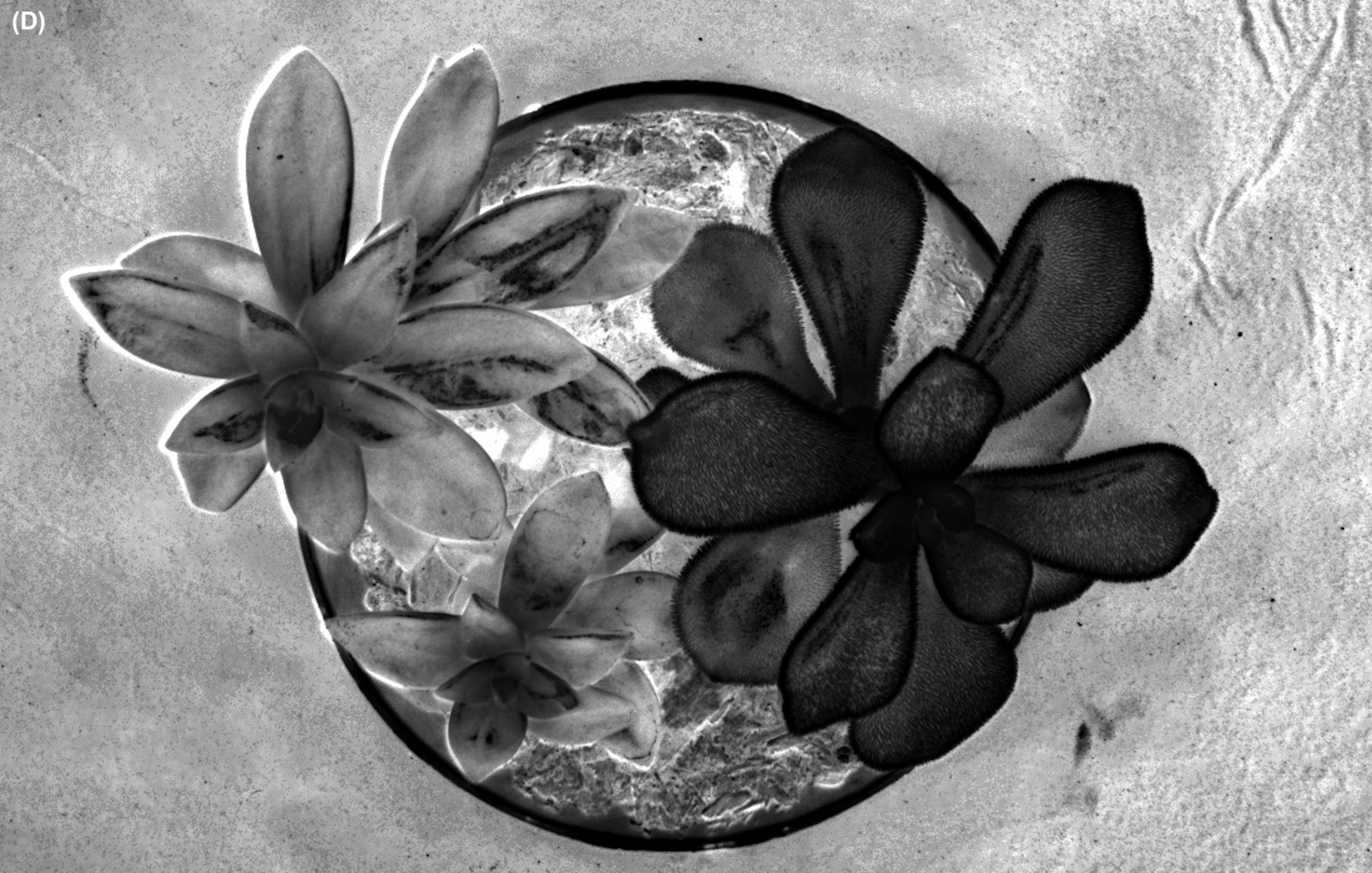 |

Figure S33. Inverted and supplementary excitation-ratio renderings. (A) 405/367 nm, the inverted rendering of the UV excitation ratio shown in Figure 7A; ratio inversion changes image contrast without altering the underlying information content and is provided for perceptual comparison only. (B) 530/940 nm, an additional visible-NIR excitation ratio not shown in Figure 7. As with the 530/850 nm ratio (Figure 7B), optical penetration depth varies substantially with excitation wavelength, and this ratio is expected to sample tissue depth in a manner distinct from the 850 nm pairing; the resulting ratio is interpreted as reflecting wavelength-dependent light transport rather than absolute reflectance. This interpretation is based on established leaf optics literature and the qualitative appearance of the images and was not independently validated here. (C) 850/530 nm, the inverted rendering of the visible-NIR excitation ratio shown in Figure 7B. (D) 940/530 nm, the inverted rendering of panel B. As in panel A, the inversions in panels C and D change image contrast without altering the underlying information content; they are provided for perceptual comparison and enable selection of feature visibility (bright-on-dark vs. dark-on-bright) without additional data acquisition.

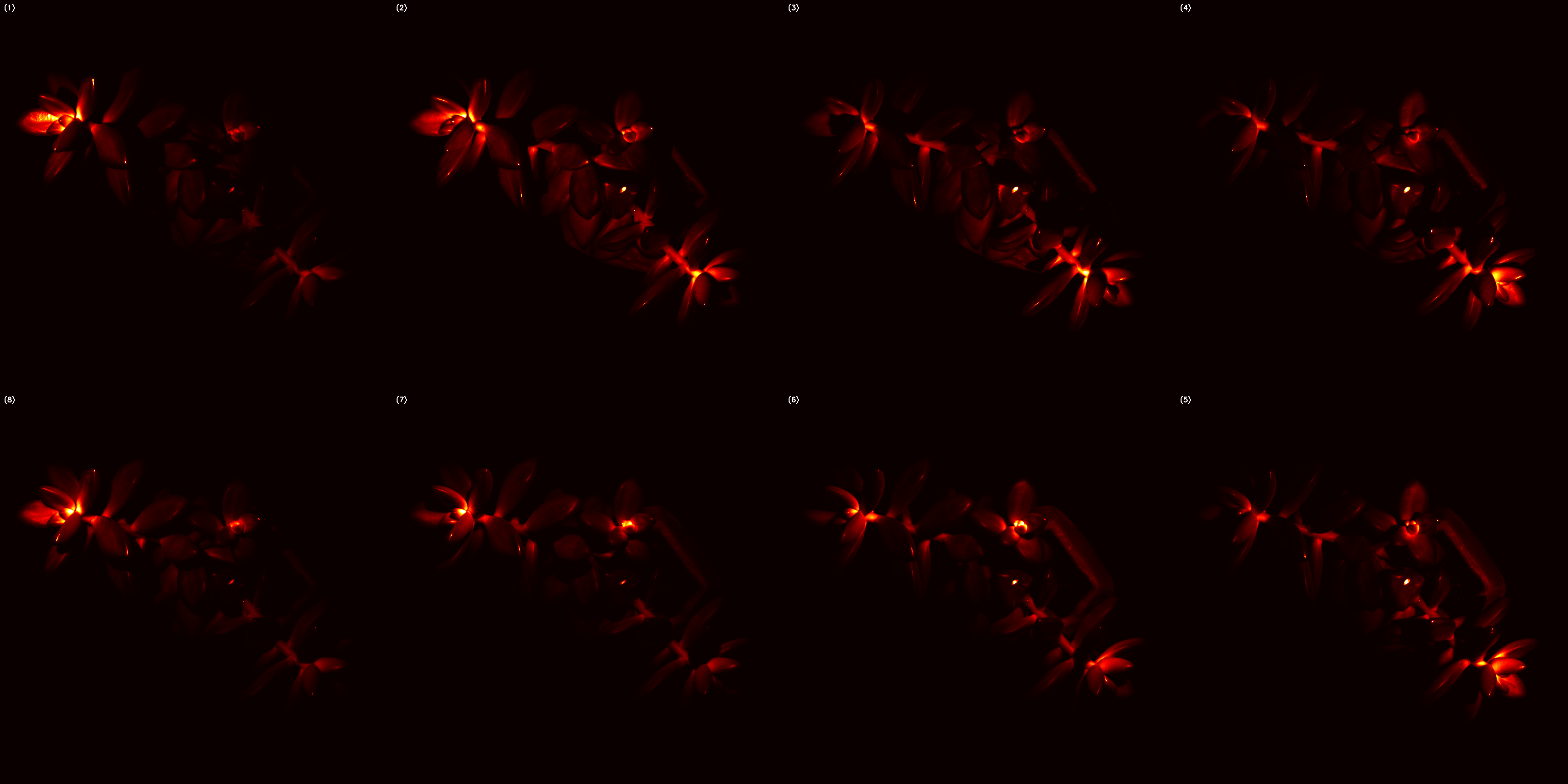

Figure S34. Spatial spectral response of a succulent plant across eight zones at 530 nm excitation. Zones 1-4 (top row) and zones 8-5 (bottom row) display differential signal intensity as a function of illumination direction.

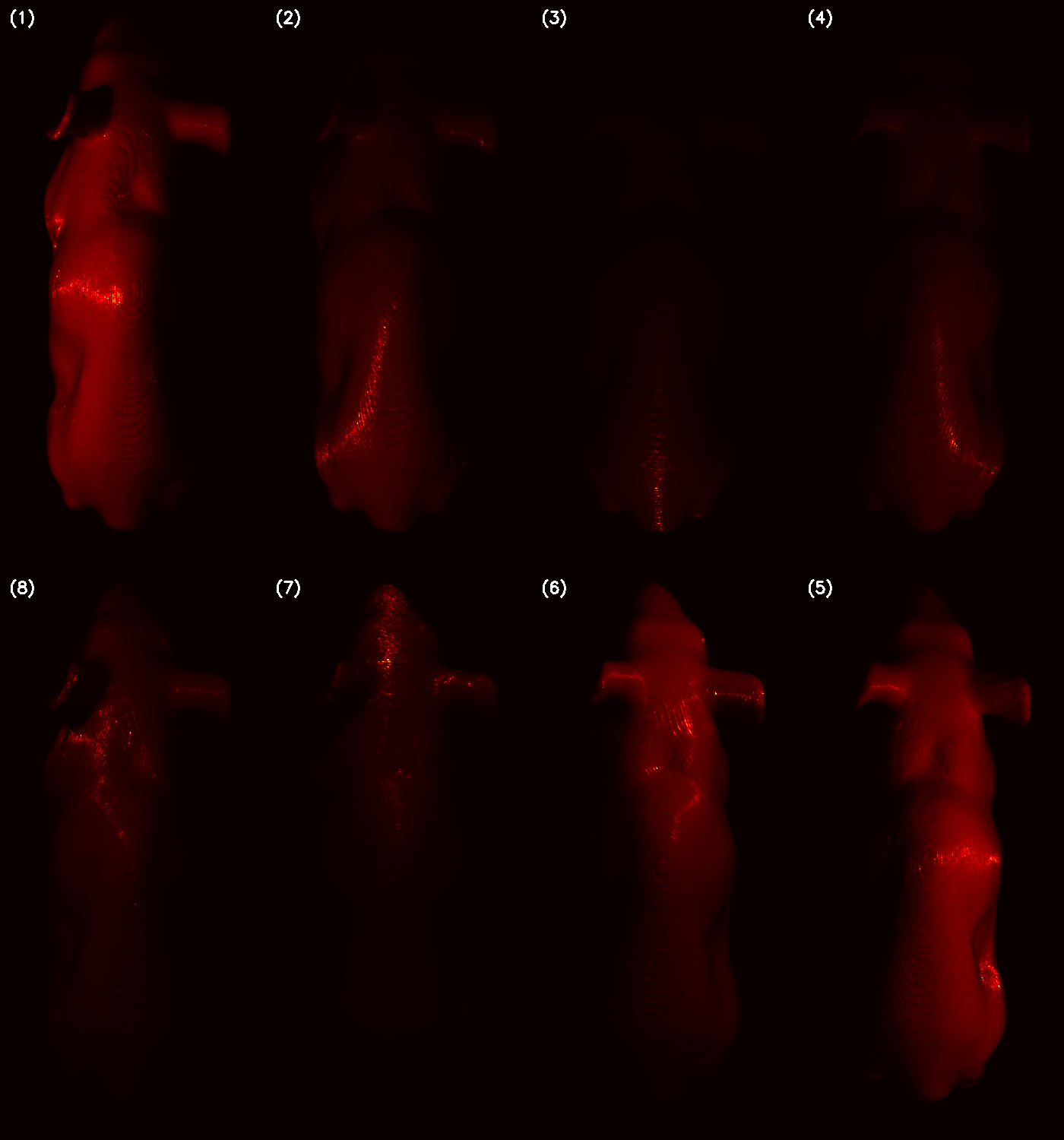

Figure S35. Spatial spectral response of 3D-printed mouse phantom across eight zones in a at 530 nm excitation. Zones 1-4 (top row) and zones 8-5 (bottom row) display differential signal intensity as a function of illumination direction.

### Light Polarization Effects

Glare suppression was evaluated by adding crossed linear polarization to the illumination and detection paths. A linear sheet polarizer was laser-cut to fit over the LED board, with a central aperture for the camera's field of view and mounted on a laser-cut plexiglass frame to keep the film flat and reduce optically active wrinkling. A second linear sheet polarizer served as a manually rotatable analyzer in front of the camera lens. Stray light was mitigated by masking the frame edges with opaque tape and blackening the aperture rim. Images of a curved, transparent sample container were acquired under 448 nm illumination at 100 ms exposure and nominal LED drive levels of 50% and 75%, with the analyzer rotated to dial positions of 0°, 45°, 90°, and 135°, corresponding to inter-polarizer angles Δθ of 90° (crossed), 45°, 0° (aligned), and 45°, respectively. Glare is reported as the saturated-pixel count Nₛₐₜ (pixels with digital number ≥ 60,000), a relative proxy for saturated image area. This metric is governed by the sensor and is intended for within-session relative comparison rather than absolute radiometric quantification.

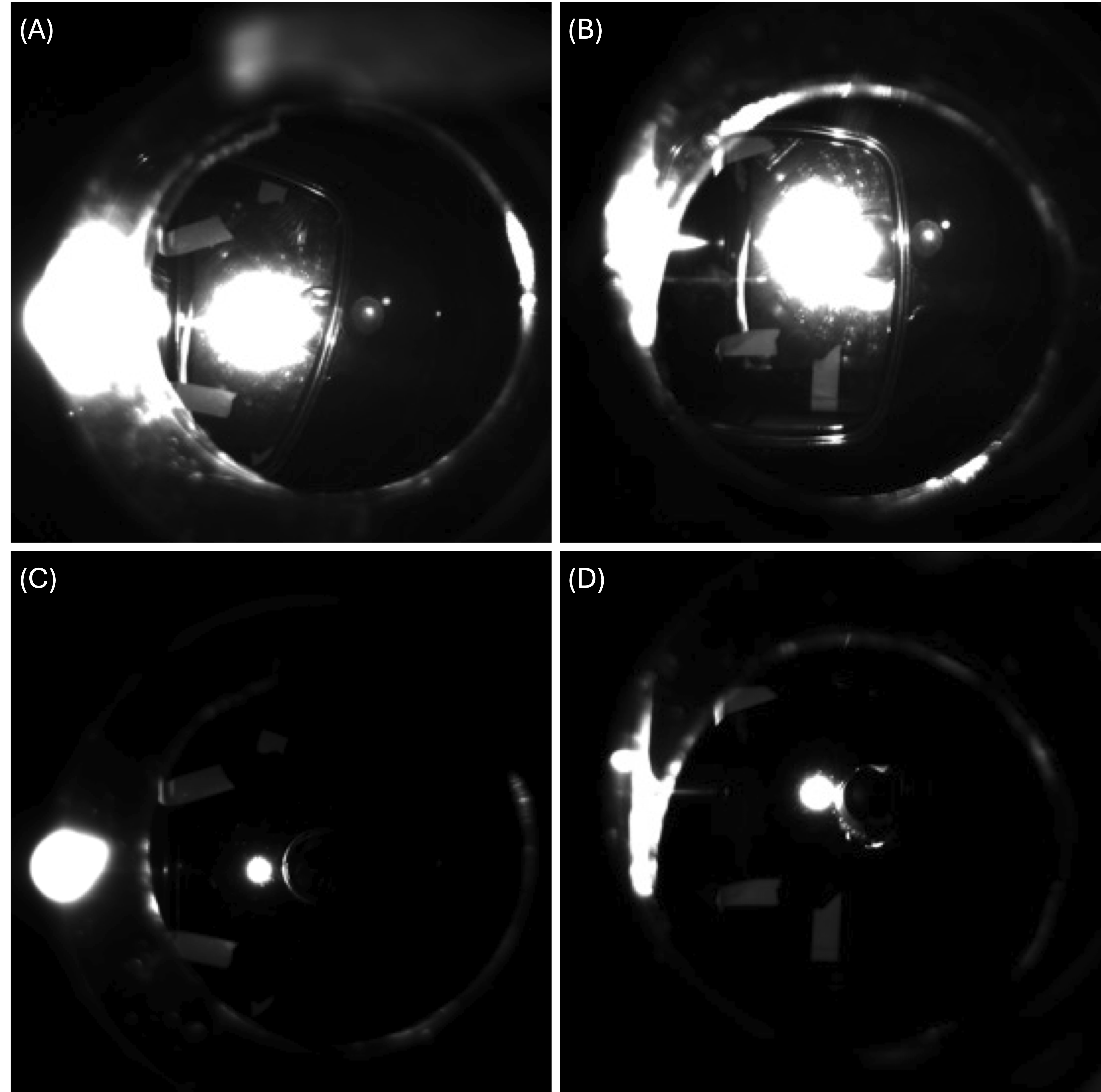

Figure S36. Cross-polarization suppression of specular glare from a curved biological sample container. Δθ denotes the angle between the transmission axes of the two polarizers (Δθ = 0°, aligned; Δθ = 90°, crossed). Top row (A, B): aligned polarizers (Δθ = 0°) at 50% and 75% nominal LED drive, showing extensive specular glare and a large saturated region over the container. Bottom row (C, D): crossed polarizers (Δθ = 90°) at the same drive levels, showing substantially reduced glare with only residual highlights. Crossing the polarizers reduced Nₛₐₜ by approximately 78-84% at 50% drive and approximately 51% at 75% drive relative to the aligned condition.
